# Localized APP pathology in the hippocampus is sufficient to result in progressive disorganization of the timing of neuronal firing patterns

**DOI:** 10.1101/2022.10.24.513188

**Authors:** Silvia Viana da Silva, Matthias G. Haberl, Kshitij Gaur, Rina Patel, Gautam Narayan, Max Ledakis, Maylin L. Fu, Edward H. Koo, Jill K. Leutgeb, Stefan Leutgeb

**Affiliations:** Neurobiology Department, School of Biological Sciences, San Diego, La Jolla, CA, USA; NeuroCure Excellence Cluster and German Center for Neurodegenerative Diseases (DZNE), Berlin, Germany; Neuroscience Research Center, Charité Universitätsmedizin Berlin, Germany; School of Medicine, Department of Neurosciences, University of California, San Diego, La Jolla, CA, USA; Kavli Institute for Brain and Mind, University of California, San Diego, La Jolla, CA, USA

## Abstract

Deficits in spatial navigation are among the early symptoms in Alzheimer’s disease patients, consistent with the hippocampal formation as the site for spatial computations and disease onset. Although the correspondence between the early symptoms and brain regions that are affected early in the disease has been recognized, it is not clear whether progressive cognitive decline is solely caused by a spreading pathology or whether a focal pathology can by itself cause aberrant neuronal activity in a larger network. These possibilities cannot be distinguished in standard disease models which broadly express APP across brain regions. We therefore generated a mouse model in which the expression of mutant human APP was limited to hippocampal CA3 cells (CA3-APP mice). We first asked whether the limited pathology in CA3 can result in memory deficits and found impaired performance of CA3-APP mice in a hippocampus-dependent memory task. By then recording in the CA1 region, we asked to what extent neuronal activity patterns emerged in a brain region which received projections from APP-expressing CA3 cells, but did itself not show any primary pathology. While the spatial firing patterns of CA1 cells were preserved, we observed a reduced theta oscillation frequency in the local field potential and in a subpopulation of principal cells in CA1. Furthermore, CA1 interneurons showed decreased theta oscillation frequencies, and this effect was even more pronounced in CA3 interneurons, which also do not directly express APP. Pathology that is highly localized and limited to presynaptic cells is thus sufficient to cause aberrant firing patterns in postsynaptic neuronal networks, which indicates that disease progression is not only from a spreading molecular pathology but also mediated by progressive physiological dysfunction.

## Introduction

Alzheimer’s disease (AD) is a neurodegenerative disorder characterized by progressive cognitive deficits. In particular, spatial navigation is impaired early in human Alzheimer’s disease patients and has emerged as a potential cognitive biomarker to detect AD at preclinical stages^1,2^. Importantly, the early deficits in spatial navigation are not only relevant as a diagnostic strategy, but also commensurate with the finding that brain circuits that support navigation, such as the hippocampus and entorhinal cortex^3^ are particularly impaired at early disease. Aberrant firing patterns have in turn been shown to accelerate disease progression^4^, and restoring dysfunction in these regions is therefore a particularly urgent target for treatment approaches.

Expressing proteins with mutations found in familial forms of Alzheimer’s disease causes wide ranging synaptic and spatial memory deficits in mice, yet it is not well understood how molecular and synaptic deficits change the network to cause memory deficits, particularly at early phases of the pathology. The majority of studies that address network dysfunction have been done in models of AD in which synapse loss^5^, deposition of amyloid-beta plaques^6–8^, or the emergence of neurofibrillary tangles is advanced^9,10^. Rodent models at advanced stages of amyloid or tau deposition display behavioral deficits associated with deteriorated spatial properties of place cells in the hippocampus^6,7,10,11^ and grid cell in the entorhinal cortex^7,12,13^, but there is generally no causal link as spatial coding deficits can emerge at a different time in disease progression than behavioral deficits.

In addition to the outstanding question of which emerging circuit dysfunction leads to the initial memory deficits in early disease stages (i.e., before extensive plaque formation and cell death), it is also not known whether broadening circuit dysfunction emerges predominantly from the spreading molecular pathology across brain regions or also from connectivity with brain areas with the initial APP pathology. These possibilities cannot be distinguished with models of panneuronal APP expression, which show an initial pathology in few cortical areas that then increasingly broadens across brain regions. Even in disease models that initially restrict APP or Tau expression to entorhinal cortex, progressive broadening of pathology to other brain regions occurs because the expression of the transgene spreads to other brain regions. To distinguish between pathology that emerges from spreading APP expression as opposed to physiological responses to prolonged local pathology, we used transgenic mice in which the expression of mutant human amyloid precursor protein (APP) remains selective for hippocampal CA3 pyramidal cells throughout the lifespan (CA3-APP mice). The CA3-APP mouse model shows no plaques or cell death in the hippocampus and thus constitutes a model to investigate the earliest features of AD related dysfunction^14^.

Using the CA3-APP mice allowed us to determine whether local AD synaptic and cellular deficits eventually lead to dysfunction in non-APP expressing neuron populations that are critical for supporting spatial navigation. By testing the mice at two different age points we were able to determine that the localized pathology resulted in dysfunctional network oscillations in a broader network, including in the hippocampal CA1 area that receives projections from CA3, but does itself not express APP. The aberrant oscillations were accompanied by memory deficits and by altered temporal firing patterns of individual principal cells and of interneurons, but the spatial firing pattern remained largely intact. This shows that altered network function can spread to areas that are themselves not the origin of Alzheimer’s disease pathology and indicates that disease progression can be driven by spreading circuit dysfunction in addition to pathological changes at the level of individual cells. These findings also further support the idea that memory dysfunction in Alzheimer’s disease can be improved by restoring brain oscillations.

## Results

To test the effects of localized pathology on behavioral and circuit dysfunction, we generated mice that selectively expressed mutant hAPP in CA3 principal cells. The selective expression of the transgene in only hippocampal CA3 cells allowed for the examination of pathology in the hippocampal CA1 region, which receives direct projections from CA3 but does itself not express hAPP. The targeting of hAPP was achieved by crossing three transgenic lines (**Figure 1A**). The Grik4-Cre^15^ transgene provided the selectivity for CA3 principal cells, while the combination of Cre-dependent tTA^16^ and tetO-APP transgenes^17^ allowed for cell-type specific expression of hAPP carrying the Swedish and Indiana mutations. We used offspring that was positive for the three transgenes (CA3-APP mice) as well as control littermates which did not carry the tetO-APP transgene (see **Supplementary Table 1** and methods for more details). We investigated the mice at two different time points of the amyloid pathology progression, at 4-6 months (young adults) and at 16-18 months (aged) (Figure 1B). We observed that the transgene was expressed in 74.5 ± 2.2% of CA3 principal cells in aged adult mice (**Supplementary Figure 1A**), and previously showed that the transgene was expressed in close to 60% of CA3 cells by 6 months^14^. No plaques were observed in our mice at any age point, and in line with previous observations, no APP expression was seen outside of the CA3 region in brain sections from young adult and aged mice^14^.

**Figure 1.**
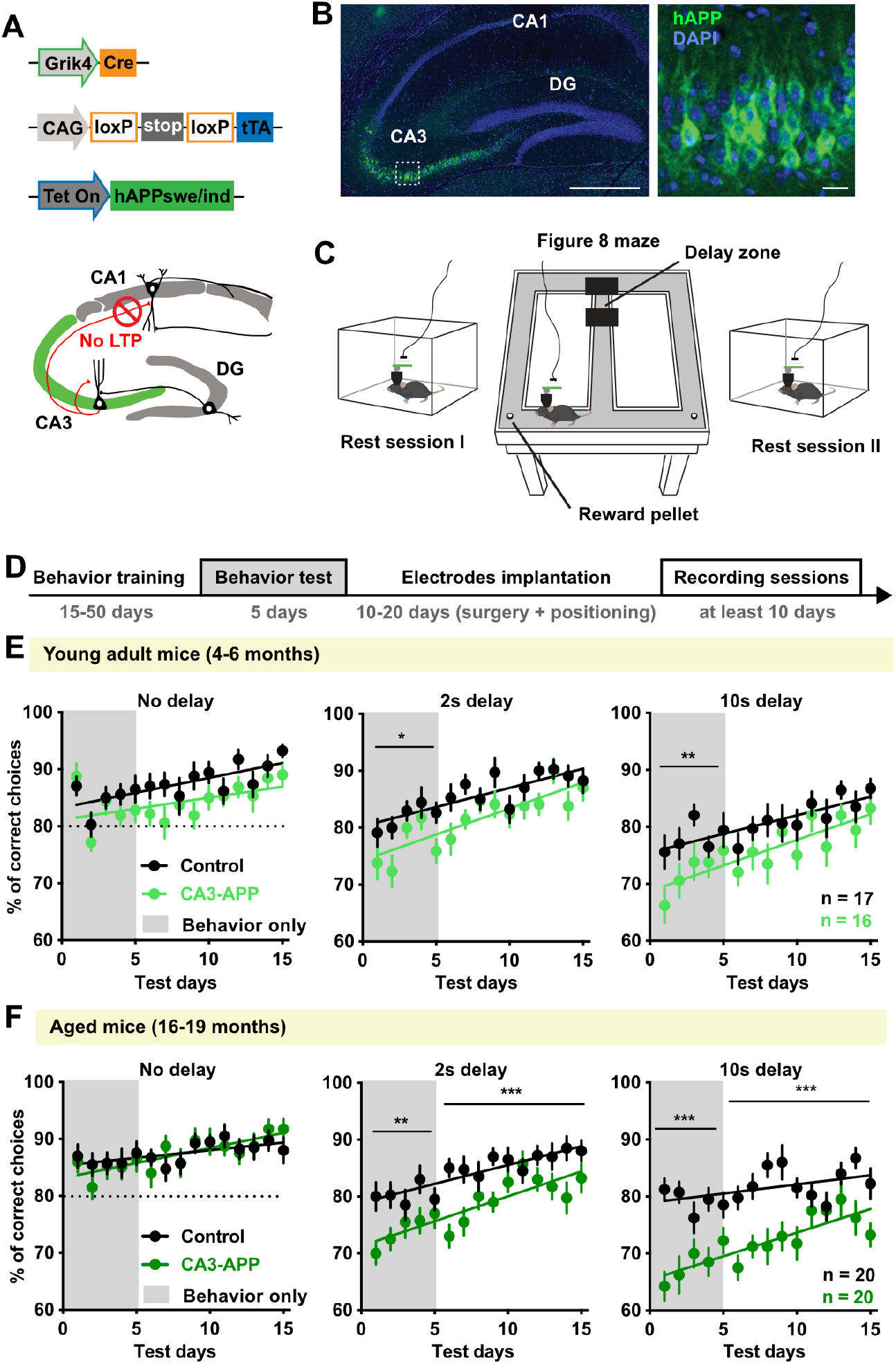
Hippocampus-dependent memory was impaired in CA3-APP mice. **A.** CA3-APP mice carried three transgenes: (i) Cre-recombinase under the regulation of the Grik4 promoter for CA3 specificity, (ii) a tetracycline-controlled transactivator protein (tTA) transgene under the control of Cre-LoxP recombination and (iii) the APP mutant gene under the control of a tetracycline-responsive promoter element (TRE). LTP of Schaffer-collateral inputs to CA1 is impaired in CA3-APP mice starting at 4 months of age (Vicario-Orri et al. 2020)^14^. **B.** Example immunostaining of sections from aged CA3-APP mice. Anti-hAPP (6E10 antibody) staining in green, DAPI in blue. Scale bars are 0.5 mm (left) and 20 μm (right). **C.** Recording paradigm included 30 laps in the figure-8 maze preceded and followed by a 10-15 min rest session in the home cage. **D.** Experimental timeline. **E.** Behavioral performance over experimental days for young adult CA3-APP mice (n = 16) and littermate controls (n = 17). During the initial 5 days of behavior testing in the delayed versions of the task, there was a reduction in % correct choices by CA3-APP mice in comparison to aged-matched controls (2-way RM ANOVA, p = 0.020 for 2-s delay and p = 0.0007 for 10-s delay, Sidak’s post-hoc test). This difference was not observed on subsequent recording days (2-way RM ANOVA, p < 0.0001 for 2-s delay and p = 0.011 for 10-s delay, Sidak’s post-hoc test; see supplementary **Figure 2C, D**). The performance of the mice improved over time in each of the delay conditions (see linear regression analysis in **Supplementary Table 2**) for both experimental groups. **F.** Behavioral performance over experimental days for aged CA3-APP mice (n = 20) and littermate controls (n = 20). In the delayed versions of the task, the percentage of correct choices was reduced in CA3-APP mice in comparison to aged-matched controls, both during the initial 5 days of testing (2-way RM ANOVA, p = 0.001 for 2-s delay and p < 0.0001 for 10-s delay, Sidak’s post-hoc test) and during the recording period (2-way RM ANOVA, p = 0.0008 for 2-s delay and p < 0.0001 for 10-s delay, Sidak’s post-hoc test; see supplementary **Figure 2E, F**). The performance of all experimental mice improved over time for all delay conditions (see linear regression analysis in **Supplementary Table 2**). * p < 0.05, ** p < 0.01, *** p < 0.001.

### CA3-APP mice displayed deficits in hippocampus-dependent spatial navigation

To test whether localized APP expression in the hippocampal CA3 area was sufficient to cause a memory deficit, we trained mice to perform a spatial alternation task in a figure-8 maze (**Figure 1C**). In this task, animals needed to remember where they came from in order to make a correct turn during the next choice. This task is hippocampus-dependent when a brief delay is inserted in the central arm^18^, while the version without delays is not hippocampus-dependent, thus controlling for general motor deficits. We first tested mice for 5 days before any surgical manipulation to ensure that differences were not arising from inflammatory processes related to the surgery^19^. We observed that CA3-APP mice performed at control levels in the continuous hippocampus-independent version of the task (p = 0.65 for young adult mice and p = 0.83 for aged mice, 2-way ANOVA with Sidak’s post-hoc test, **Figure 1D, E**). In the hippocampus-dependent version with a delay of 2 s or 10 s, the performance of CA3-APP mice was impaired. The impairment already emerged in young adult CA3-APP mice (p = 0.020 for 2-s delay and p = 0.0007 for 10-s delay, Sidak’s post-hoc test) but was more pronounced in the aged group (p = 0.001 for 2-s delay and p < 0.0001 for 10-s delay, Sidak’s post-hoc test; **Supplementary Figure 1D, E**).

### Activity patterns of CA1 principal cells and interneurons were altered by APP expression in CA3 principal cells

We next aimed to understand whether the neuronal activity in the behavioral task was altered by APP expression in CA3, and if so, how pathological activity might be correlated with the memory impairments. In particular, we investigated whether APP expression resulted in a network dysfunction in cell populations that are postsynaptic to the cells with amyloid pathology. We therefore recorded from the hippocampal CA1 area, which is the main target of CA3 axons (**Supplementary Figure 2A, B**). The recordings were performed during an additional 10 days of testing in the spatial alternation task (**Supplementary Figure 1B**) and were from putative principal cells and putative interneurons (see **Supplementary Figure 2C** for classification) and also included the local field potential (LFP). Neuronal activity was also recorded during short rest periods before and after the behavioral testing for assessing cell stability over time and sparsity of cells, as some cells are silent during running. Only cells that were stable throughout the entire recording session were used for further analysis, and firing rate and spike shape were used to classify recorded spikes as putative principal cells or putative interneurons.

During behavioral sessions while recording, young adult CA3-APP performed at similar levels as control littermates (2-s delay: p = 0.083; 10-s delay: p = 0.058, 2-way ANOVA with Sidak’s post-hoc test) while aged CA3-APP mice continued to display increased erroneous choices with both the 2-s delay (p = 0.0008, Sidak’s post-hoc test) and the 10-s delay (p < 0.0001, Sidak’s post-hoc test, **Supplementary Figure 1C-F**). Control as well as transgenic mice showed an improvement in their performance across experimental days in both age groups (all slopes are different from zero, **Figure 1E and F**, see also **Supplementary Table 2**). In summary, young adult CA3-APP mice thus displayed only a transient memory deficit, while aged CA3-APP mice showed a robust impairment in spatial memory.

We next analyzed the firing properties of CA1 principal cells and began with the average firing rate throughout the entire recording session, which includes rest periods and running in the maze in all three delay conditions. Young adult CA3-APP mice showed a small reduction in the average firing rate of putative CA1 principal cells (control 1.6 ± 0.1 Hz, CA3-APP 1.4 ± 0.1 Hz, p = 0.0048, Mann-Whitney (MW) test, **Figure 2A, B**), and the reduction was more pronounced in aged CA3-APP mice (control 1.3 ± 0.06 Hz, CA3-APP 1.0 ± 0.04 Hz, p = 0.0006, MW test, **Figure 2B**). Despite these rate differences, the percentage of active cells in the maze (firing rate > 0.1Hz) was similar between genotypes in young adult (control 90.7 ± 2.6%, CA3-APP 90.7 ± 3.2%, p = 0.997, two-sample t-test) and aged mice (control 81.4 ± 1.7%, CA3-APP 77.5 ± 6.3%, p >0.999, MW test, **Figure 2C**). The average velocity of control and CA3-APP mice was similar in both young adult and aged groups when running in the maze (**Supplementary Figure 3A, B**), but we found a small reduction in the velocity of young adult CA3-APP mice during the rest periods (**Supplementary Figure 3A**). Since velocity and brain state can affect the firing rate of neurons, we therefore repeated the analysis of firing rate with only data while running in the maze when no differences in velocity were found. We again observed reduced firing rates in young and aged CA3-APP mice compared to age-matched controls (**Supplementary Figure 3C-D** and **Table 3**). These results confirm that the reduction in firing rates is independent of behavior.

**Figure 2.**
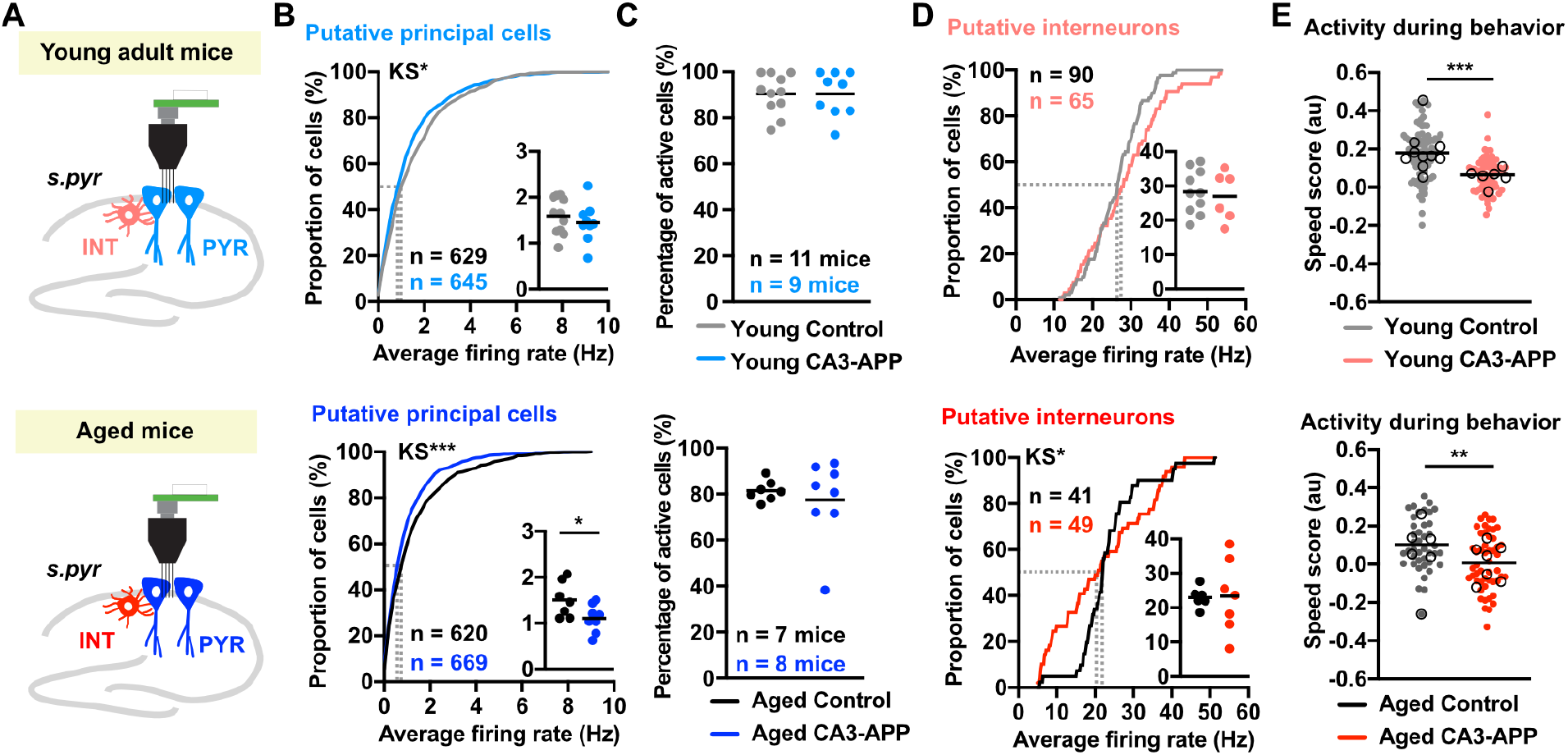
Firing rates of CA1 principal cells were largely reduced in aged CA3-APP mice. **A.** Classification of single units as putative principal cells or interneurons was done based on their firing rates and action potential shape (see supplementary **Figure 2B**). Young adult mice, upper panel; aged mice, lower panel. **B.** Cumulative distribution of average firing rate of all putative CA1 principal cells during the entire recording session (rest and behavior). The inset shows the same data per mouse (young adult: p = 0.45; aged: p = 0.041, two-sample t-test). CA1 principal cells of CA3-APP mice have reduced firing rates compared to principal cells of aged-matched control mice (young adult: p = 0.0048; aged: p = 0.0006, MW test). **C.** Percentage of cells active during the figure-8 task did not differ between CA3-APP mice and age-matched controls (young adult control, n = 11; CA3-APP, n = 9; aged control, n = 7; CA3-APP, n = 8 mice). **D.** Cumulative distribution of average firing rate of putative interneurons during the entire recording session. The average firing rates of CA1 interneurons did not differ between CA3-APP mice and age-matched controls (young adult: p = 0.084, two-sample t-test; aged: p = 0.36, MW test). The inset graph shows the same data plotted per mouse (young adult: p = 0.71; aged: p = 0.94, two-sample t-test). **E.** The speed score of interneurons, calculated using periods of mobility in the maze, was reduced in both young adult and aged CA3-APP mice compared to their respective age-matched controls (young adult: p < 0.0001; aged: p = 0.0024, two-sample t-test). Horizontal lines depict the mean of all cells in a group, and open circles the mean per mouse. All statistics are reported in **Supplementary Table 5**.

We then analyzed the burst activity of CA1 principal cells during the task and identified a reduction in young adult (control 0.28 ± 0.01, CA3-APP 0.23 ± 0.01, p < 0.0001, MW test), but not in aged CA3-APP mice (control 0.24 ± 0.01, CA3-APP 0.24 ± 0.01, p = 0.21, MW test) compared to age-matched controls (**Supplementary Figure 2D**). However, the lack of effect for the comparison between the aged groups arises from a reduction in the burst index of control principal cells in aged mice (**Supplementary Figure 3I**). Next, we evaluated the firing of putative CA1 interneurons recorded from young adult mice and found similar average firing rates between genotypes (control 26.3 ± 0.7 Hz, CA3-APP 28.6 ± 1.2 Hz, p = 0.084, two-sample t-test) (**Figure 2D**). In the aged group, the mean firing rate of interneurons did also not differ (control 23.4 ± 1.3Hz, CA3-APP 21.6 ± 1.7Hz, p = 0.36, MW test), but there was a notable increase in the number of cells with abnormally low or abnormally high firing rates in aged CA3-APP mice (p = 0.0002, Kolmogorov-Smirnov test, **Figure 2D**). Hippocampal interneurons are known to be reliably speed modulated – a characteristic that could be used for updating position information during spatial navigation^20^. Interestingly, we observed a decrease in the speed modulation of CA1 interneurons during the behavioral task in both CA3-APP age groups (young adult: control 0.15 ± 0.01, CA3-APP 0.11 ± 0.01, p < 0.0001, two-sample t-test; aged: control 0.15 ± 0.02, CA3-APP 0.05 ± 0.001, p < 0.0001, two-sample t-test) (**Figure 2E**). Despite the decrease in speed modulation compared to controls, interneurons from young adult CA3-APP mice still retained some speed modulation (median value of speed modulation compared to zero, Wilcoxon signed rank test p<0.0001 for both control and CA3-APP; **Figure 2E**). Also, interneurons in aged control mice remained speed modulated (p < 0.0001, Wilcoxon signed rank test), but speed modulation in aged CA3-APP mice was no longer detectable (p = 0.67 Wilcoxon signed rank test) (**Figure 2E**). Consistent with the modulation of firing rate by running speed in only the control mice, we observed higher average firing rates of interneurons when the velocity was higher during behavior sessions compared to the sleep sessions (**Supplementary Figure 3A, C, E**). Taken together, we thus observed a decrease in the firing rate of CA1 principal cells with CA3-APP expression, which is exacerbated in aged mice. Conversely, the overall mean firing rate of interneurons remained similar across genotypes at both age points, but with diminished speed modulation in young adult and, to a larger extent, in aged CA3-APP mice.

### CA1 place cell coding remained unaltered despite changes in firing rate

In order to identify possible mechanisms underlying the deficits in spatial memory observed in our animal model we compared the spatial firing patterns between CA3-APP and control mice while they performed the alternation task. We selected laps with a 10-s delay for our analysis, as this was the condition when CA3-APP mice showed the most pronounced behavioral impairment. A lap was considered as the period from one reward location to the next reward location (**Figure 3A-B**). Only correct laps were used for the analysis to exclude effects from taking different turns at the choice point. First, we asked whether there was a difference in the fraction of principal cells coding for place in the maze. We found that the percentage of place cells was similar between control and CA3-APP mice for both age groups (young adult: control 69.3 ± 5.6%, CA3-APP 62.8 ± 5.3%, p = 0.27; aged: control 61.4 ± 2.4%, CA3-APP 59.1 ± 5.2%, p = 0.87, MW test, **Figure 3C**). We then evaluated the properties of place cells that were active during the task. As expected, we found place cell coding for each segment of the maze (**Figure 3B**) in left-turn and right-turn laps. Consistent with the reduction in mean firing rates (see **Figure 2**) we found a reduction in the peak firing rate of CA1 place cells in young adult CA3-APP mice and again to a larger extent in aged CA3-APP mice (young adult: control 20.3 ± 0.7 Hz, CA3-APP 17.7 ± 0.7 Hz, p = 0.0065; aged: control 21.2 ± 0.8 Hz, CA3-APP 14.8 ± 0.6 Hz, p = 0.001, MW test, **Figure 3D**). The spatial information carried by CA1 place cells nonetheless remained intact in CA3-APP mice at both age points (young adult p = 0.13; aged p = 0.15, MW test, **Figure 3E**). However, we found that CA1 place fields in CA3-APP mice had, on average, fewer place fields when compared to control littermates (young adult p = 0.028; aged p = 0.0083, MW test, **Figure 3F**) and that place fields in aged CA3-APP mice were slightly smaller (young adult p = 0.82; aged p < 0.0001, MW test, **Figure 3G**) and less stable than in controls (young adult p = 0.57; aged p = 0.015, MW, **Figure 3H**). It is important to note that, while significant, the effect sizes of these changes were small, and these minor changes could be a consequence of the reduced firing rates in CA3-APP mice.

**Figure 3.**
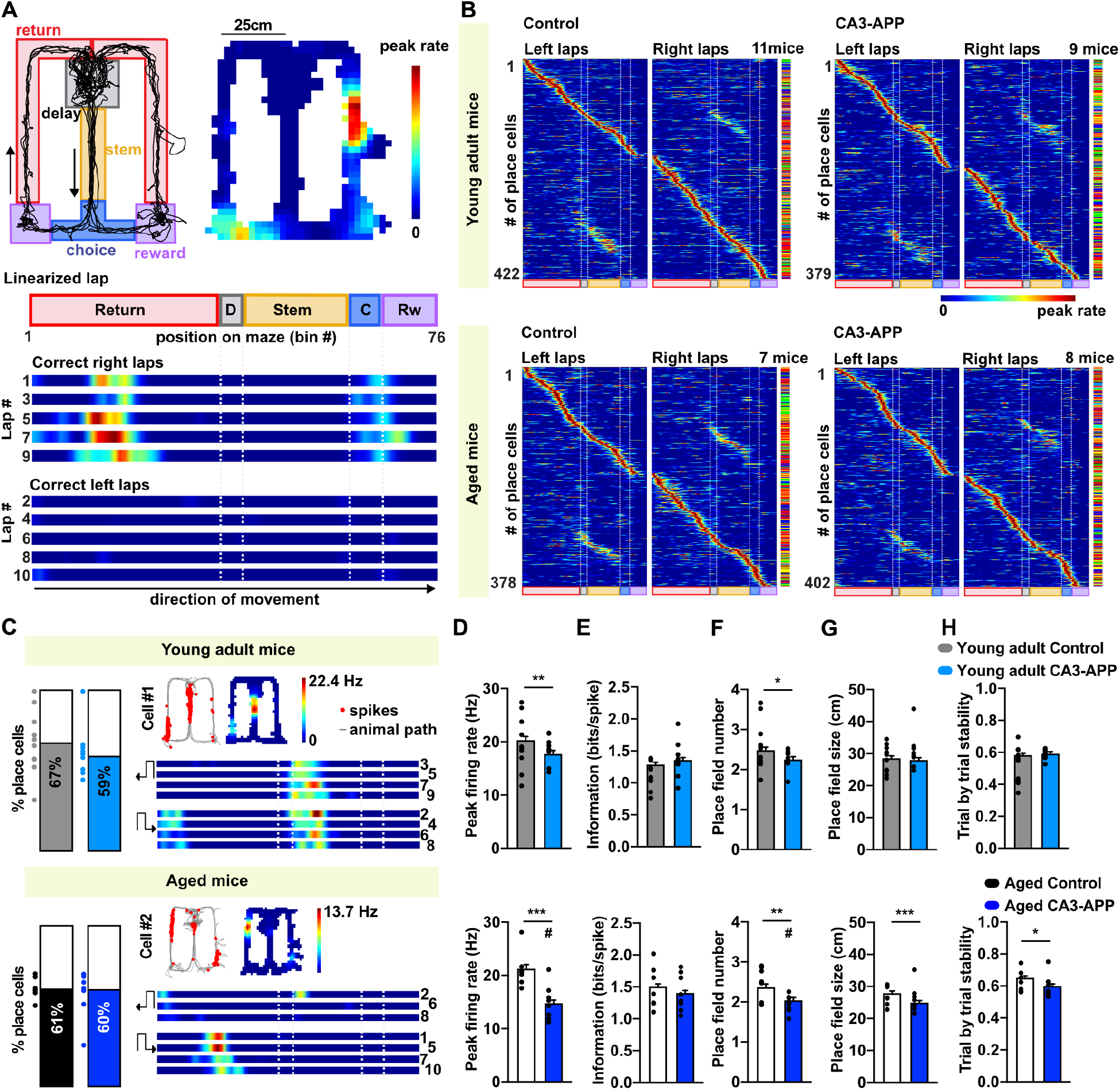
CA1 Place code remained accurate in CA3-APP mice. **A.** Left top: Example of mouse path (black) and parsing of figure-8 maze into sub-sections. The direction of movement is depicted by arrows. Right top: Example of a place cell recorded while the mouse ran in the maze. Spike density in two-dimensional space is averaged over all laps and shown as a color-coded map. Bottom: Place cell firing is shown separately for each lap in linearized, color-coded maps (76 bins of 2.5 cm). **B.** Average linearized rate maps for all CA1 place cells recorded in the four experimental groups. Left-turn and right-turn trials are shown separately. Cells are ordered by the location of the bin with the highest firing rate. In the vertical bar to the right of each panel, each individual mouse is assigned a color to show the distribution of cells across mice. **C.** Left: % CA1 principal cells with place fields. Dots represent the same data per mouse. Right: Example linearized rate maps of place cells from a young adult (top) and from an aged CA3-APP mouse (bottom). **D.** Peak firing rates of CA1 place cells were reduced in young adult CA3-APP and aged CA3-APP mice compared to their respective age-matched controls (young adult: p = 0.0065; aged: p = 0.001, MW test). In aged mice, the difference was also statistically different when data was analyzed per mouse (indicated by #, p = 0.0032, n = 7 for control and 8 for CA3-APP mice). **E.** Information content of place cells from CA3-APP mice did not differ from aged-matched control mice. **F.** The number of place fields per place cell was slightly lower in CA3-APP compared to aged-matched control mice (young adult: p = 0.028; aged: p = 0.0083, MW test). For cells from aged CA3-APP mice compared to age-matched controls, the difference was also significantly different when data was analyzed per mouse (represented by #, p = 0.0057, n = 7 for control and 8 for CA3-APP mice). **G.** Place field size of CA1 place cells from aged CA3-APP mice was smaller than from cells from aged-matched control mice (p < 0.0001, MW test). In young adult CA3-APP mice the difference was not statistically significant (p = 0.82, MW test). **H.** Stability of place cells’ firing was analyzed by a lap-by-lap correlation across location bins. A small reduction in stability was found in aged CA3-APP mice compared to aged-matched control mice (p = 0.015, MW test), but no difference was observed in young adult mice (p = 0.57, MW test). For D-H, bar graphs represent the data per cell, and black dots represent the average per mouse. Error bars correspond to SEM for the cell-wise analysis. All statistics are reported in **Supplementary Table 6**.

To successfully perform the spatial alternation task, mice needed to process information about the immediately preceding trajectory and retain the information before making the next choice. A mechanism for retaining behaviorally relevant information is trajectory-dependent firing that begins on the return arm and continues during the delay. We therefore first examined how distinct the firing is between left and right sides of the maze^21,22^ by computing spatial correlations between the left and right maze sections. Lower spatial correlations correspond to more distinct spatial coding between the two sides, and we did not find any differences to age-matched controls for either young adult or aged CA3-APP mice (**Supplementary Figure 4D**), suggesting that place coding remained arm-selective with CA3-APP expression. Next, we examined whether the information about the preceding trajectory is retained during the delay intervals, which might support memory-guided behavior and spatial navigation^21–23^. We observed cells that fired differently in the central arm depending on whether mice came from the left or right side and were about to perform a right or left correct turn in the maze in all our experimental groups (young adult: control 35.4 ± 4.9%, CA3-APP 26.1 ± 4.8%, p = 0.20; aged: control 33.9 ± 3.9%, CA3-APP 26.4 ± 5.2%, p = 0.24, two-sample t-test), and did not detect differences in the spatial properties of cells on the center arm other than the reduced place field size that was also detected in other maze segments (**Supplementary Figure 4F-H**). In summary, despite rate differences, place coding remained robust in the CA1 region of CA3-APP mice. Our observation contrasts with the marked degeneration of the CA1 place code found in animal models with pan-neuronal expression of AD-related proteins^11^ and indicate that early deficits in spatial memory in AD might not be directly correlated with a severely degraded hippocampal place code.

### LFP oscillations in the theta range were shifted in frequency in CA3-APP mice

Since spatial coding remained accurate, even in aged CA3-APP mice with pronounced memory deficits, we next examined whether there were other functional deficits that could underlie deficits in spatial working memory. Brain oscillations are known to organize spiking activity and are linked to memory processes^23^. Using the same maze segments as for the analysis of firing rates, we analyzed hippocampal oscillations in different regions of the maze. To be able to analyze oscillations without preselecting different frequency ranges, we first normalized the LFP signal in the working memory task to the LFP recorded during rest periods in the home cage (running speed < 2 cm/s) before the start of the memory task (see **Supplementary Figure 5A-B**). Using the normalized LFP of all laps, we then constructed a spectrogram per mouse along a linearized path in which each bin represents the average time spent in each region of the maze (**Supplementary Figure 6A-C**, see Methods for more details). The normalized spectrograms allowed for the unbiased identification and definition of LFP bands that were relevant during different phases of the memory task (**Figure 4A**). Based on the normalized spectrograms of control mice, we defined the boundaries of five evident LFP bands: delta (2-6 Hz), theta (6-12 Hz), beta (14-26 Hz), low gamma (26-50 Hz) and high gamma (50-120 Hz). Then, for each part of the maze, we computed a power spectrum density function (PSD) of the LFP signal and determined the peak frequency at each one of the defined LFP bands (**Figure 4A and D**, **Supplementary Figure 6**). In analyses of oscillation frequencies, we identified a reduction in the peak frequency of theta (control 10.5 ± 0.1 Hz, CA3-APP 9.7 ± 0.2 Hz, p = 0.037) and high-gamma oscillations (control 93.7 ± 2.9 Hz, CA3-APP 76.2 ± 2.8, p = 0.0062, MW test) recorded in the CA1 pyramidal layer of young adult CA3-APP mice. The quantification for the central arm of the maze (with stem and choice regions combined) is shown in **Figure 4B**, but similar results were obtained for other regions of the maze (**Supplementary Figures 6 and 7**). Importantly, frequency differences did not emerge from differences in running speed profiles of the mice – speed differences were mostly present at the choice point (**Supplementary Figure 5C, D**) while theta frequency differences were observed at all phases and when controlling for velocity (**Supplementary Figure 5 E, F**). The peak frequency of delta (3.5 ± 0.1 Hz, 3.1 ± 0.1 Hz, p = 0.14), beta (20.6 ± 0.2 Hz, 19.6 ± 0.2 Hz, p = 0.074), and gamma oscillations (33.2 ± 1.1 Hz, 32.0 ± 1.1 Hz, p = 0.58, MW test) was similar between control and CA3-APP mice during the memory task, and no differences in normalized power were detected for any of the analyzed LFP bands (**Figure 4C**). Compared to young adult CA3-APP mice, the reduction in the frequency of theta oscillations was even more pronounced in CA1 LFP recordings of aged CA3-APP mice (control 10.1 ± 0.2 Hz, CA3-APP 8.8 ± 0.2 Hz, p = 0.00014, MW test, **Figure 4D and E**), but again with no differences in power. High gamma oscillations were also slower in the aged CA3-APP mice (control 98.4 ± 4.3 Hz, CA3-APP 86.7 ± 3.5 Hz, p=0.042) while delta (control 3.2 ± 0.1 Hz, CA3-APP 3.4 ± 0.1 Hz, p = 0.21), beta (control 19.9 ± 0.5 Hz, 18.6 ± 0.6 Hz, p = 0.056) and low gamma oscillation frequencies (control 35.9 ± 1.3 Hz, CA3-APP 34.9 ± 0.8 Hz, p = 0.47, MW test) were not different from age-matched controls. We found a small decrease in the power of the beta band (which overlaps with the second harmonic of theta) in aged CA3-APP mice compared to age-matched controls (control 1.6 ± 0.1, CA3-APP 1.3 ± 0.05, p = 0.043, MW test), which was most prominent in the central arm of the maze, particularly in the choice section (**Figure 4D, F**, **Supplementary Figure 6G**). No other differences in the power of hippocampal oscillations were found in aged CA3-APP mice compared to controls (see **Supplementary Table 9**). It is important to note that aged CA3-APP mice move slower, specifically at the choice section of the maze (**Supplementary Figure 5D**). We therefore analyzed the relationship between the velocity of the mouse and frequency and power of theta oscillations. Our analysis revealed that the frequency reduction in the choice segment cannot be explained by differences in running speed (**Supplementary Figure 5E, F**), as changes is frequency were seen at all velocities. Furthermore, theta oscillation frequency was also reduced in other parts of the maze, such as the return arms or the stem, where the velocity of the animals did not differ (**Supplementary Figure 5E, F**).

**Figure 4.**
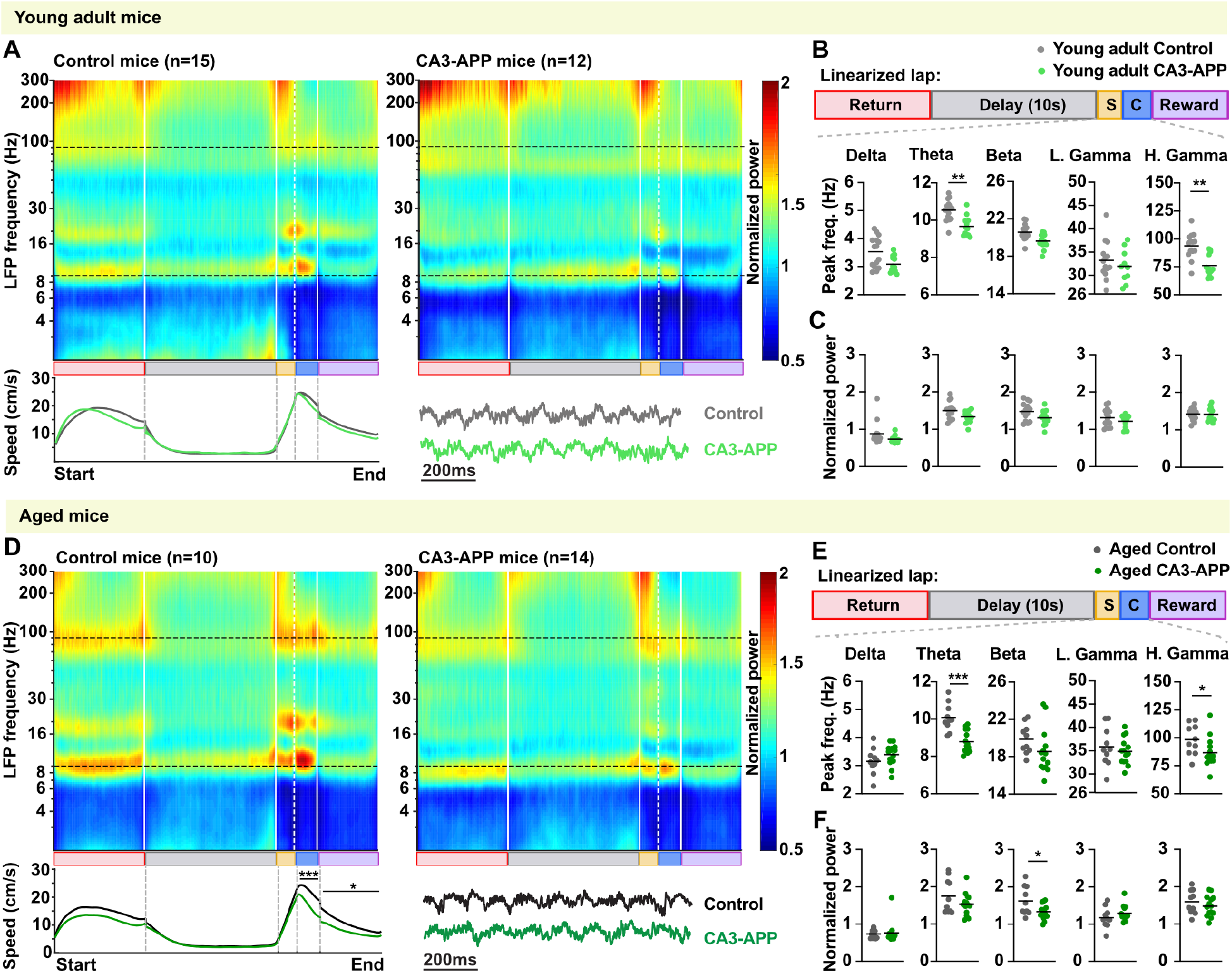
Theta and gamma oscillation frequencies were reduced in young adult and aged CA3-APP mice. **A.** Average CA1 LFP spectrogram of a lap for young adult and aged CA3-APP mice. An average spectrogram was calculated per session and all sessions were combined, first per mouse and then over all mice of the same genotype. The average spectrogram per genotype is shown. The plot at the bottom of each panel shows the running speed of mice across the different sub-sections of a lap in the figure-8 maze. Young adult CA3-APP mice (n = 15 mice) showed no difference in running speed in the maze compared to young adult control mice (n = 12 mice). A more detailed analysis of running speed data is presented in **Supplementary Figure 5C and Supplementary Table 7**. **B.** Quantification of frequency in different LFP bands. Quantification is shown for the center arm (including stem and choice point). Analyses for other maze sections are provided in **Supplementary Figure 6 and Supplementary Table 9** and generally follow the same pattern. The theta (p = 0.037) and high gamma frequencies (p = 0.0062, MW test adjusted for multiple comparisons by Holm-Šídák method) of young adult CA3-APP mice were reduced compared to age-matched controls. **C.** Quantification of power in different LFP frequency bands. We found no difference in CA1 LFP power between young adult CA3-APP mice and their control littermates. **D.** Aged CA3-APP mice show a small reduction in running speed which is only present at the choice and reward sections of the maze (n = 10 for control and n = 14 for CA3-APP mice). Data are displayed as in A. **E.** Similar to young adult mice, aged CA3-APP mice showed a pronounced reduction in theta oscillation frequency compared to age-matched control mice (p = 0.00014), and a reduction in high gamma oscillation was also found (p = 0.042, MW test adjusted for multiple comparisons by Holm-Šídák method). Data are displayed as in B. **F.** Power in the Beta band was decreased in aged CA3-APP mice compared to control mice (p = 0.043, MW test adjusted for multiple comparisons by Holm-Šídák method). Data are displayed as in C. This difference is only found in the choice zone (see **Supplementary Figure 6E** for bysection analysis).

### Temporal properties of principal cells and interneurons were altered in CA3-APP mice

Since LFP oscillations reflect the temporally coordinated neuronal activity of inputs to a brain region and of local circuits^24,25^, we aimed to understand whether slower theta oscillations are associated with changes in the temporal activity patterns of principal cells and interneurons in the CA1 region. We first measured the oscillation frequency of the spiking of principal cells and confirmed that a similar percentage of cells from young adult control and CA3-APP mice had a peak within the theta band (85.7% and 88.1%, respectively, p = 0.25, Chi-square test, **Supplementary Figure 8A**). These cells showed a decrease in the oscillation frequency of the spike patterns with CA3 APP expression (9.9 ± 0.1 Hz for control and 9.4 ± 0.1 Hz for CA3-APP young adult mice, p < 0.0001, MW test **Figure 5A**), but no difference in their theta modulation amplitude (control 0.94 ± 0.01, CA3-APP 1.11 ± 0.01, p = 0.17, MW test, **Figure 5B**). As previously described^26^ the average frequency of CA1 principal cell spiking is faster than the LFP, which was also observed here for all groups of mice. However, each cell’s frequency and the ongoing LFP frequency were shifted to similar extents, such that the frequency difference did not differ between control and CA3-APP mice (control 1.2 ± 0.1 Hz, CA3-APP 1.2 ± 0.1 Hz, p = 0.81, MW test, **Supplementary Figure 8A**). In line with this, the relationship of CA1 principal cells with the ongoing theta oscillations seems otherwise normal in young adult CA3-APP mice as they have similar preferred phase of firing (control 142 ± 5°, CA3-APP 140 ± 5°, p = 0.93, MW test) and a similar extent of theta phase locking (control 0.15 ± 0.01, CA3-APP 0.16 ± 0.01, p = 0.056, MW test, **Supplementary Fig 8B**).

**Figure 5.**
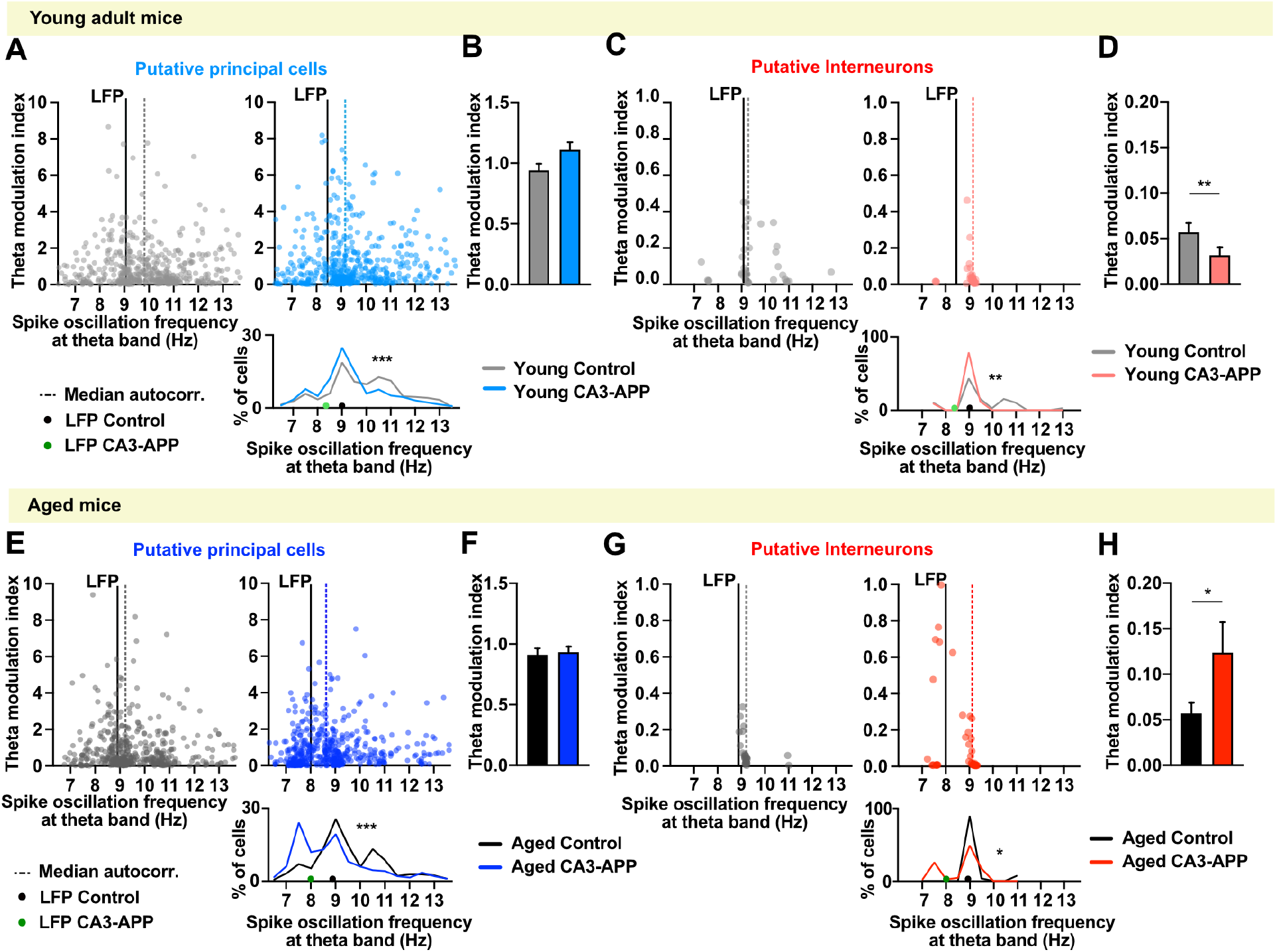
Spike oscillation frequency of CA1 principal cells and interneurons was reduced in CA3-APP mice. **A.** For each neuron we calculated the spike-time autocorrelation and then used FFT analyses to obtain the predominant spike oscillation frequency. The distribution of the peak oscillation frequencies is plotted against the amplitude of their theta modulation (i.e., theta modulation index). Only cells with a detectable peak within the theta band are displayed, and the dashed line represents the median of their frequency distribution. The mean LFP frequency of the sessions in which these cells were recorded is indicated by the solid black line. Spike oscillation frequencies of CA1 principal cells were lower in young adult CA3-APP mice than in littermate controls (n = 481 cells for control, n = 488 cells for CA3-APP, p < 0.0001, MW test). **B.** Average theta modulation amplitude (theta modulation index) of all active CA1 principal cells. There was no difference between cells from young adult CA3-APP compared to cells from age-matched mice (n = 561 cells for control, n = 554 cells for CA3-APP, p = 0.1732, MW test). **C.** In young adult CA3 APP mice, there were no interneurons with high spike oscillation frequencies, which resulted in a different distribution than in young adult controls (n = 39 cells for control, n = 34 cells for CA3-APP, p = 0.0024, MW test). Data are displayed as in A. **D.** Theta modulation amplitude of CA1 interneurons recorded from young adult CA3-APP mice was lower than for interneurons from age-matched control mice (n = 90 cells for control, n = 60 cells for CA3-APP, p = 0.0039, MW test). **E.** When compared to age-matched controls, CA1 interneurons from aged CA3-APP mice showed a major reduction in the spike oscillation frequency with a large fraction of cells oscillating between 7 and 8 Hz (n = 452 cells for control, n = 484 cells for CA3-APP, p < 0.0001, MW test). Data are displayed as in A. **F.** Theta modulation amplitude of CA1 principal cells of aged CA3-APP mice did not differ from age-matched controls (n = 499 cells for control, n = 527 cells for CA3-APP, p = 0.46, MW test). **G.** When compared to controls, CA1 interneurons from aged CA3-APP mice showed reduced spike oscillation frequencies, with >20% of interneurons now oscillating below the LFP frequency (n = 28 cells for control, n = 47 cells for CA3-APP, p = 0.029, MW test). Data are displayed as in A. **H.** CA1 Interneurons from aged CA3-APP mice showed an increase in theta modulation amplitude compared to age-matched controls (n = 41 cells for control, n = 50 cells for CA3-APP, p = 0.038, MW test). All statistics are reported in **Supplementary Table 11**.

For CA1 interneurons, a similar percentage of cells from young adult control (43.3%) and CA3-APP mice (56.7 %) had spike patterns with rhythmicity within the theta band (p = 0.11, Chi-square test), but the average frequency was lower in CA3-APP than in control mice (control: 9.5 ± 0.2 Hz, CA3-APP: 9.0 ± 0.1 Hz; p = 0.0024, MW test, **Figure 5C**). Again, the differences between each interneuron’s frequency and the LFP frequency remained preserved with APP expression (control 0.9 ± 0.3 Hz, CA3-APP 0.8 ± 0.1 Hz, p = 0.33, MW test, **Supplementary Figure 8C**). A small reduction in the amplitude of theta modulation was found for CA1 interneurons in young adult CA3-APP compared to control littermates (control 0.06 ± 0.01, CA3-APP 0.03 ± 0.01, p = 0.0039, MW test; **Figure 5D**), but without differences in phase preference (control 165 ± 13°, CA3-APP 154 ±16°, p = 0.40) or phase locking (control 0.15 ± 0.11, CA3-APP 0.14 ± 0.01, p = 0.54) (**Supplementary Figure 8D**). Overall, in young adult mice with APP expression, there was a minor reduction of the frequency of principal cells and interneurons, but this occurred in parallel with a reduction of the LFP frequency such that the frequency differences between the two types of oscillations were preserved.

In aged mice, most active CA1 principal cells had spike patterns with rhythmicity within the theta band (90.6 % for control and 91.8% for CA3-APP, p = 0.48, Chi-square test), but cells in CA3-APP mice showed a pronounced reduction in their frequency (control: 9.5 ± 0.1 Hz, CA3-APP: 8.8 ± 0.1 Hz; p < 0.0001, MW test), with a subset of about 20% of the cells oscillating at frequencies as low as ~7.5 Hz (**Figure 5E**). Nevertheless, the distance between the preferred frequency of individual cells and the LFP frequency did not differ from control littermates (control 0.9 ± 0.1 Hz, CA3-APP 0.9 ± 0.1 Hz, p = 0.46, MW test, **Supplementary Figure 8E**), indicating that cells’ oscillations were altered along with the network oscillations. As observed in young adult CA3-APP mice, we found no differences in theta modulation amplitude (control 0.06 ± 0.01, CA3-APP 0.03 ± 0.01, p = 0.46, MW test, **Figure 5F**), preferred phase (control 165 ± 13°, CA3-APP 154 ± 16°, p = 0.90, MW test) or phase locking of principal cells (control 0.15 ± 0.01, CA3-APP 0.14 ± 0.01, p = 0.96, MW test, **Supplementary Figure 8F**). For CA1 interneurons, 94.0% from aged CA3-APP mice were theta modulated, in contrast to 68.3% in control mice (p = 0.0013, Chisquare test). Resembling the frequency patterns in CA1 principal cells, a subset of CA1 interneurons (~20%) from aged CA3-APP mice was oscillating at ~7.5 Hz. The effect was sufficiently pronounced in interneurons to result in a decrease in the average spike oscillation frequency (8.7 ± 0.1 Hz, compared to 9.3 ± 0.1 Hz in control mice, p = 0.029, MW test, **Figure 5G**), but the shift was less than for the LFP frequency so that the difference between the CA1 interneurons spike oscillation frequency and the LFP frequency was significantly larger in aged CA3-APP mice compared to age-matched controls (control 0.4 ± 0.1, CA3-APP 1.0 ± 0.1, p = 0.0003, **Supplementary Figure 5G**). Furthermore, we observed a substantial increase in theta modulation amplitude of CA1 interneurons with APP expression (control 0.06 ± 0.01, CA3-APP 0.12 ± 0.03, p = 0.0380, MW test; **Figure 5H**). The increase in theta modulation was most pronounced in the subset of interneurons oscillating at the reduced ~7.5 Hz frequency. Again, we did not find changes in preferred theta phase (control 206 ± 22°, CA3-APP 203 ±18°, p = 0.97, MW test) or phase locking (control 0.19 ± 0.01, CA3-APP 0.23 ± 0.01, p = 0.068, MW test; **Supplementary Figure 5H**) of CA1 interneurons. In summary, the shift in the oscillation frequency in aged mice was more pronounced than in young adult mice and was mostly due to the appearance of a subset (~20-30%) of both principal cells and interneurons now oscillating at a lower frequency of 7-8 Hz.

### Impairments in phase precession and pairwise timing in aged CA3-APP mice

Place cells in the hippocampus are known to shift their firing to earlier phases of the theta cycle as the animal transverses its place field (‘theta phase precession’)^26,27^. We asked whether the changes in theta-related temporal organization resulted in the reorganization of the cell’s temporal firing patterns with respect to LFP and with respect to each other. To address this, we analyzed phase shifts over the course of spike trains (see methods) and found that the majority of CA1 place cells showed some level of phase precession (**Figure 6A-C**). Place cells from young adult CA3-APP mice had control levels of phase precession (Median slope: control −0.27, CA3-APP −0.26, p = 0.97), while aged CA3-APP mice showed a small reduction in phase precession compared to age-matched controls (control −0.39, CA3-APP −0.30; p = 0.028, MW test). The phase precession offset (Phi0) was also reduced in aged CA3-APP mice (control −0.44 ± 0.045π, CA3-APP −0.26 ± 0.048π, p = 0.011, MW test) but not in young adult mice (control −0.17 ± 0.048π, CA3-APP −0.19 ± 0.049π; p = 0.62, MW test). Interestingly we only saw a reduction in phase precession when the task was hippocampus-dependent (2-s and 10-s delay), but not during the continuous version of the task. (**Supplementary Figure 9A, B**).

**Figure 6.**
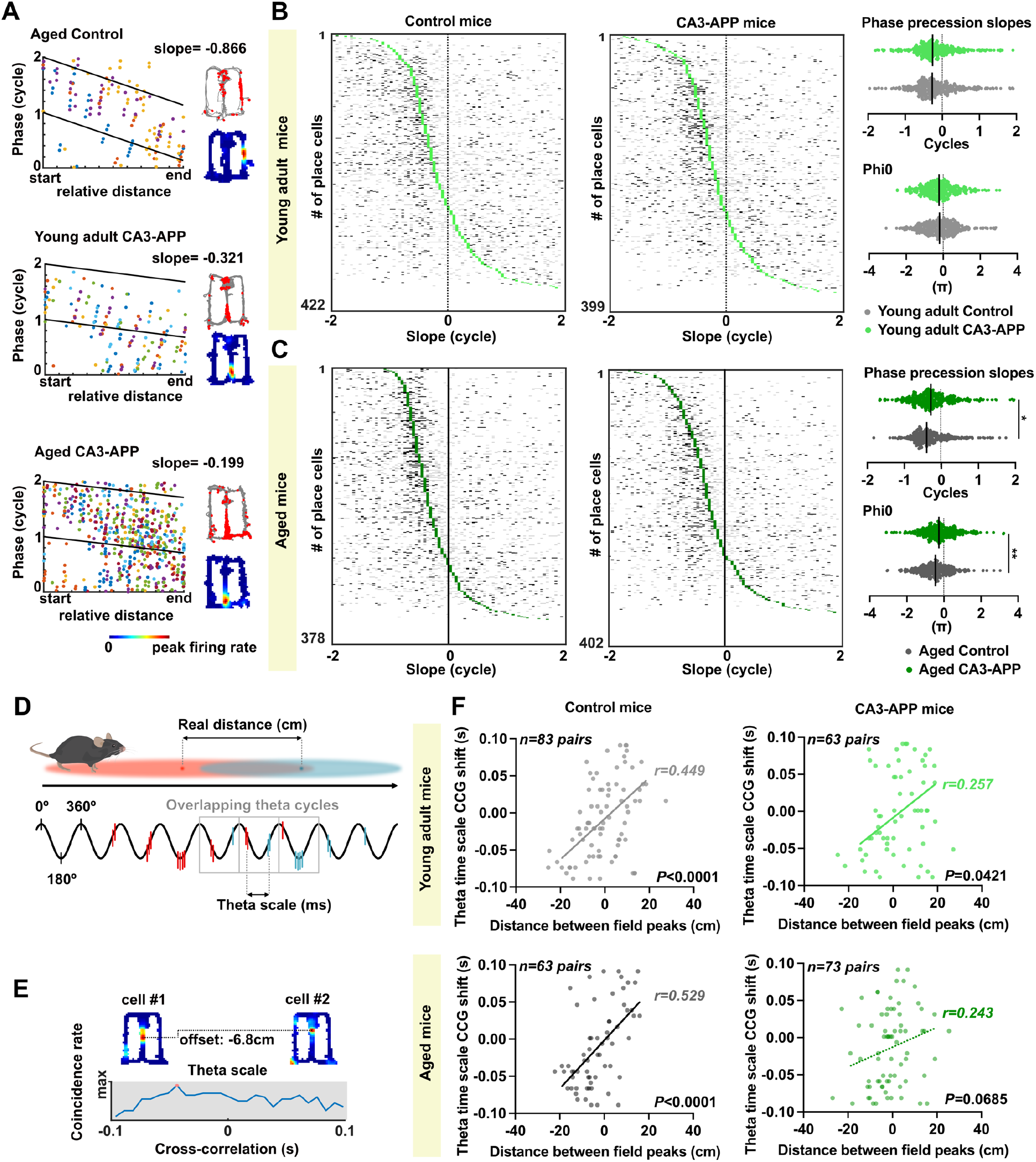
Phase precession and theta sequences were impaired in aged but not in young adult CA3-APP mice. **A.** Phase precession slopes of example CA1 place cells. Each spike train in the phase-distance plot is depicted in a different color. Place maps of the corresponding cells are shown to the right. Top, spike locations (red dots) plotted on the mouse’s path (gray line). Bottom: spike rate map color-coded with red as the peak rate. **B.** Distribution of phase precession slopes. Each row shows the phase precession slope of each spike train (gray ticks, non-significant; black ticks, significant) of one place cell in the figure-8 maze. The median (green tick) was calculated by including significant and non-significant slopes, and cells are ordered by their median slope (in green). Phase precession (n = 386 out of 422 place cells for control and n = 351 out of 399 for CA3-APP, p = 0.097, MW test) and phase offset (p = 0.62, MW test) in young adult CA3-APP mice did not differ from littermate controls. **C.** Same as in B, but for aged CA3-APP mice. There was a reduction in phase precession slope in aged CA3-APP mice compared to littermate controls (n = 342 out of 378 place cells for control and n = 351 out of 402 for CA3-APP, p = 0.028, MW test), and the phase offset was closer to zero in CA3-APP mice compared to controls (p = 0.0019, MW test). **D.** Schematic for the comparison between the spatial distance of CA1 place field peaks and the temporal distance within a theta cycle. Theta-time scale firing distance is measured as the time bin with the peak cross-correlation. Both measurements can be obtained from pairs of overlapping place fields. **E.** Example metrics obtained from a pair of simultaneously recorded cells with overlapping place fields. **F.** Correlation between place field distance and shift in theta-time scale firing was found in young adult mice of both genotypes (Spearman correlation p < 0.0001, n = 83 and p = 0.042, n = 63 pairs of overlapping place fields for control and CA3-APP respectively) and in aged control mice (Spearman correlation n = 63 pairs of overlapping place fields, p < 0.0001), but not in aged CA3-APP mice (Spearman correlation n = 73 pairs of overlapping place fields, p = 0.069). All statistics are reported in **Supplementary Table 12**.

We then evaluated whether the reduction in phase precession during the memory task could alter the pairwise temporal organization of CA1 place cells. For pairs of simultaneously recorded cells with overlapping fields, we computed their distance on the maze (behavioral scale) and their temporal distance within a theta cycle (theta time scale). As expected when behavioral sequences correspond to time-compressed sequences within the theta cycle^28^, we saw a clear correlation between distance and theta time in young adult and aged control mice (young and aged: p <0.0001, Spearman Correlation test). However, the correlation was weaker in young CA3-APP mice (p = 0.048) and no longer significant in aged CA3-APP mice (control p <0.0001, CA3-APP p = 0.070, **Figure 6D**). As a consequence, the slope of the relation between theta time scale versus behavioral scale (i.e., sequence compression) was reduced in aged CA3-APP mice (**Supplementary Figure 10E**), which indicates that the precise temporal order within the theta cycle no longer consistently corresponded to the sequential activation in the maze.

### Network alterations in the CA3/DG region of CA3-APP mice were not more severe than in CA1

Unlike previous studies, we performed recordings from cells that did not directly expressed hAPP, such that physiological deficits can be assumed to arise from dysfunctional synaptic inputs. However, the observed effects were most pronounced for temporal firing patterns, and it is conceivable that more severe deficits could have been detected when directly recording from the neuron population with APP expression. We therefore next performed analyses of spatial and temporal firing patterns of recordings from the CA3/DG regions of the hippocampus (**Figure 7A, D**) and asked whether we could identify larger deficits. Similar to the CA1 region, we found a reduction in theta frequency in the CA3 region of aged CA3-APP mice (control 9.4 ± 0.2 Hz, CA3-APP 8.5 ± 0.1 Hz, p = 0.0029, **Figure 7E**), and again a less pronounced reduction in young adult CA3-APP. In CA3, the difference did not reach significance in young adult CA3-APP mice compared to age-matched control recordings from CA3/DG (control 10.4 ± 0.2 Hz, CA3-APP 9.7 ± 0.2 Hz, p = 0.30792, MW test, **Figure 7B**), but was nonetheless comparable to the effects observed in CA1 recordings (p > 0.999, Kruskal-Wallis test with Dunn’s multiple comparisons test). For the frequency of high gamma oscillations, the difference did not reach significance for the CA3/DG recordings (young adult control 62.7 ± 1.6 Hz, CA3-APP 59.8 ± 0.9 Hz, p = 0.091; aged control 64.7 ± 1.7 Hz, CA3-APP 61.7 ± 0.9 Hz, p = 0.74, **Figure 7B, E, Supplementary Figure 11A**). Since CA3/DG LFP recordings did not show higher levels of dysfunction than CA1 LFP recordings, we next asked to what extent the spatial and temporal firing properties cells in this region were altered. Unlike what was observed in CA1, the firing rates of CA3/DG principal cells remained unchanged in both age groups of CA3-APP mice (young adult control 0.84 ± 0.06 Hz, CA3-APP 0.80 ± 0.03 Hz, p = 0.058; aged control 1.2 ± 0.01 Hz, CA3-APP 1.1 ± 0.1 Hz, p = 0.96, MW test, **Supplementary Figure 13A, B**), but we detected a reduction in the bursting activity of principal cells of CA3-APP mice (young adult control 0.24 ± 0.1, CA3-APP 0.20 ± 0.004, p <0.0001; aged control 0.28 ± 0.01, CA3-APP 0.23 ± 0.01, p <0.0001), which was comparable to CA1. Similar to CA1, CA3/DG interneurons also did not show differences in their mean firing rate (young adult control 22.9 ± 1.4 Hz, CA3-APP 21.6 ± 0.9 Hz, p = 0.65; aged control 22.6 ± 1.7 Hz, CA3-APP 20.5 ± 1.0 Hz, p = 0.37), but had reduced speed modulation in young adult CA3-APP mice (control 0.20 ± 0.02, CA3-APP 0.11 ± 0.02, p = 0.0002) and aged CA3-APP mice (control 0.03 ± 0.04, CA3-APP 0.1 ± 0.02, p = 0.60, MW test, **Supplementary Figure 13F**). However, normal aging also seems to strongly affect speed modulation in interneurons recorded from the CA3/DG areas, as control aged mice lack speed modulation (p = 0.42, Wilcoxon signed rank test, **Supplementary Table 15**) such that there was no longer a statistical difference between aged control and CA3-APP mice.

**Figure 7.**
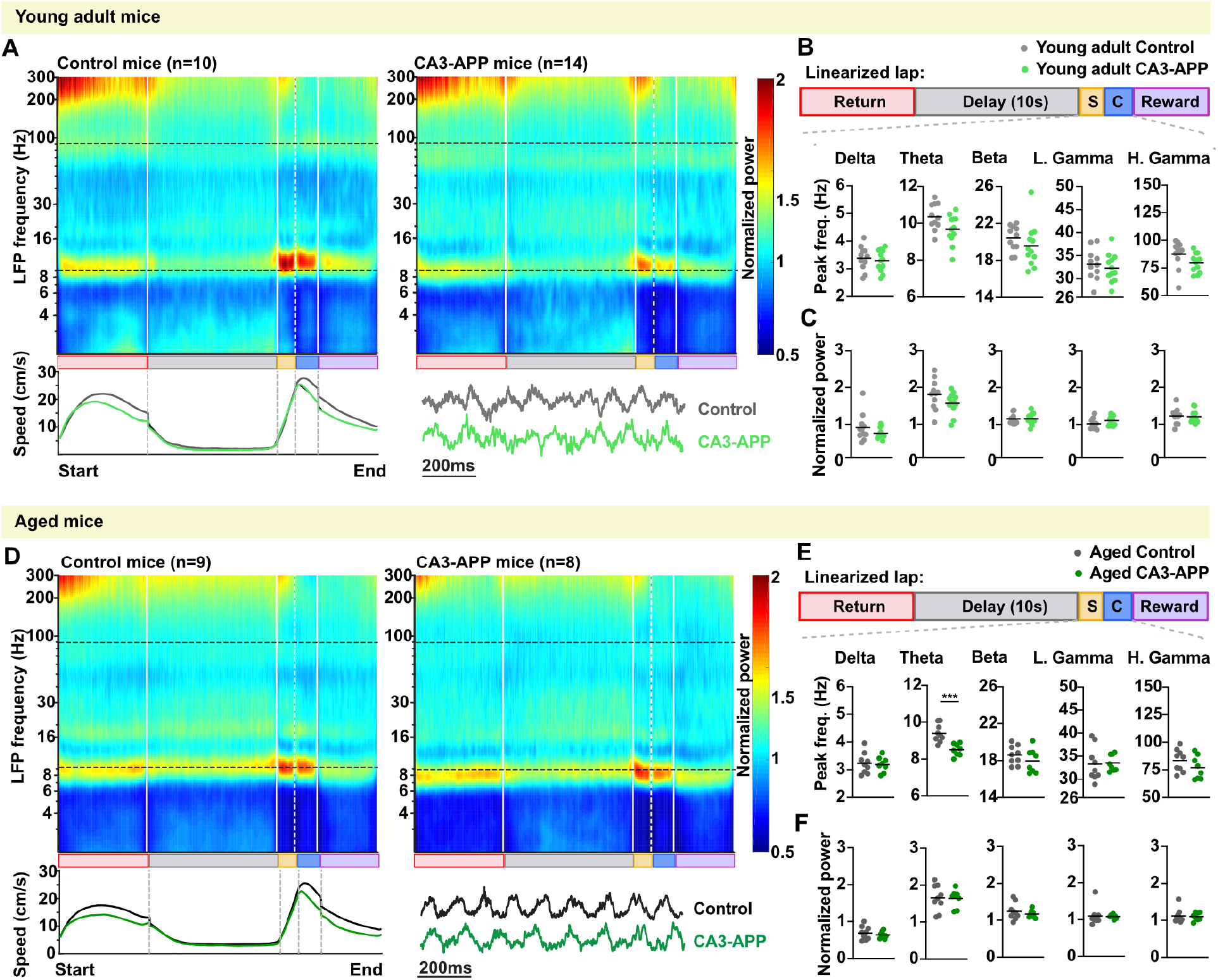
At recording sites in the CA3/DG area, proximity to hAPP expression in CA3 resulted in lesser or comparable effects on LFP frequency compared to effects at CA1 recording sites. **A.** Average CA3/DG LFP spectrogram is depicted as in Figure 4A. The plot at the bottom of each panel shows the running speed of mice across the different sub-sections of a lap in the figure-8 maze. Young adult CA3-APP mice (n = 9) showed no difference in running speed in the maze compared to young adult control mice (n = 14 mice). A more detailed analysis of running speed data is presented in **Supplementary Figure 5C and Table 7**. **B.** Quantification of frequency in different LFP bands. Quantification is shown for the center arm (including stem and choice point). Analyses for other maze sections are provided in **Supplementary Figure 11 and Supplementary Table 13** and generally follow the same pattern. The theta (p = 0.31) and high gamma frequencies (p = 0.091, MW test adjusted for multiple comparisons by Holm-Šídák method) recorded in the CA3/DG region of young adult CA3-APP mice did not differ from age-matched controls (p > 0.999, Kruskal-Wallis test with Dunn’s multiple comparisons test). **C.** Quantification of CA3/DG power in different LFP frequency bands. LFP power did not differ between young adult CA3-APP mice and their control littermates (p > 0.05, MW test adjusted for multiple comparisons by Holm-Šídák method). **D.** The running speed of aged CA3-APP did not differ from littermate controls. Data are displayed as in A. **E.** Aged CA3-APP mice showed a pronounced reduction in theta oscillation frequency in the CA3/DG area compared to age-matched control mice (p = 0.0029, MW test adjusted for multiple comparisons by Holm-Šídák method). Data are displayed as in B. **F.** CA3/DG LFP power did not differ between young adult CA3-APP mice and their control littermates. Data are displayed as in C. All statistics are reported in **Supplementary Table 13**.

### Spatial and temporal properties of principal cells and interneurons in the CA3/DG region of CA3-APP mice showed comparable effects to those in the CA1 region

We next evaluated whether place cells in CA3/DG were altered in CA3-APP mice. The percentage of place cells recorded in the CA3/DG region of CA3-APP mice was not changed in young adult or aged CA3-APP mice compared to littermate controls (young adult: control 64 ± 9%, CA3-APP 52 ± 8 %, p = 0.31, **Supplementary Figure 12A-C**; aged: control 66 ± 11%, CA3-APP 72 ± 10%, p = 0.70, two-sample t-test, **Supplementary Figure 12F, G**). While the peak firing rate of CA3/DG place cells was not altered in CA3-APP mice (young control 23.3 ± 1.5 Hz, CA3-APP 20.6 ± 0.8 Hz, p = 0.46; aged control 20.1 Hz ± 0.9, CA3-APP 18.7 ± 0.8 Hz, p = 0.70) their burst index was reduced in both young (control 0.25 ± 0.01, CA3-APP 0.19 ± 0.01, p <0.0001) and aged mice (control 0.30 ± 0.01, CA3-APP 0.24 ± 0.03, p <0.0001, MW test, **Supplementary Figure 12D, H**). Information content of CA3/DG place cells was reduced in CA3-APP mice already in young adults (control 1.7 ± 0.1, CA3-APP 1.5 ± 0.04, p = 0.013) and this difference remained in the aged groups (control 1.4 ± 0.03, CA3-APP 1.3 ± 0.04, p = 0.019, MW test). Furthermore, the number of place fields in the maze remained unaltered (young control 2.1 ± 0.1, CA3-APP 2.1 ± 1.2, p = 0.69; aged control 2.1 ± 0.1, CA3-APP 2.4 ± 0.1, p = 0.013), while place field size was significantly reduced in CA3/DG cells of young adult CA3-APP mice (control 22.0 ± 0.8, CA3-APP 25.5 ± 0.6, p = 0.0034) but did not reach significance in aged mice (control 23.9 ± 0.6, CA3-APP 24.3 ± 0.3, p = 0.75, MW test). No significant changes were found in trial by trial stability and or spatial correlation across sides in CA3-APP mice in both aged groups (**Supplementary Figure 12E, I**). Similarly to what we observed in CA1, a large portion of place cells displayed differences in the central arm between left-turn and right-turn trials and this was not altered in CA3-APP mice (young adult: control 39 ± 7%, CA3-APP 23 ± 5%, p = 0.10; aged: control 33 ± 7%, CA3-APP 43 ± 5%, p = 0.32, two-sample t-test, **Supplementary Figure 13A-C**). The information content and trial by trial stability of CA3/DG central-arm cells was not altered in CA3-APP mice of both age groups, but their place field size was larger in young mice (control 21.9 ± 1.1cm, CA3-APP 27.6 ± 1.2 cm, p = 0.001) but not aged CA3-APP (control 26.0 ± 1.4 cm, CA3-APP 27.5 ± 1.4 cm, p = 0.2360, MW test, **Supplementary Figure 12D-F**). In summary, similarly to CA1 place cells, there were no consistent patterns in place field changes with CA3-APP expression, and if significant differences were detected, the effect sizes for differences in place cell properties of CA3/DG neurons were quite small.

Furthermore, we found that changes in temporal firing properties of CA3/DG cells were quite similar to the ones we report for CA1 cells (**Figure 5**). There was already a reduction of the spike oscillation frequency of principal cells in young adult CA3-APP mice (control 9.9 ± 0.1 Hz, CA3-APP 9.3 ± 0.1Hz, p <0.0001, MW test, **Figure 8A**), but it was less than the reduction in LFP theta frequency such that the difference between the cells’ frequency and the LFP frequency increased (control 1.2 ± 0.1 Hz, CA3-APP 1.9 ± 0.1 Hz, p = 0.0009, MW test, **Supplementary Figure 16A**). At this age, interneurons also showed a small reduction in spike oscillation frequency (control 9.3 ± 0.1 Hz, CA3-APP 8.9 ± 0.1Hz, p = 0.016, MW test, **Figure 8C**) but without changes in the difference between the cells’ and the LFP oscillation frequency (control 1.4 ± 0.3 Hz, CA3-APP 1.4 ± 0.2 Hz, p = 0.64, MW test, **Supplementary Figure 16C** inset). No difference was found in the amplitude of theta modulation of principal cells (control 0.84 ± 0.06, CA3-APP 0.94 ± 0.05, p = 0.50, MW test, **Figure 8B**) or interneurons (control 0.15 ± 0.04, CA3-APP 0.09 ± 0.01, p = 0.48, MW test, **Figure 8D**). In CA3/DG recordings of aged CA3-APP mice we observed a strong reduction in the spike oscillation frequency of both principal cells (control 9.3 ± 0.1 Hz, CA3-APP 8.7 ± 0.1Hz, p < 0.0001, MW test, **Figure 8E**) and interneurons (control 9.0 ± 0.1 Hz, CA3-APP 8.0 ± 0.1Hz, p < 0.0001, MW test, **Figure 8G**), and similarly to CA1 recordings, there was a substantial increase in the fraction of cells with oscillations at ~7.5 Hz. The downward shift for principal cell theta oscillations in CA3-APP mice was pronounced, such that they were at a closer frequency to LFP theta oscillations than in controls (control 2.2 ± 0.1 Hz, CA3-APP 0.9 ± 0.1 Hz, p < 0.0001, MW test, **Supplementary Figure 16E**) and showed reduced theta modulation amplitude (control 0.87 ± 0.05, CA3-APP 0.61 ± 0.05, p < 0.0001, MW test, **Figure 8F**). Interneurons also showed a reduction in the difference to LFP theta oscillation frequency (control 0.6 ± 0.2 Hz, CA3-APP 0.3 ± 0.1 Hz, p = 0.028, MW test, **Supplementary Figure 16G**), but an increase in theta modulation amplitude (control 0.19 ± 0.04, CA3-APP 0.25 ± 0.05, p < 0.0001, MW test, **Figure 8H**). Strikingly, almost all interneurons in the CA3/DG region had a spike oscillation frequency at ~7.5 Hz, which was at or below the frequency of LFP theta oscillations.

**Figure 8.**
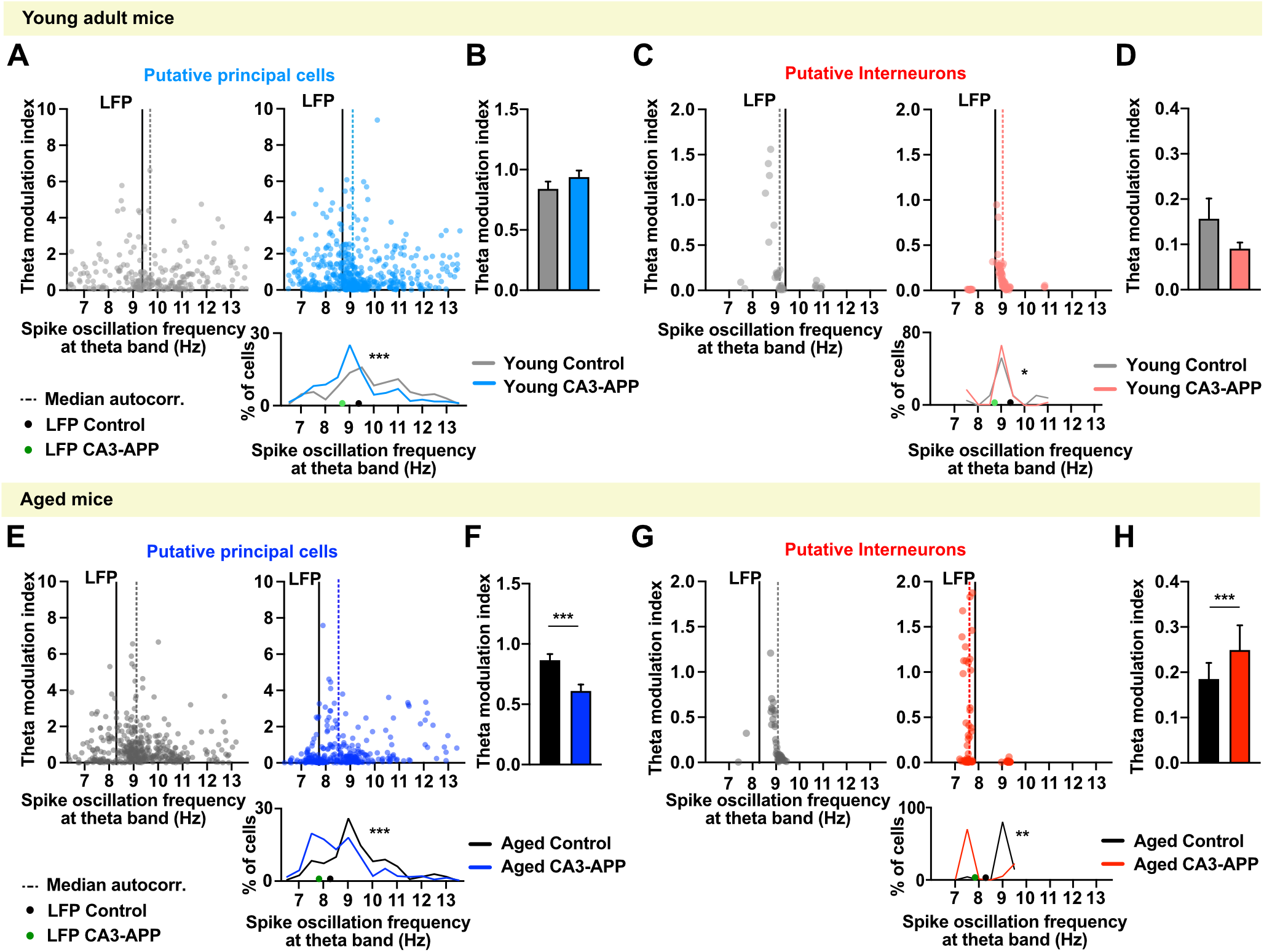
Spike oscillation frequencies of CA3/DG principal cells are reduced to a similar extent as for CA1 principal cells, but oscillation frequencies of interneurons in aged CA3-APP mice show a pronounced effect of hAPP expression. **A.** The distribution of the peak oscillation frequencies is plotted against the amplitude of their theta modulation (i.e., theta modulation index). Only cells with a detectable peak within the theta band are displayed, and the dashed line represents the median of their frequency distribution. The mean LFP frequency of the sessions in which these cells were recorded is indicated by the solid black line. Spike oscillation frequencies of CA3/DG principal cells were lower in young adult CA3-APP mice than in littermate controls (n = 226 cells for control, n = 500 for CA3-APP, p < 0.0001, MW test). **B.** Average theta modulation amplitude (theta modulation index) of all active CA3/DG principal cells. There was no difference between cells from young adult CA3-APP compared to cells from age-matched mice (n = 297 for control, n = 566 for CA3-APP, p = 0.50, MW test). **C.** In young adult CA3 APP mice, there was a minor reduction in the spike oscillation frequency of CA3/DG interneurons compared to age-matched controls (n = 36 cells for control, n = 87 for CA3-APP, p = 0.016, MW test). Data are displayed as in A. **D.** The average theta modulation index of all CA3/DG interneurons. Theta modulation amplitude of interneurons recorded from young adult CA3-APP mice did not differ from age-matched control mice (n = 59 for control, n = 111 for CA3-APP, p = 0.48, MW test). **E.** When compared to age-matched controls, principal cells from aged CA3-APP mice showed a major reduction in the spike oscillation frequency with a large fraction of cells oscillating between 7 and 8 Hz (n = 380 cells for control, n = 289 for CA3-APP, p < 0.0001, MW test). Data are displayed as in A. **F.** Theta modulation amplitude of CA3/DG principal cells of CA3-APP mice was reduced compared to age-matched controls (n = 425 for control, n = 316 for CA3-APP, p < 0.0001, MW test). **G.** Compared to littermate controls, CA3/DG interneurons from aged CA3-APP mice showed a pronounced reduction in spike oscillation frequencies, with almost all interneurons now oscillating below 8 Hz (n = 52 cells for control, n = 106 for CA3-APP, p < 0.0001, MW test). Data are displayed as in A. **H.** Interneurons from aged CA3-APP mice showed an increase in theta modulation amplitude compared to age-matched controls mice (n = 52 for control, n = 106 for CA3-APP, p < 0.0001, MW test). Data and statistics in **Supplementary Table 17**.

## Discussion

To examine whether highly localized expression of hAPP alters neuronal activity patterns in cell populations that receive projections from hAPP-expressing neurons, but themselves do not harbor hAPP, we generated mice that express hAPP in only hippocampal CA3 principal cells. With these mice, we performed memory testing and recorded from the hippocampal CA1 region during the memory task. Despite the restricted pattern of APP expression in only one hippocampal subregion, hippocampal-dependent memory was partially impaired in young adult CA3-APP mice and substantially impaired in aged CA3-APP mice, indicating that even without amyloid deposits and widespread pathology, hAPP can disrupt neural circuit function. In recordings while mice performed the behavioral task, CA1 principal cells of CA3-APP mice exhibited reduced firing rates during behavior, but despite these rate changes, hippocampal spatial coding remained largely intact, even at advanced ages (16-19 months). We rather found a striking number of changes in temporal firing patterns. First, theta and high gamma oscillation frequencies were decreased, but without changes in LFP amplitude. Second, subpopulations of principal cells and interneurons showed spike oscillation patterns at lower frequencies than in controls, along with a reduction of theta phase precession of place cells and impaired sequential firing of place cells during the task. While some of these deficits were already detectable in the CA1 region of young adult CA3-APP mice, the deterioration of temporal coding was more pronounced in aged mice and thus correlated with the progressive worsening of memory deficits. Physiological deficits were thus pronounced in a brain region that receives synaptic inputs from a cell population that expresses hAPP, but does not locally express APP. In particular, effects on the temporal and sequential organization of hippocampal patterns were pronounced, even without overt amyloid deposition, cell death, or deficit in place coding. Effects of hAPP on temporal coding are thus a possible cause for early deficits in spatial navigation associated with Alzheimer’s disease.

The predominant effect on temporal firing rather than on spatial properties of CA1 cells differs from previous studies in pan-neuronal models of amyloidosis that have linked impaired spatial properties of spatially modulated cells, such as place^6,11,29^ and grid cells^7,13^ with memory deficits. However, the majority of previous recordings were performed at advanced stages of amyloid pathology, when plaques, robust spine loss and sometimes cell death were already present. Similarly, major effects on place cells and grid cells were also observed in tau models, again at stages with substantial neurodegeneration^9,30^, which makes it challenging to infer whether memory deficits and spatial coding deficits arise as a direct effect of the primary pathology or more indirectly from loss of synapses and from cell death. Even in healthy mice, the extent of spatial coding deficits is not a good predictor for memory deficits, as shown in studies in which hippocampal-dependent memory deficits occur while the place code in CA1 is intact^31,32^. It has therefore been suggested that the generation of distinct hippocampal maps across different environments may be a better indicator for memory dysfunction in disease models^7^. While this may be relevant for memory tasks that require discrimination between spatial contexts, the previous recordings were not performed in a discrimination task. In contrast, we performed our recordings in a hippocampus-dependent memory task, and the task included a stem section where sensory inputs were corresponding such that differences in firing patterns directly relate to past or future turn direction ^33^. When restricting our analysis to the stem, the firing patterns of CA1 cells were as distinct between turn directions in CA3-APP mice as in control mice, which makes it unlikely that less distinct coding in the stem is the cause for the memory deficits.

Since we did not observe substantial deficits in spatial coding in CA1, which is the brain region that receives projections from APP-expressing CA3 cells but does itself not express hAPP, we also examined the possibility that the pronounced deficits that have previously been reported in AD models with pan-neuronal APP expression may emerge preferentially in brain regions with local hAPP expression. Importantly, the hippocampal CA3 region has strong recurrent connectivity, and circuit dysfunction in this region can therefore emerge from any combination of cellular APP expression and presynaptic APP-expression in inputs from other CA3 cells, which is the combination that would be typical for pan-neuronal models. We therefore examined whether we could detect more severe deficits in circuit function in the CA3 region compared to the CA1 region, which receives inputs from cells with hAPP but does not locally express hAPP. The effects of APP expression on hippocampal firing patterns were generally very similar between the CA3/DG region and the CA1 region in APP-CA3 mice. Some effects, such as those on high gamma oscillation frequency, were less pronounced in the CA3/DG region than in CA1, but this could be a consequence of greater prominence of high gamma oscillations in CA1 compared to CA3^34^. These findings suggest that local hAPP expression does not further exacerbate the deterioration of the cells’ firing patterns, and that in turn, synaptic inputs from hAPP-expressing cells are sufficient to result in circuit dysfunction in CA1. Our finding that synaptic inputs are sufficient to result in lower firing rates are consistent with the finding that dentate granule (DG) cells show lower levels of immediate-early gene expression when their input neurons, but not DG cells express APP^35^. However, here we did not show extensive dysfunction in the cells’ firing patterns in the target region, and in many hours of recordings from each mouse, there was no evidence of the emergence of epileptiform activity, as has been described with APP expression that was targeted to entorhinal cortex. A major difference between our CA3-APP model and the EC-APP model is that the latter is only partially regionally restricted. In EC-APP mice, the APP expression is also observed in cortical areas other than EC, and plaques are deposited at the site where epileptiform activity emerges^35^.

While many firing properties show a comparable or lesser effect in CA3/DG than in CA1, one effect that is more pronounced in the CA3/DG cell population is the decrease in the spike oscillation frequency of interneurons. According to anatomical studies, the interneurons in this region are mostly composed of basket cells (CCK or PV positive) and bistratified cells (SOM and PV positive)^36^, and all of these receive projections from CA3 principal cells, which express hAPP in the CA3-APP mice. However, similar to the interneurons in CA1, the CA3/DG interneurons themselves do not directly express APP. Our study is therefore the first to evaluate interneuron physiology in the context of AD without directly expressing AD-related proteins in interneurons^37–39^. The fact that a large fraction of interneurons in CA3-APP mice show major temporal reorganization, such as large shifts in oscillation frequency and lack of speed modulation, points to the importance of considering interneurons as major hubs for the manifestation of disease pathology. While a previous study pointed to the role of sodium channels that could be altered by either cellular or presynaptic APP expression^37^, our results highlight the importance of investigating how synapses between APP expressing afferent neurons and interneurons are altered in the context of Alzheimer’s disease^40–42^. In particular, we also find that a subset of principal neurons shows a decrease in spike oscillation frequency that corresponds to the shift in the frequency that is observed for interneurons. This suggests that feedback loops between excitatory and inhibitory cells could amplify the pathology, but with APP expression in principal neurons that are presynaptic to interneurons driving the changes.

As a consequence of the changes in oscillations and feedback loops in networks that include interneurons and principal cells, CA3-APP mice show a large reduction in theta frequency and in the associated high gamma oscillations, which are thought to emerge locally^43^. Recent studies have also described a reduction in theta frequency in pan-neuronal models of AD and that optogenetic stimulation at theta frequency has the potential to ameliorate hippocampal memory in these mice (J20 and APP/PS1)^39,44^. Our work adds to these studies by showing that a reduction in theta oscillation frequency can already emerge from a restricted pathology, which does not have to directly include interneurons. Our findings that effects on oscillations appear prior to other physiological changes and are present along with behavioral deficits, provides further support for using the strategy of theta pacing to rescue memory function. Furthermore, the more pronounced spatial navigation deficits in aged compared to young adult CA3-APP mice were accompanied by an impairment in sequential firing within theta cycles and by a reduction in phase precession in the version of the memory task that is hippocampus dependent (i.e., with 2-s and 10-s delay). While our findings of a disrupted temporal organization of hippocampal firing patterns might appear to be at odds with a previous report from a Tau AD model where place cells were largely impaired but the sequential activity of cells was maintained^9^, it is important to note that the Tau model is characterized by a loss of more than 50% of neurons in the CA1 pyramidal layer at the time of recordings. Therefore, the rigid firing sequences might represent an advanced disease state when compensatory mechanisms maintain highly stereotyped sequential firing of hippocampal cells. In contrast, our model would resemble early pathology when hippocampal firing becomes progressively disorganized and can no longer support the computations for spatial memory and navigation, which is characteristic for early AD patients. In particular, the restricted expression of APP in only one hippocampal subregion shows that early effects on circuits are primarily on the temporal organization of firing patterns and that interneurons are important in mediating these effects. During disease progression, other circuit elements are therefore not only affected by spreading molecular pathology, but dysfunction also emerges in the absence of primary pathology, such as observed here for the firing properties of interneurons and CA1 cells.

## Acknowledgements

We would like to thank Dr. M. Sabariego for discussions and behavioral protocol design and Dr. I. Zutshi for invaluable discussions and analysis Scripts. We also thank Dr. B. Boublil, G DeGuia, Tiffany L. Huynh and Ellen Lee for technical assistance. This work was funded by NIH grants R01 NS084324, R01 NS102915, R01 NS097772 to S.L. and R01 MH119179 to J.K.L.

## Experimental methods

### Approvals

All experimental procedures were conducted at the University of California, San Diego according to National Institutes of Health guidelines and were approved by the Institutional Animal Care and Use Committee.

### Animals subjects

CA3-APP mice were generated as previously described^14^. In brief, ROSA26-ZtTA transgenic mice (JAX Stock No: 012266^16^) were first crossed with tetO-APPswe/ind transgenic mice (MMRRC Stock No: 34846-JAX^17^ and the resulting ZtTA/tet-APP bi-transgenic mice were then crossed with Grik4-cre mice (JAX Stock No: 006474^15^). Triple transgenic male breeders were crossed to CD-1 female mice to obtain the experimental mice with mixed strain backgrounds (C57Bl6/J and CD-1). This crossing strategy ensured large litters, which were necessary to obtain the needed number of triple transgenic CA3-APP mice. Littermate mice lacking the tetO-APP transgene were used as control to account for possible leakage of the promoter. The number of animals analyzed is reported in each figure and summarized in **Supplementary Table 1**, along with their sex and exact genotype. Mice were kept on a 12-hour light-dark cycle (light on from 7AM to 7PM) and all the experimental data was recorded in the light phase to improve the chances that mice were quiet during the rest sessions before and after behavior testing. All mice were group housed (maximum 5 per cage) with food and water *ad libitum* until the beginning of behavioral training at 4 or 16 months, and from then they were single housed and food restricted.

### Behavior equipment

Mice were trained on a figure-8 shaped maze (**Figure 1C**) 50 cm by 75 cm long. The maze runways were made of dark grey Plexiglas 5 cm wide with a 0.5 cm high border. The figure-8 maze was elevated 55 cm above the floor, positioned in the center of a room where a bench, an acquisition system and a dim light source served as prominent visual cues. Two barriers made of vinyl-covered cardboard were used to confine the animals in the initial 20 cm of the central runway (delay area, **Figure 3A**). Sugar pellets were attached bellow the runaways at the reward sites to provide permanent odor cues. During the rest periods mice were in their home cage inside a taller transparent plexiglass box that was placed on the bench, 1.2 cm from the floor.

### Behavior training and testing

One week before the start of behavioral training transgenic mice and littermate controls were single housed and food restricted to 85-90% of their baseline body weight. Nesting materials and light enrichment was provided. Mice were handled for 5-10 minutes a day during a *habituation phase*, where they were also allowed to forage for sugar pellets (20mg; Test Diet) scattered in the figure-8 maze. Once mice consistently ate the scattered sugar pellets, behavior training was initiated. First, mice were guided and trained to run in one direction and to alternate between left and right turns to find reward at the end of the right and left arms (reward area). The experimenter was always on the side where the rewards were located to refill. Mice ran one session of 20 minutes a day, and after 2 consecutive days of consistently eating the reward pellets and running without turning back, mice advanced to the next phase of behavior training during which all barriers and guidance were removed. During this phase mice ran 20 minutes or 60 laps receiving a reward only for correct (alternating) turns. Once the criterium of >80% correct turns for 4 out of 5 consecutive days was reached, mice advanced to the testing phase, where a delay period was introduced in the central arm of the figure-8 maze.

The testing phase consisted of 5 consecutive days when mice ran 60 laps a day separated into 6 blocks with different delays periods (no delay, 2 s delay and 10 s delay). The order at which the different delay blocks were presented was pseudorandomized to ensure that each condition had the same levels of motivation/hunger. All mice followed the same order. After the behavior testing mice were allowed 5-10 days with *ad libitum* food before a microdrive implant surgery was performed. Mice were then allowed 5 days of post-op recovery before food restriction resumed. While tetrodes were lowered into the hippocampus (between 6-15 days) mice were trained to run in the figure-8 maze with elastics supporting the tether, and a maximum of 30 laps without any delay were performed per day. Once the tetrodes reached the hippocampus, daily recordings with 60 laps and with the different delay conditions commenced. At least 10 days of behavioral data was obtained from with the same design as for recording days, even if the microdrive implant failed to produce data (due to a failure of the microdrive to advance, a broken ground, or unstable recordings).

### Electrode implant surgery

Tetrodes were prepared by twisting 4 insulated platinum wires (0.017 mm of diameter, California Fine Wire Company) and melting the insulation to bind the four electrodes together. The electrodes were loaded into a 16-channel microdrive^45^ and plated with platinum to achieve stable impedances near 200 MΩ. The microdrive was fitted with an optic fiber attached to the tetrode bundle (200 μm diameter optic fiber, Thor Labs) which provided stability and guidance. Mice were anesthetized with 1.5-2% isoflurane vaporized in oxygen in a stereotaxic apparatus. Analgesia was provided by a subcutaneous lidocaine (1%) injection in the animal’s head and an intraperitoneal injection of buprenorphine (0.02 mg/kg). Four or 5 small stainless steel anchor screws were positioned in the skull of the animal. A ground screw was positioned touching the left hemisphere cortex (AP −0.7; ML −2.6) and a small circular craniotomy was performed above the hippocampus on the right hemisphere (AP −1.9; ML +1.8). The tetrode tips were slowly inserted 0.4 mm beyond the dorsal surface of the brain. The exposed tetrode wires were protected by a gel (Na-Alginate cured by CaCl2) and a guide cannula, and the microdrive was secured with several layers of dental acrylic to the skull and the anchor screws. A subset of aged mice was implanted with 16-channel silicon probes (Neuronexus A1×16-3mm-50-177-CM16LP). In this case, the probe was directly lowered into the CA1 region of the hippocampus (−1.7 mm from the surface of the brain). A ground wire was positioned on the surface of the cortex in the right hemisphere and a reference wire was positioned in the left hemisphere cortex (AP-0.7; ML −2.6, same place as the ground screw for the microdrives). The reference/ground wires and the probe where then secured in place with dental acrylic. Animals were allowed to recover from surgery in their home cage over a heating blanket until awake.

### Electrophysiology recordings

Tetrode advancement started 5 days after the surgery and accrued to a maximum of 0.25 mm per day to retain stability. Recordings were performed only when tetrodes reached hippocampal principal layers (CA1, CA3 and DG) and well isolated clusters could be observed. Depending on the stability of clusters, tetrodes were advanced at the end of the day to ensure that a different set of cells was recorded the following day. The recordings stopped when all tetrodes had crossed the hippocampal layers or when the end of the drive range was reached (~3 mm). The implanted microdrive was connected to a digital Neuralynx recording system (Neuralynx, Bozeman, MT) through a multichannel, headstage preamplifier. The headstage and multi-wire tether were supported by a system of elastics to counteract their weight. Unit activity was amplified (5,000-20,000 times) and band-pass filtered (0.6-6 kHz) for spike sorting. Spike waveforms above a threshold of 35-50 μV were time-stamped and digitized at 32 kHz for 1 ms. Continuous LFP in the 0.1-900 Hz band was recorded from one of the wires from each tetrode at a sampling rate of 32 kHz. The x-y position of the mouse was tracked at 30 Hz by a video camera mounted above the experimental area. The camera detected a red and a green LED located on either side of the headstage. A maximum of 2 recording sessions were performed per day and they were separated by at least 2 h. The number of sessions completed per mouse varied between 4 to 20, depending on (1) the stability of the drive and (2) when tetrodes reached the layer (tetrodes were not independently movable). While we avoided double-counting cells by advancing tetrodes every day, in some cases the same cells could have been included in more than one analysis day.

### Histology and tetrode locations

At the end of the recording procedures, mice were anesthetized with isoflurane (4%) and given an overdose of sodium pentobarbital (>30 mg/kg) intraperitoneally. They were then perfused transcardially with Ringer’s solution (135 mM NaCl, 5.4 mM KCl, 1.8 mM CaCl^2^, 1 mM MgCl^2^, 5 mM HEPES, pH 7.4) followed by 4 % paraformaldehyde (PFA, Affymetrix USB). Brains were removed from the skull and post-fixed overnight in 4 % PFA at 4 °C. For a subset of mice (part of the aged group) the brain was extracted after the perfusion with cold Ringer’s solution and the right hemisphere was then fixed for 48-60h in 4% PFA. All brains (or hemibrains) were transferred to a 30% sucrose solution for cryopreservation 48h before sectioning. Coronal sections (40 μm) from the entire dorsal hippocampus were cut with a freezing microtome. Half of the sections were mounted and stained with cresyl violet to identify the small lesions caused by the electrodes and these locations where then mapped onto the stereotaxic atlas of the mouse brain. The remaining sections were saved for immunohistochemistry.

### Immunohistochemistry

For sections used for immunohistochemistry, slices were washed and then incubated for 20 min in 70% acetic acid for antigen retrieval. The slices were washed 3 times with water and blocked for 1 h at room temperature (RT) with a blocking solution comprising 10% horse serum (HS), 0.2% bovine serum albumin (BSA), and 0.05% Triton-X in 1x PBS. Slices were incubated overnight at 4 °C with primary antibody (Biotin anti-β-Amyloid, 1-16 Antibody, clone 6E10 from BioLegend) dissolved in a carrier solution made with 1% HS, 0.2% BSA, and 0.05% Triton-X in 1x PBS at a concentration of 1:500. After overnight incubation and 4 washes for 10 min each with 1x PBS, the slices were incubated for at least 2 h at RT with secondary Streptavidin, Alexa Fluor™ 488 conjugate, from ThermoFisher at a concentration of 1:1000. Next, the slices were washed four times with 1x PBS, mounted on glass slides, and coverslipped with DAPI containing Prolong Diamond Antifade medium™. Complete sections were imaged using a virtual slide microscope (Olympus, VS120), and individual planes of higher magnification images for quantification were obtained from a confocal microscope using 40x magnification lenses (Olympus FluoView, FV1000).

### Spike sorting and cluster quality

A customized version of the spike sorting software MClust, MATLAB 2009b (Redish, A.D. Mclust. https://redishlab.umn.edu/mclust) was used for spike sorting. Clustering was performed manually in two-dimensional projections of the parameter space using waveform amplitude, waveform energy, and the peak-to-valley difference. For a given cluster unit, the same boundaries were used across the rest and behavioral task segments of a recording session. Single units were excluded from analysis if they were not separable from either the noise or other single units. To ensure stability all units included in the study were required to have spikes in both the pre and post-behavioral resting sessions, while they were not required to have spikes during the behavioral blocks. Cluster quality was assessed by calculating the L-ratio and isolation distance^46^ (**Supplementary Figure 2B**) for each cluster during behavior. The L-ratio is calculated by dividing L by the number of spikes within the cluster, where L is calculated from the sum of 1 minus the Chi-square cumulative distribution function of the Mahalanobis distance of the spikes outside the cluster.

### Unit classification

Isolated units were classified using a set of criteria applied to their average firing rate and peak-valley ratio of their spike waveform (**Supplementary Figure 2C**) calculated from the entire session (behavior, pre- and post-resting periods). The criteria definitions were chosen to accommodate possible changes in rate or spike properties that could occur as result of our genetic manipulation. Putative principal cells were defined as units with (i) a peak-valley ratio above 0.9, (ii) an average firing rate below a threshold defined by a slope [0.4*mean firing rate with a offset of -(peak-valley ratio + 1)] that allowed higher average firing rates for cells with higher peak-valley ratio, and (iii) a maximum mean firing rate of 10 Hz.

The classification of putative principal cells was the same for recordings in CA1 and CA3/DG units. The vast majority of putative interneurons in the CA1 region were recorded within *stratum pyramidale*, whereas recordings in the CA3/DG regions include some units recorded from the hilar region. Therefore, putative interneurons were defined separately for CA1 and CA3/DG recordings, in order to include the more diverse classes of interneurons in the latter regions. Putative interneurons in CA1 were defined by following criteria: (i) a firing rate of 5-60Hz, (ii) a peak-valley ratio below 2, and (iii) a ratio of firing rate to peak-valley ratio above a slope [Slope 2: 0.1*mean firing rate with an offset of -(peak-valley ratio - 0.5)] which includes interneurons with lower firing rates (min. 5 Hz) if they have a low peak-valley ratio, a known feature of interneurons. Units recorded in CA3/DG were classified as putative interneurons if they (i) were not classified as principal cells, (ii) had a firing rate of 5-60 Hz, and (iii) had a maximum peak-valley ratio of 2. Units that did not fulfill the inclusion criteria were excluded from further analysis (**Supplementary Figure 2C**).

### LFP analysis

#### Average spectrograms

Local field potentials (LFPs) obtained from the hippocampus were referenced to a ground screw in the case of microdrives recordings or to the reference wire for the silicon probe recordings (both were located at the same coordinate). For each recording session the selection criteria for the tetrode for LFP analysis was that it had (1) clearly defined clusters and (2) the highest amplitude of sharp-wave ripples (SWR). For the recordings performed with the silicon probes, the selected channel was the one immediately below the channel registering the highest SWR power.

Raw LFP signals were downsampled to 2 kHz and parsed by the region where the animal was localized during behavioral sessions. The spectrogram was calculated for frequencies of 2-300 Hz using a Morlet wavelet transformation (MATLAB package) ^47^. Power spectral density (PSD) was calculated for each maze section, and the most prominent peak frequencies were defined using the average PSD from all correct-turn trials of one behavioral session per day within the following ranges: delta 2-6 Hz, theta 6-12 Hz, beta 14-26 Hz, low gamma 26-50 Hz, high gamma 50-150 Hz. Power within each frequency band was defined as the power at the peak frequency. Calculated peak frequencies and powers were averaged per animal from different recording days. We could not consistently detect peaks in the power spectrum density at higher frequencies in the raw signal of many animals (**Supplementary Figures 7 and 11** and **Supplementary Tables 10 and 14**).

For better visualization, the parsed spectrograms were resized to match the average time duration of all the animals spent in this region, in order to allow averaging across trials of different duration. Normalized spectrograms were calculated by dividing spectrograms by the PSD of the resting period (r1) on the same recording day, so that normalized spectrograms for the behavioral task reflect amplitudes as multiples of the amplitude during rest. This normalization was considered more rigorous than a Z-score or a 1/f correction as it (i) is independent of the activity of the mice during the delay and allows for a much better comparison between mice (as some quietly wait for the barrier to be lifted while others move between the two barriers), (ii) provides an internal baseline that does not depend on the exact location of the recording or how good we can fit an exponential to the signal, and (iii) normalizes equally between high and low frequencies (the variation of signal standard deviations is higher at lower frequencies)

### Firing Properties of Single Units

#### Average firing rate

was calculated from the total number of spikes that occurred during the entire session (three delay conditions and two resting sessions). No speed threshold was applied. *Speed score* was calculated as following. Instantaneous firing rate was calculated for spikes that occurred when the animal was running at a velocity above 2 cm/s. Pearson’s correlation between instantaneous firing rate and instantaneous velocity was then calculated to obtain the speed score of each cell^20^. The average of the three delay conditions was calculated weighted by the number of spikes of each session (d0, d2, d10). *Active cells* were defined as cells with a firing rate above 0.1 Hz during behavior (approximately 60 spikes per ~10 minute-session). *Bursting index* was defined as the total number of spikes occurring in bursts (inter spike interval <6 ms) divided by the total number of spikes^48^.

### Place Cell Analysis

The path of the animal in the figure-8 maze was linearized and binned into 76 spatial bins of 2.5 cm x 2.5 cm. A lap started and ended at the reward location. Right and left starting laps were analyzed separately and correct and incorrect laps were analyzed separately. Because control mice have very few incorrect laps, only correct laps recorded from the 10-s delay condition (d10) were used. A binned rate map was calculated for each cell dividing spikes per time spent in each bin, and smoothed with a one-dimensional version of the boxcar filter. *Place cells* were defined as cells with (i) a peak firing rate of ≥3 Hz in at least one spatial bin and (ii) at least one place field between 12.5 cm and 70 cm. The *place field(s)* of a cell were defined as the space before and after the peak in which the firing rate remained above 20% of the peak firing rate. The start and end points (20% of the peak value) of the place field were determined on up-sampled (linear interpolated) data. To determine the *number of place fields per cell* and to avoid counting peaks in the central arm (where left and right turns overlap) twice, place fields occurring in the central arm (delay + stem) with an overlap above 25% of the place field dimensions were removed from the turning side with the smaller place field peak firing rate. The *peak firing rate* of a place cell was defined as the spatial bin with the highest rate value. *Spatial information*, defined as the information density per spike (I), was calculated as previously described^27^ using the average rate and occupancy maps for left and right laps. The following formula was used:

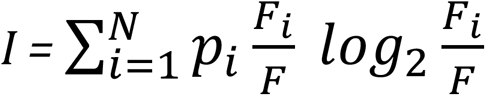

where *p_i_* is the probability of occupancy in bin *i, F_i_* is the mean firing rate for bin *i* and *F* is the mean firing rate.

### Place cell spatial correlations

All spatial correlations were calculated from the linearized rate maps of correct laps during the 10-s delay condition (d10). *Trial by trial correlation* as a measure of stability was calculated as a Pearson’s cross-correlation of the firing rates between every combination of trials in the same direction. Spatial bins corresponding to the delay area were removed. To account for cells with a place field only in one side of the maze, the correlation was calculated separately for left-turn and right-turn laps, and maximum correlation value was used. The *correlation across sides* was calculated as the average Pearson’s cross-correlation between the rate maps of left-turn and right-turn laps. Only bins corresponding to the non-overlapping return arms were used. *Splitter cells* were defined as place cells where the following criteria were fulfilled for at least two consecutive spatial bins (5 cm): (i) the firing rate was at least 5 Hz and (ii) at least twice as high as for the opposite turn direction. This analysis excluded the choice region, where the animal’s path already diverged, as they might have a very different set of visual input depending if they are turning left or right. The splitter acuity/selectivity index was calculated from the inverse of a Pearson’s correlation between the average rate maps of left-turn and right-turn laps along the delay and the central arm [S = (1/(r+1))-0.5], where a larger value indicates stronger distinction between left and right turns.

### Temporal properties of single units

To determine the theta modulation (spike oscillation frequency and theta modulation strength) of each unit, the Fourier transform of the autocorrelation of all spikes was calculated (time resolution: 5ms) and filtered in the theta band from 6 to 14 Hz. Based on spike and LFP timestamps, the theta phase at the time of every spike were interpolated. Using circular statistics, we calculated the mean vector length and the mean phase of the spikes of each cell and the significance of theta modulation.

### Phase precession and sequence compression

For each place cell, phase precession was calculated using bursts of activity (spike interval <500ms and including at least 5 spikes). The slopes of each spike train (significant or not) were then used to obtain the median value of phase precession. For the sequence compression analysis, 2D place fields were defined in the maze (2.5 cm by 2.5 cm bins were included if the rate in the bin was at least 0.3 Hz, if there were a minimum number of 8 bins above threshold, and if at least one bin had a peak rate of >3 Hz). Minimum overlap between fields was 15%, and place fields overlapping across the delay area were excluded, as we could not establish the earliest place field. Spatial distance between place fields was the linear distance between the centers of mass of two place fields. Crosscorrelograms were generated for all overlapping cell pairs (bin size: 7.5 ms, from −3 s to 3 s). The *real time* was defined as the highest peak in the crosscorrelogram, while the theta time was defined as the highest peak within the −100 ms to +100 ms interval. Correlations between distance, real time and theta time were performed using Pearson’s correlation tests.

## Supplementary Material

for *Viana da Silva et al.*

## List of Abbreviations

avg: average
ctrl: control mice/cells
tg: CA3-APP mice/cells
stat. metrics: statistical metrics
df: degrees of freedom
sig: significance
suppl: supplementary
KS: Kolmogorov-Smirnov statistical test
MW: Mann-Whitney statistical test
RM: repeated measurements
rest sess: rest session
ns: non significant
INT: interneurons
PYR: pyramidal cell
d0, d2, d10: recording sessions in the maze with either no delay, a 2-s delay or a 10-s delay in the central arm
corr: correlation
rec: recording
dist: distribution

---

**Supplementary Table 1.** Genotype and sex of experimental mice.

| Experimental Group | Genotype | Number of mice |
| --- | --- | --- |
| **Young adult mice** |  |  |
| **CA3-APP mice** | C+ L+ A+ | **16 (9 males, 7 females)** |
| **Control** |  | **17 (10 males, 7 females)** |
|  | C- L- A- | 7 |
|  | C+ L- A- | 6 |
|  | C+ L+ A- | 3 |
|  | C- L+ A- | 1 |
| **Aged mice** |  |  |
| **CA3-APP mice** | C+ L+ A+ | **20 (12 males, 8 females)** |
| **Control** |  | **20 (12 males, 8 females)** |
|  | C- L- A- | 7 |
|  | C+ L- A- | 6 |
|  | C+ L+ A- | 5 |
|  | C- L+ A- | 2 |

Transgenes:
C, Grik-Cre transgene
L, LacZ-tTA transgene
A, TRE-APP transgene

**Supplementary Figure 1.**
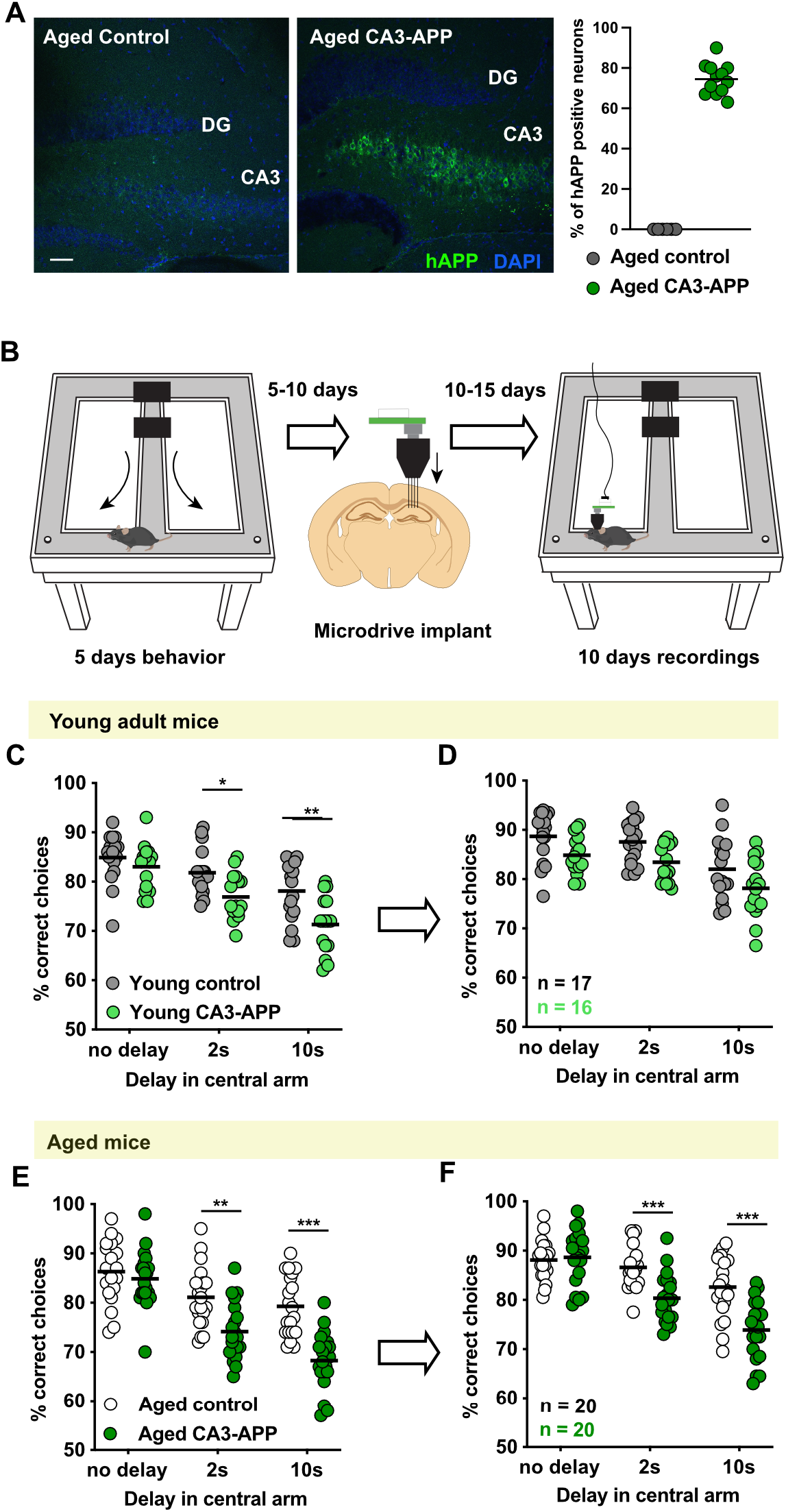
Young adult CA3-APP mice were transiently impaired, and aged CA3-APP mice were persistently impaired in a hippocampus-dependent memory task. **A.** Quantification of the number of APP expressing cells in aged mice (from two slices per mouse, n = 6 mice per genotype). In CA3-APP mice, 74.5 ± 2.2% of CA3 cells were hAPP positive. No hAPP-positive cells were detected in other hippocampal or cortical areas, or in any brain region in control mice. Scale bar is 100 μm. **B.** Timeline of the experimental design. After the initial behavioral training in the figure-8 maze without delay, mice underwent 5 days of behavioral testing in the spatial alternation task with no delay, 2-s delay, and 10-s delay. A microdrive was implanted 5 to 10 days after behavior testing, and tetrodes were slowly advanced towards hippocampal cell layers. Recordings commenced once hippocampal cellular activity was detected, and behavioral testing with recordings was performed for 10 days. **C.** Preoperative performance of young adult CA3-APP mice and control littermates in the figure-8 maze spatial alternation task. The performance of young adult CA3-APP mice (n = 16) did not differ from young adult control mice (n = 17) in the continuous version of the task, but the percentage of correct choices was lower in CA3-APP mice than in control mice when a 2-s delay or 10-s delay was introduced (p = 0.65 for no delay, p = 0.020 for 2-s delay, and p = 0.0007 for 10-s delay, Sidak’s post-hoc test). Each dot is an individual’s average percent correct trials over 5 days in a delay condition. Horizontal lines correspond to medians. See **Supplementary Table 2** for statistics. **D.** Performance of young adult CA3-APP mice in the spatial alternation task during the recording phase. No differences between genotypes were found (p = 0.083 for no delay, p = 0.058 for 2-s delay, and p = 0.079 for 10-s delay, Sidak’s post-hoc test). Each dot is an individual’s average percent correct trials over 10 days in a delay condition. Horizontal lines correspond to medians. See **Supplementary Table 2** for statistics. **E.** The preoperative performance of CA3-APP mice (n = 20) and control littermates (n = 20) did not differ when there was no delay. The introduction of a delay in the central arm resulted in decreased performance of aged CA3-APP compared to aged control mice (p = 0.83 for no delay, p = 0.001 for 2-s delay, and p < 0.0001 for 10-s delay, Sidak’s post-hoc test). Data are displayed as in C. See **Supplementary Table 2** for statistics. **F.** The performance of CA3-APP aged mice during the recording phase was reduced when a 2-s delay and when a 10-s delay were introduced in the central arm (p = 0.98 for no delay, p = 0.0008, for 2-s delay, and p < 0.0001 for 10-s delay, Sidak’s post-hoc test). Data are displayed as in D. See **Supplementary Table 2** for statistics. * p < 0.05, ** p < 0.01, *** p < 0.001.

**Supplementary Table 2.**
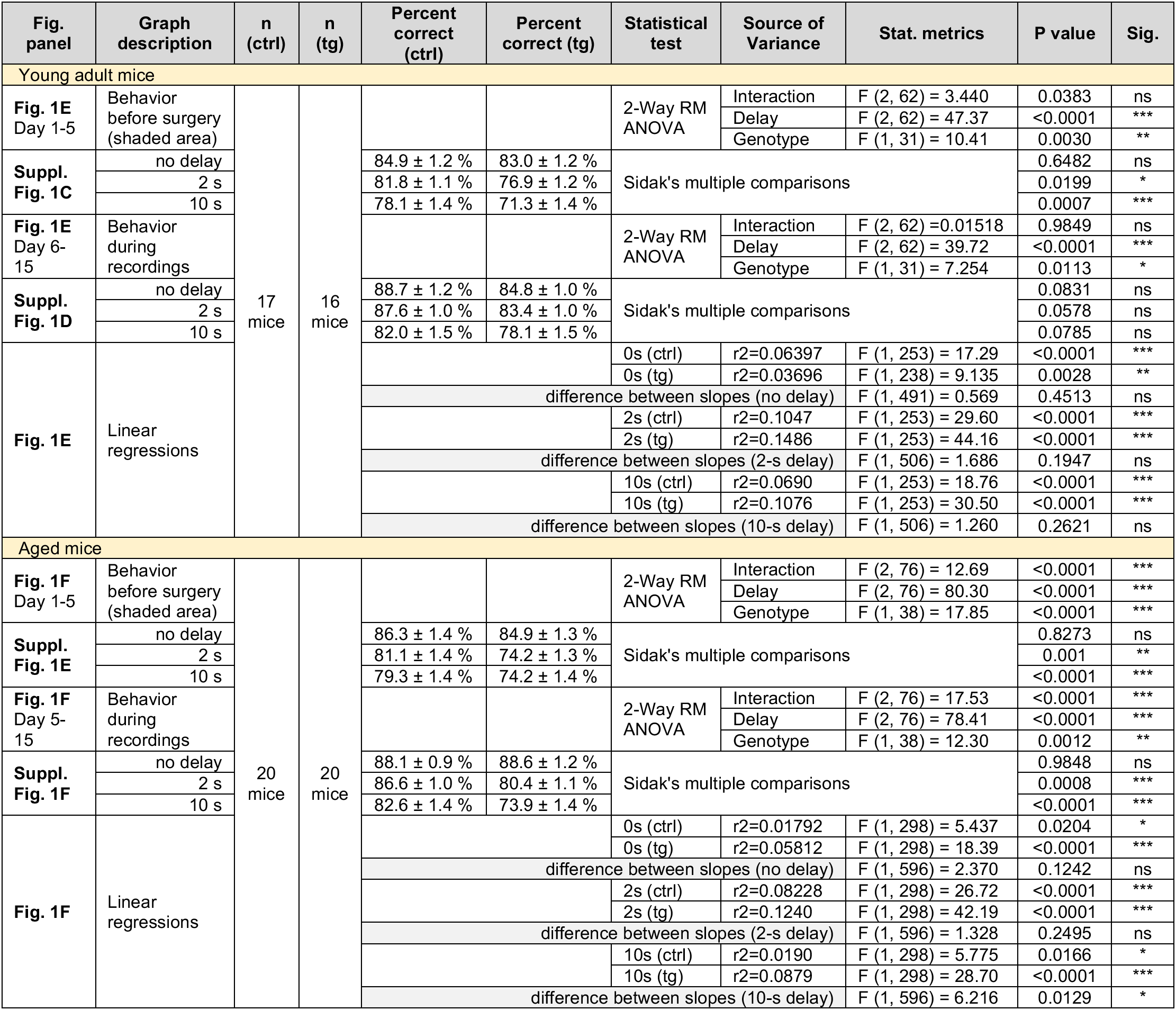
Behavior data and statistics, related to Figure 1 and Supplementary Figure 1.

**Supplementary Figure 2.**
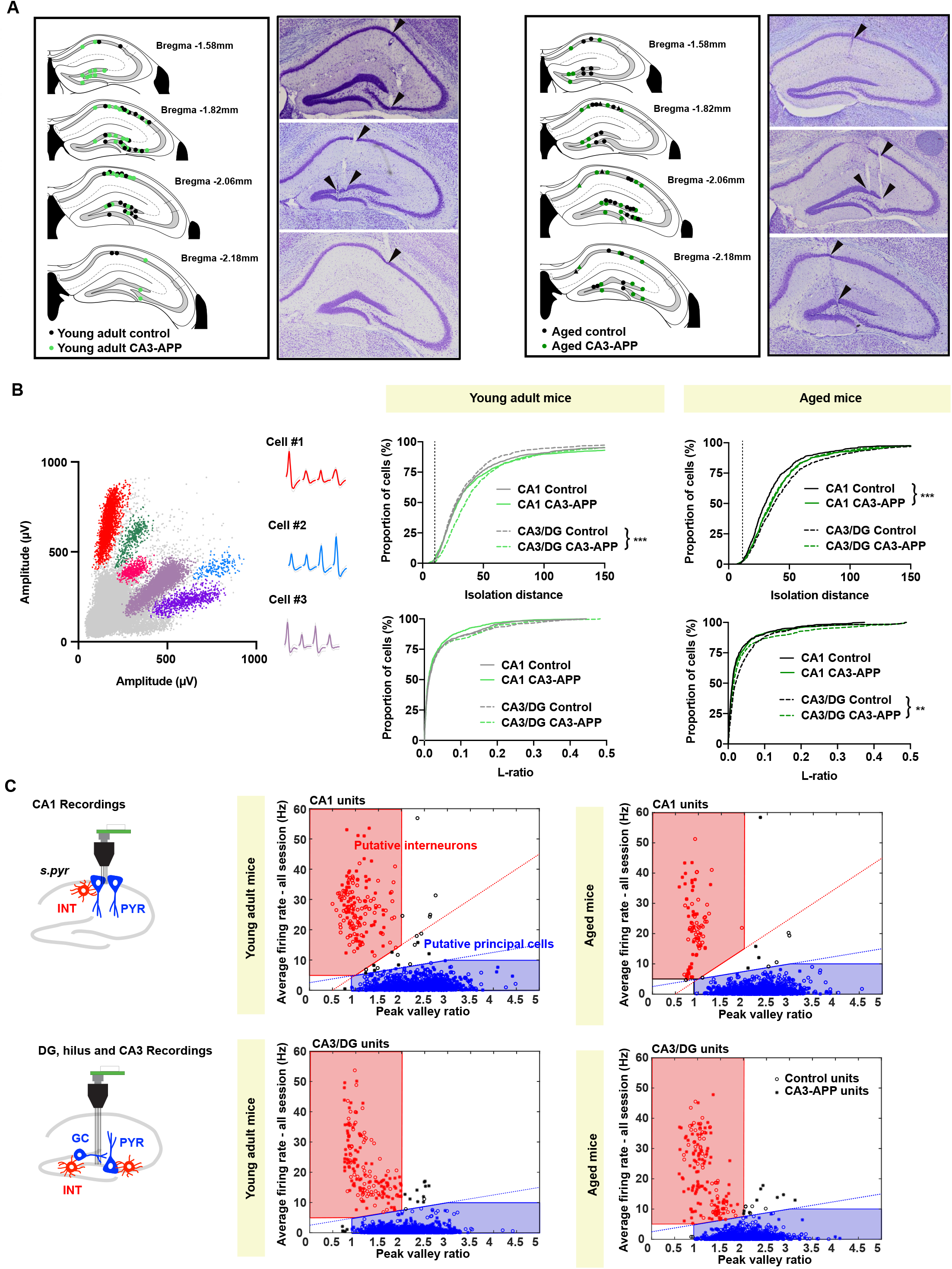
Recordings were performed in the hippocampal CA1 area or in the CA3/hilar/DG region, and putative principal cells and interneurons were distinguished. **A.** Summary of all tetrode positions reconstructed from histological material. Because electrodes were closely spaced within animals, some dots represent more than one tetrode from the same mouse. Examples of cresyl violet stained sections from 3 different young adult mice and 3 different aged mice. Positions of recordings performed with tetrodes are depicted as dots, and positions of recordings performed with silicon probes in aged mice are depicted as triangles. **B.** Example clusters from an aged CA3-APP mouse, and measurements of cluster quality. Upper panels show isolation distance and lower panels L-ratio. Although there were minor differences between control and CA3-APP cells, more than 99% of clusters had an isolation distance above a commonly used cut-off of 10 (Schmitzer-Torbert *et al.* 2005)^46^. **C.** Single-unit classification was based on firing rate and on peak-valley ratio. Panels display the distribution of clusters recorded from control mice (circles) and CA3-APP mice (stars). The criteria used to define putative principal cells and interneurons are indicated by the blue and red boxes. Definitions for interneurons differed between CA1 and CA3/DG to include putative interneurons with intermediate firing rates (~10 Hz) that were frequently observed in CA3/DG. Units outside of the defined criteria were excluded from further analysis.

**Supplementary Figure 3.**
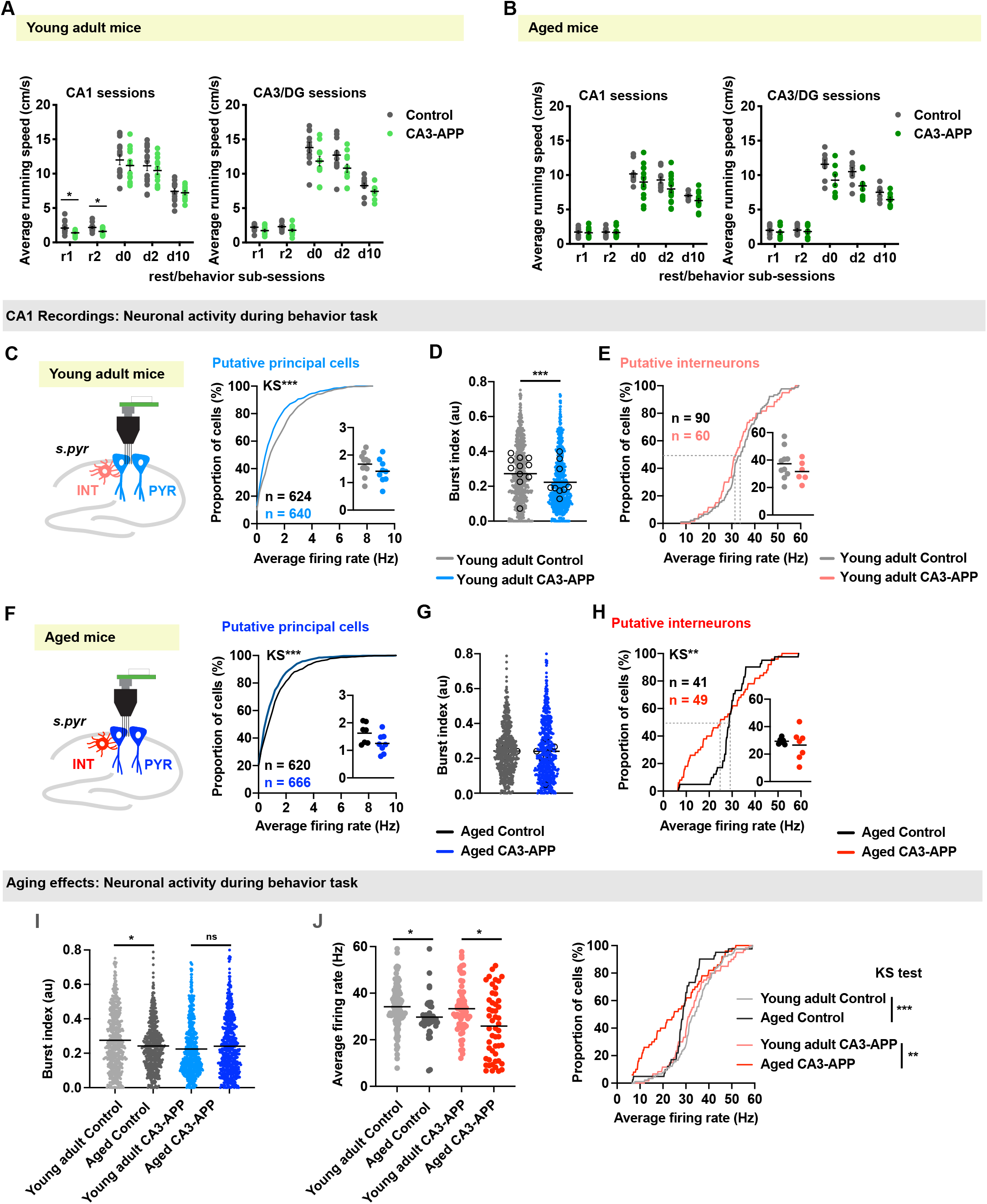
Behavioral patterns were similar between CA3-APP and control mice, but firing patterns of hippocampal CA1 cells differed. **A.** Average running speed of mice during different sub-sessions of the behavioral task: rest before and after running in the figure-8 maze (r1 and r2, respectively) and while running in the figure-8 maze with different delays in the central arm (d0, d2 and d10 respectively). Sessions when electrodes were located in CA1 or in CA3/DG were analyzed separately. The velocity of young adult CA3-APP mice compared to young adult control mice during CA1 recordings was slightly lower during rest sub-sessions, but did not differ during figure-8 maze sub-sessions (**Supplementary Table 4**). Each dot is the average for one mouse. **B.** There were no differences in average velocity between aged CA3-APP and aged control mice in rest or in run subsessions. Data are displayed as in A. **C.** Distribution of average firing rates of putative principal cells during running in the figure-8 maze (d0, d2, d10). The firing rates of principal cells were lower in young adult CA3-APP mice than in young adult control mice (control 1.6 ± 0.1 Hz, CA3-APP 1.2 ± 0.1 Hz, p < 0.0001, MW test). Inset shows data pooled by mouse (n = 11 for control and n = 9 for CA3-APP mice). **D.** Burst index of principal cells during running in the maze. Dots depict values of individual cells, horizontal lines the mean value of all cells pooled, and open circles the mean value per mouse. Principal cells from young adult CA3-APP mice have a reduced burst index compared to cells from young adult control mice (control 0.28 ± 0.01, CA3-APP 0.23 ± 0.01, p < 0.0001, MW test). **E.** Distribution of average firing rates of putative interneurons during running in the maze. Inset shows data pooled by mouse (n = 10 for control and n = 6 for CA3-APP). Average rates did not differ between young adult control and CA3-APP mice. **F.** Similar to the result in young adult mice, the average firing rate of CA1 principal cells in the maze was lower in aged CA3-APP mice (n = 8) compared to aged control mice (n = 7) (control 1.3 ± 0.1 Hz, CA3-APP 1.0 ± 0.04 Hz, p = 0.0007, MW test). Data are displayed as in C. **G.** The burst index of CA1 principal cells did not differ between aged CA3-APP mice and their control littermates, which differs from the reduction that we observed in young adult mice (control 0.24 ± 0.01, CA3-APP 0.24 ± 0.01, p = 0.21, MW test). The lack of a difference between the aged groups can be explained by a reduced burst index of control cells in aged mice (see below for a direct comparison between age groups). Data are displayed as in D. **H.** While there is no difference in the mean firing rate of CA1 interneurons from aged CA3-APP mice (n = 7) compared to aged control mice (n = 6, control 29.7 ± 1.4 Hz, CA3-APP 25.9 ± 2.0 Hz, p = 0.18, MW test), there is a statistical difference in the distribution (p = 0.0002, KS test). Data are displayed as in E. **I.** Effect of age on the burst index of principal cells. Bursting was reduced in aged control mice compared to young adult control mice (Kruskal-Wallis test with Dunn’s multiple comparisons test p = 0.038), and there was no further reduction in aged CA3-APP mice compared to aged control mice (Kruskal-Wallis test with Dunn’s multiple comparisons test p = 0.21; see G). **J.** Effect of age on average firing rate of interneurons. The firing rate in aged controls was lower than in young adult controls (Kruskal-Wallis test with Dunn’s multiple comparisons test p = 0.011), and the decrease with age was also observed for the comparison between the young adult and aged CA3-APP groups (Kruskal-Wallis test with Dunn’s multiple comparisons test p = 0.012). See **Supplementary Table 5** for all data and statistics.

**Supplementary Table 4.**
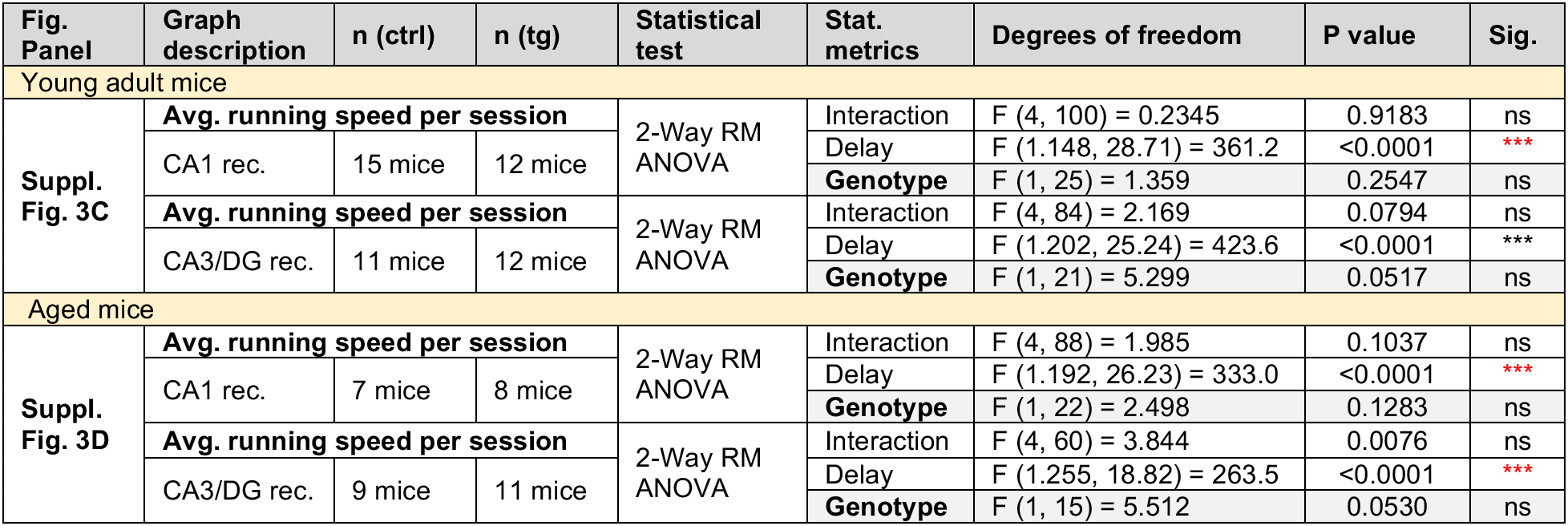
Average running speed in recording sub-sessions. Data and statistics for Supplementary Figures 3A and B.

**Supplementary Table 5.** Firing properties of CA1 single units, related to Figure 2 and Supplementary Figure 3.

| Fig. panel | Graph description | n (ctrl) | n (tg) | Normal Dist? | ctrl mean ± SEM | tg mean ± SEM | Statistical test | Stat. metrics | Df | P value | Sig |
| --- | --- | --- | --- | --- | --- | --- | --- | --- | --- | --- | --- |
| Young adult mice |  |  |  |  |  |  |  |  |  |  |  |
| CA1 pyramidal cells |  |  |  |  |  |  |  |  |  |  |  |
| Fig. 2B | Avg. firing rate in entire session | 629 | 645 | no/no | 1.6 ± 0.1 Hz | 1.4 ± 0.1 Hz | MW | U=184347 | | 0.0048 | ** |
|  | Distribution all cells - control vs CA3-APP |  |  |  |  |  | KS | D=0.08868 |  | 0.0134 | * |
| | Pooled per mouse | 11 | 9 | yes/yes | 1.6 ± 0.1 Hz | 1.4 ± 0.1 Hz | two sample t test | t=0.7778 | df=18 | 0.4468 | ns |
| Fig. 2C | % active cells CA1 | 11 | 9 | yes/yes | 90.7 ± 2.6 % | 90.7 ± 3.2 % | two sample t test | t=0.003526 | df=18 | 0.9972 | ns |
| CA1 interneurons |  |  |  |  |  |  |  |  |  |  |  |
| Fig. 2D | Avg. firing rate in entire session | 90 | 65 | yes/yes | 26.3 ± 0.7 Hz | 28.6 ± 1.2 Hz | two sample t test | t=1.742 | df=153 | 0.0835 | ns |
|  | Distribution all cells – control vs CA3-APP |  |  |  |  |  | KS | D=0.1897 |  | 0.132 | ns |
| | Pooled per mouse | 10 | 6 | yes/yes | 28.3 ± 2.0 Hz | 27.0 ± 2.9 Hz | two sample t test | t=0.3771 | df=14 | 0.7117 | ns |
| Fig. 2E | Speed modulation index | 90 | 60 | yes/yes | 0.15 ± 0.01 | 0.11 ± 0.01 | two sample t test | t=5.847 | df=148 | <0.0001 | *** |
|  | control different from 0? |  |  |  |  |  | Wilcoxon signed rank | W=3833 |  | <0.0001 | *** |
|  | CA3-APP different from 0? |  |  |  |  |  | Wilcoxon signed rank | W=1226 |  | <0.0001 | *** |
| | Pooled per mouse | 10 | 6 | no/yes | 0.19 ± 0.03 | 0.05 ± 0.02 | MW | U=4 | | 0.003 | ** |
| Aged mice |  |  |  |  |  |  |  |  |  |  |  |
| CA1 pyramidal cells |  |  |  |  |  |  |  |  |  |  |  |
| Fig. 2B | Avg. firing rate in entire session | 620 | 669 | no/no | 1.3 ± 0.1 Hz | 1.0 ± 0.04 Hz | MW | U=184461 | | 0.0006 | *** |
|  | Distribution all cells - control vs CA3-APP |  |  |  |  |  | KS | D=0.1079 |  | 0.0011 | ** |
| | Pooled per mouse | 7 | 8 | yes/yes | 1.5 ± 0.2 Hz | 1.1 ± 0.1 Hz | two sample t test | t=2.275 | df=13 | 0.0405 | * |
| Fig. 2C | %Active cells CA1 | 7 | 8 | no/no | 81.4 ± 1.7 % | 77.5 ± 6.3 % | MW | U=28 | | >0.9999 | ns |
| CA1 interneurons |  |  |  |  |  |  |  |  |  |  |  |
| Fig. 2D | Avg. firing rate in entire session | 41 | 49 | no/no | 23.4 ± 1.3 Hz | 21.6 ± 1.7 Hz | MW | U=891 | | 0.3606 | ns |
|  | Distribution all cells – control vs CA3-APP |  |  |  |  |  | KS | D=0.335 |  | 0.0133 | * |
| | Pooled per mouse | 6 | 7 | yes/yes | 23.0 ± 1.2 Hz | 23.4 ± 4.0 Hz | two sample t test | t=0.08251 | df=11 | 0.9357 | ns |
| Fig. 2E | Speed modulation index | 41 | 49 | yes/yes | 0.15 ± 0.02 | 0.05 ± 0.01 | two sample t test | t=3.119 | df=89 | 0.0024 | ns |
|  | control different from 0? |  |  |  |  |  | Wilcoxon signed rank | W=621 |  | <0.0001 | *** |
|  | CA3-APP different from 0? |  |  |  |  |  | Wilcoxon signed rank | W=91 |  | 0.6667 | ns |
| | Pooled per mouse | 6 | 7 | yes/yes | 0.06 ± 0.07 | 0.01 ± 0.04 | two sample t test | t=0.6331 | df=11 | 0.5396 | ns |

| Supplementary Figure 3, similar analysis to Figure 2 but only includes data from run sub-sessions (no delay, 2-s delay, and 10-s delay) |  |  |  |  |  |  |  |  |  |  |  |
| --- | --- | --- | --- | --- | --- | --- | --- | --- | --- | --- | --- |
| Young adult mice |  |  |  |  |  |  |  |  |  |  |  |
| CA1 pyramidal cells |  |  |  |  |  |  |  |  |  |  |  |
| Suppl. Fig. 3C | Avg. firing rate in run sub-sessions | 624 | 640 | no/no | 1.6 ± 0.1 Hz | 1.2 ± 0.01Hz | MW | U=169295 |  | <0.0001 | *** |
|  | Distribution all cells - control vs CA3-APP |  |  |  |  |  | KS | D=0.08868 |  | 0.0134 | * |
|  | Pooled per mouse | 11 | 9 | yes/yes | 1.7 ± 0.1 Hz | 1.4 ± 0.1 Hz | two sample t test | t=1.390 | df=18 | 0.1815 | ns |
| Suppl. Fig. 3D | Burst index | 551 | 554 | no/no | 0.28 ± 0.01 | 0.23 ± 0.01 | MW | U=127323 |  | <0.0001 | *** |
|  | Pooled per mouse | 11 | 9 |  | 0.28 ± 0.03 | 0.23 ± 0.03 | two sample t test | t=1.383 | df=18 | 0.1837 | ns |
| CA1 interneurons |  |  |  |  |  |  |  |  |  |  |  |
| Suppl. Fig. 3E | Avg. firing rate in run sub-sessions | 90 | 60 | yes/yes | 34.2 ± 1.0 Hz | 33.3 ± 1.4 Hz | two sample t test | t=0.4890 | df=148 | 0.6256 | ns |
|  | Distribution all cells – control vs CA3-APP |  |  |  |  |  | KS | D=0.1333 |  | 0.5441 | ns |
|  | Pooled per mouse | 10 | 6 | yes/yes | 37.3 ± 3.5 Hz | 31.7 ± 3.1 Hz | two sample t test | t=1.092 | df=14 | 0.2931 | ns |
| Aged mice |  |  |  |  |  |  |  |  |  |  |  |
| CA1 pyramidal cells |  |  |  |  |  |  |  |  |  |  |  |
| Suppl. Fig. 3F | Avg. firing rate in run sub-sessions | 620 | 666 | no/no | 1.3 ± 0.1 Hz | 1.0 ± 0.04Hz | MW | U=183770 |  | 0.0007 | *** |
|  | Distribution all cells - control vs CA3-APP |  |  |  |  |  | KS | D=0.12 |  | 0.0002 | *** |
|  | Pooled per mouse | 7 | 8 | yes/yes | 1.6 ± 0.1 Hz | 1.3 ± 0.1 Hz | two sample t test | t=2.004 | df=13 | 0.0664 | ns |
| Suppl. Fig. 3G | Burst index | 499 | 527 | no/no | 0.24 ± 0.01 | 0.24 ± 0.01 | MW | U=125517 |  | 0.2083 | ns |
|  | Pooled per mouse | 7 | 8 | yes/yes | 0.24 ± 0.01 | 0.22 ± 0.03 | two sample t test | t=0.6062 | df=13 | 0.5548 | ns |
| Suppl. Fig. 3I | Age comparisons |  |  |  |  |  | Kruskal-Wallis | KW=30.72 |  | <0.0001 | *** |
|  | Young adult ctrl vs aged ctrl |  |  |  |  |  | Dunn's multiple comparisons test |  |  | 0.0383 | * |
|  | Aged CA3-APP vs aged CA3-APP |  |  |  |  |  | Dunn's multiple comparisons test |  |  | 0.2055 | ns |
| Suppl. Fig. 3H | Avg. firing rate in run sub-sessions | 41 | 49 | no/no | 29.7 ± 1.4 Hz | 25.9 ± 2.0 Hz | MW | U=857 |  | 0.1824 | ns |
|  | Distribution all cells – control vs CA3-APP |  |  |  |  |  | KS | D=0.12 |  | 0.0002 | *** |
|  | Pooled per mouse | 6 | 7 | yes/yes | 29.4 ± 0.9 Hz | 26.6 ± 4.1 Hz | two sample t test | t=0.6092 | df=11 | 0.5548 | ns |
| Suppl. Fig. 3J | Age comparisons |  |  |  |  |  | Kruskal-Wallis | KW=18.45 |  | 0.0004 | *** |
|  | Young adult ctrl vs aged ctrl |  |  |  |  |  | Dunn's multiple comparisons test |  |  | 0.0108 | * |
|  | Aged CA3-APP vs aged CA3-APP |  |  |  |  |  | Dunn's multiple comparisons test |  |  | 0.0116 | * |
|  | Distribution all cells – Young adult ctrl vs Aged ctrl |  |  |  |  |  | KS | D=0.3800 |  | 0.0006 | *** |
|  | Distribution all cells – Young adult CA3-APP vs Aged CA3-APP |  |  |  |  |  | KS | D=0.3433 |  | 0.0032 | ** |

**Supplementary Figure 4.**
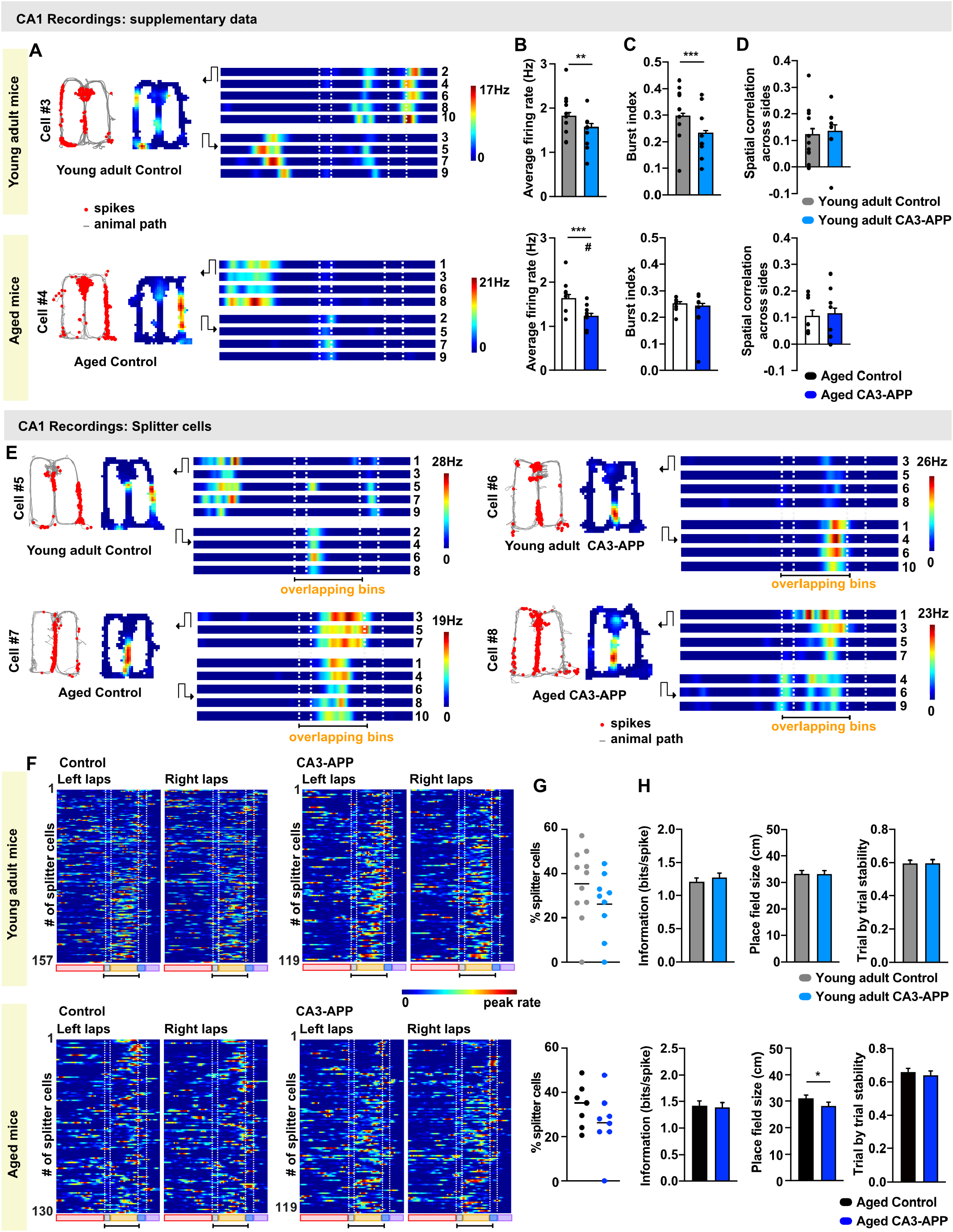
Characterization of CA1 place and splitter place cells. **A.** Examples of control CA1 place cells from young adult and aged mice. **B.** Average firing rate of place cells during behavior is lower in young adult and in aged CA3-APP mice compared to age-matched control mice (young adult: control 1.8 ± 0.1 Hz, n = 422 cells; CA3-APP 1.6 ± 0.1 Hz, n = 379 cells, p = 0.0014; aged: control 1.6 ± 0.1 Hz, n = 378 cells; CA3-APP 1.2 ± 0.1 Hz, n = 402, p = 0.001, MW test). In aged mice, the difference was also statistically different when data was analyzed per mouse (indicated by #, p = 0.019, n = 7 for control and n = 8 for CA3-APP mice). **C.** Burst index of place cells during behavior is lower in young adult CA3-APP mice compared to age-matched controls (control 0.30 ± 0.01, n = 422 cells; CA3-APP: 0.24 ± 0.01, n = 379 cells, p < 0.0001, MW test). In aged CA3-APP mice, there is no longer a difference to age-matched controls (control 0.25 ± 0.01, n = 378 cells; CA3-APP 0.24 ± 0.01, n = 402, p = 0.0640, MW test), but note that this is a consequence of reduced bursting in aged control mice (see **Supplementary Figure 3I**). **D.** Place cells of young adult and aged control mice were distinct between left and right return arms, as indicated by the low correlation coefficients for comparisons across arms. In CA3-APP mice of both age groups, the firing on either the left or the right arm remained as distinct as in age-matched controls (young adult: control 0.13 ± 0.02, n = 406 cells; CA3-APP 0.14 ± 0.02, n = 363 cells, p = 0.68; aged: control 0.11 ± 0.02, n = 354 cells; CA3-APP 0.12 ± 0.02, n = 374, p = 0.56, MW test). **E.** Example cells that were selectively active on the center arm in either only right-turn or left-turn laps (‘splitter cells’). One cell is from a young adult CA3-APP mouse, and one cell is from an aged CA3-APP mouse. Each panel includes the path (gray) with spike locations (red dots), a map of spike density (color-coded; 0 Hz, blue; maximum rate, red), linearized color-coded maps for each trial (0 Hz, blue; maximum rate, red; left turn trials, top; right turn trials; bottom). **F.** Color-coded firing rates of all splitter cells in each experimental group. Cells are ordered by their degree of selectivity for either correct right-turn or left-turn trials. **G.** The percentage of place cells that reached the criteria for splitter cells in the figure-8 maze were calculated per mouse. The percentage was not statistically different between genotypes in both the young adult and aged mice (young adult: control 35.4 ± 4.9%, n = 11 mice; CA3-APP 26.1 ± 4.8%, n = 9 mice; p = 0.20; aged: control 33.9 ± 3.9%, n = 7 mice; CA3-APP 26.4 ± 5.2%, n = 8 mice, p = 0.24, two sample T-test). **H.** The information score (young adult p = 0.41; aged p = 0.98, MW test), and trial-by-trial stability of splitter cells (young adult p = 0.85; aged p = 0.93, MW test) did not differ between CA3-APP mice and age-matched control mice. Similarly to what was observed in place cells in general, there was a significant reduction of place field size of splitter cells from aged, but not from young adult CA3-APP mice compared to aged-matched control mice (young adult: control 32.7 ± 1.2 cm, n = 173 cells; CA3-APP 32.8 ± 1.3 cm, n = 119, p = 0.56; aged: control 31.1 ± 1.2 cm, n = 130; CA3-APP: 28.2 ± 1.3 cm, n = 119, p = 0.037, MW test). All bar graphs show mean ± SEM. See **Supplementary Table 6** for statistics.

**Supplementary Table 6.** Properties of CA1 Place cells. Data and statistics for Figure 3 and Supplementary Figure 4.

| Fig. panel | Graph description | n (ctrl) | n (tg) | Normal Dist? | ctrl mean ± SEM | tg mean ± SEM | Statistical test | Stat. metrics | Df | P value | Sig. |
| --- | --- | --- | --- | --- | --- | --- | --- | --- | --- | --- | --- |
| **Young adult mice** |  |  |  |  |  |  |  |  |  |  |  |
| **Fig. 3C** | **% place cells (PC)** | 422 | 379 |  | 67.1% | 58.8% | n too high for Chi-square test |  |  |  |  |
| | per mouse | 11 | 9 | yes/no | 69.3 ± 5.6% | 62.8 ± 5.3% | MW | U=34.5 | | 0.2689 | ns |
| **Suppl. Fig. 4A** | **Avg. PC firing rate** | 422 | 379 | no/no | 1.8 ± 0.1Hz | 1.6 ± 0.1Hz | MW | U=69537 | | 0.0014 | ** |
| | per mouse | 11 | 9 | yes/yes | 1.9 ± 0.1Hz | 1.5 ± 0.1Hz | two sample t test | t=2.087 | df=18 | 0.0513 | ns |
| **Fig. 3D** | **PC peak firing rate** | 422 | 379 | No/no | 20.3 ± 0.7Hz | 17.7 ± 0.7Hz | MW | U=71067 | | 0.0065 | ** |
| | per mouse | | | yes/yes | 19.7 ± 1.5Hz | 18.6 ± 0.9Hz | two sample t test | t=0.6160 | df=18 | 0.5456 | ns |
| **Suppl. Fig. 4B** | **PC Burst index** | 422 | 379 | no/no | 0.30 ± 0.01 | 0.24 ± 0.01 | MW | U=61515 | | <0.0001 | *** |
| | per mouse | | | yes/yes | 0.31 ± 0.03 | 0.23 ± 0.03 | two sample t test | t=2.049 | df=18 | 0.0554 | ns |
| **Fig. 3E** | **PC Information (b/sp)** | 422 | 379 | no/no | 1.3 ± 0.04 | 1.4 ± 0.4 | MW | U=74987 | | 0.1276 | ns |
| | per mouse | | | yes/yes | 1.1 ± 0.1 | 1.4 ± 0.1 | two sample t test | t=1.835 | df=18 | 0.083 | ns |
| **Fig. 3F** | **# of place fields** | 422 | 379 | no/no | 2.5 ± 0.1 | 2.3 ± 0.1 | MW | U=73057 | | 0.0281 | * |
| | per mouse | | | yes/no | 2.6 ± 0.2 | 2.3 ± 0.1 | MW | U=27.5 | | 0.0992 | ns |
| **Fig. 3G** | **Size of place fields** | 422 | 379 | no/no | 28.6 ± 0.7cm | 28.0 ± 0.7cm | MW | U=79217 | | 0.8182 | ns |
| | per mouse | | | yes/no | 28.5 ± 1.2cm | 30.2 ± 2.0cm | MW | U=45 | | 0.7664 | ns |
| **Fig. 3H** | **Trial by trial stability** | 422 | 379 | no/no | 0.58 ± 0.01 | 0.59 ± 0.01 | MW | U=78112 | | 0.5701 | ns |
| | per mouse | | | yes/yes | 0.53 ± 0.03 | 0.59 ± 0.01 | two sample t test | t=1.626 | df=18 | 0.1213 | ns |
| **Suppl. Fig. 4C** | **Across sides corr.<sup>5</sup>** | 406 | 363 | no/no | 0.13 ± 0.02 | 0.14 ± 0.02 | MW | U=73431 | | 0.6819 | ns |
| | per mouse | | | yes/no | 0.12 ± 0.03 | 0.14 ± 0.03 | MW | U=41 | | 0.5516 | ns |
| **Suppl. Fig. 4G** | **% Splitter cells (of PC)** | 173 | 119 |  | 41.0% | 31.4% |  |  |  |  |  |
| | per mouse | 11 | 9 | yes/yes | 35.4 ± 4.9% | 26.1 ± 4.8% | two sample t test | t=1.336 | df=18 | 0.1983 | ns |
| **Suppl. Fig. 4H** | **Splitter cells: Information** | 173 | 119 | no/no | 1.2 ± 0.6 | 1.3 ± 0.7 | MW | U=9708 | | 0.4089 | ns |
| | **Splitter cells: Place field size** | 173 | 119 | no/no | 32.7 ± 1.2 | 32.8 ± 1.3 | MW | U=9893 | | 0.5626 | ns |
| | **Splitter cells: Trial by trial corr.** | 173 | 119 | no/no | 0.60 ± 0.02 | 0.60 ± 0.02 | MW | U=10159 | | 0.8501 | ns |
| **Aged mice** |  |  |  |  |  |  |  |  |  |  |  |
| **Fig. 3C** | **% place cells (PC)** | 378 | 402 |  | 61.0% | 60.1% | n too high for Chi-square test |  |  |  |  |
| | per mouse | 7 | 8 | yes/no | 61.4 ± 2.4% | 59.1 ± 5.2% | MW | U=26 | | 0.8665 | ns |
| **Suppl. Fig. 4A** | **Avg. PC firing rate** | 378 | 402 | no/no | 1.6 ± 0.1 Hz | 1.2 ± 0.1 Hz | MW | U=65597 | | 0.001 | *** |
| | per mouse | 7 | 8 | yes/yes | 1.7 ± 0.2 Hz | 1.2 ± 0.1 Hz | two sample t test | t=2.690 | df=13 | 0.0186 | * |
| **Fig. 3D** | **PC peak firing rate** | 378 | 402 | no/no | 21.3 ± 0.8 Hz | 14.8 ± 0.6 Hz | MW | U=54391 | | <0.0001 | *** |
| | per mouse | | | yes/yes | 21.0 ± 1.3 Hz | 14.8 ± 1.1 Hz | two sample t test | t=3.612 | df=13 | 0.0032 | ** |
| **Suppl. Fig. 4B** | **PC Burst index** | 378 | 402 | no/no | 0.25 ± 0.01 | 0.24 ± 0.01 | MW | U=70153 | | 0.064 | ns |
| | per mouse | | | yes/no | 0.25 ± 0.01 | 0.22 ± 0.03 | MW | U=27 | | 0.9551 | ns |
| **Fig. 3E** | **PC Information (b/sp)** | 378 | 402 | no/no | 1.5 ± 0.04 | 1.4 ± 0.04 | MW | U=71402 | | 0.1457 | ns |
| | per mouse | | | yes/yes | 1.5 ± 0.1 | 1.4 ± 0.1 | two sample t test | t=0.4900 | df=13 | 0.6323 | ns |
| **Fig. 3F** | **# of place fields** | 378 | 402 | no/no | 2.4 ± 0.1 | 2.0 ± 0.1 | MW | U=68110 | | 0.0083 | ** |
| | per mouse | | | yes/yes | 2.5 ± 0.2 | 1.9 ± 0.1 | two sample t test | t=3.300 | df=13 | 0.0057 | ** |
| **Fig. 3G** | **Size of place fields** | 378 | 402 | no/no | 27.9 ± 0.7cm | 25.0 ± 0.6cm | MW | U=63196 | | <0.0001 | *** |
| | per mouse | | | yes/yes | 27.2 ± 1.2cm | 25.9 ± 1.5cm | two sample t test | t=0.6297 | df=13 | 0.5398 | ns |
| **Fig. 3H** | **Trial by trial stability** | 378 | 402 | no/no | 0.65 ± 0.01 | 0.6 ± 0.01 | MW | U=67973 | | 0.0149 | * |
| | per mouse | | | yes/no | 0.63 ± 0.03 | 0.6 ± 0.02 | MW | U=17 | | 0.2319 | ns |
| **Suppl. Fig. 4C** | **Across sides corr.<sup>5</sup>** | 354 | 374 | no/no | 0.11 ± 0.02 | 0.12 ± 0.02 | MW | U=64532 | | 0.5570 | ns |
| | per mouse | | | no/yes | 0.11 ± 0.03 | 0.11 ± 0.03 | MW | U=25 | | 0.7789 | ns |
| **Suppl. Fig. 4G** | **% Splitter cells (of PC)** | 130 | 119 |  | 34.4% | 29.6% |  |  |  |  |  |
| | per mouse | 7 | 8 | yes/yes | 33.9 ± 3.9% | 26.4 ± 5.2% | two sample t test | t=1.220 | df=13 | 0.2440 | ns |
| **Suppl. Fig. 4H** | **Splitter cells: Information** | 130 | 119 | no/no | 1.4 ± 0.1 | 1.4 ± 0.1 | MW | U=7722 | | 0.9824 | ns |
| | **Splitter cells: Place field size** | 130 | 119 | no/no | 31.1 ± 1.2 | 28.3 ± 1.3 | MW | U=6551 | | 0.0371 | * |
| | **Splitter cells: Trial by trial corr.** | 130 | 119 | no/no | 0.66 ± 0.02 | 0.65 ± 0.02 | MW | U=7688 | | 0.9347 | ns |

<sup>5</sup> less cells as some are in central arm

**Supplementary Figure 5.**
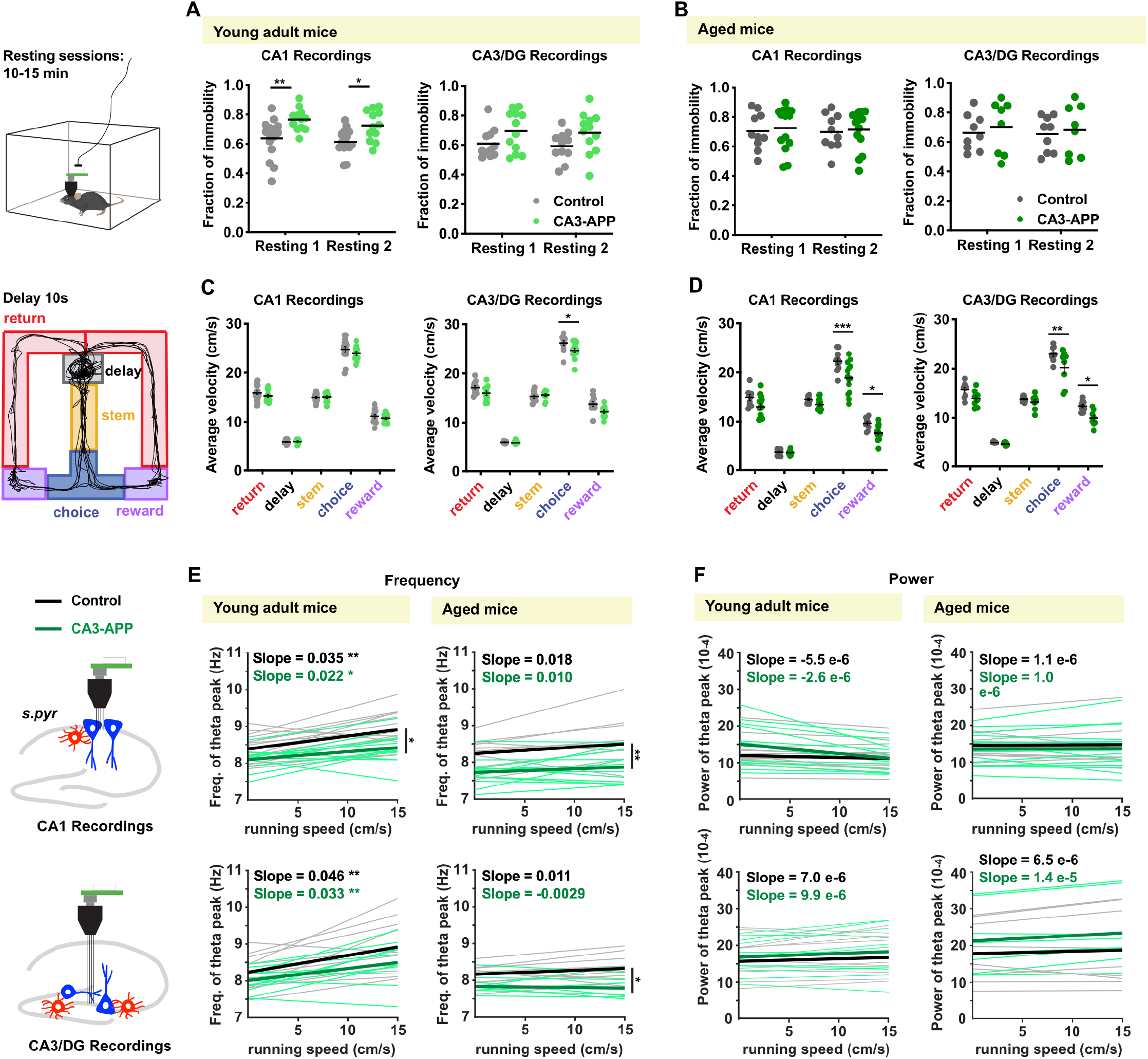
Controls for velocity differences across different parts of the behavior task. **A.** Fraction of immobility (<2 cm/s) during rest sessions. In the recording sessions where electrodes were placed at CA1 sites, CA3-APP young adult mice spent, on average, less time moving than age-matched controls. A similar tendency, although not statistically significant, was observed in the recordings from CA3/DG sites. Mice were resting for an average of ~60% of their time during the 10-15 minute sessions in the home cage. **B.** No difference in the fraction of immobility was detected in aged CA3-APP mice compared to aged control mice. See **Supplementary Table 7** for data and statistics. **C.** Analysis of running speed in different sections of the figure-8 maze. Running speed of young adult CA3-APP mice did not differ from control littermates in any sections of the maze during recording sessions from CA1 sites. A small reduction in running speed in only the choice section was detected for young adult CA3-APP mice compared to young adult controls during recording sessions from CA3/DG sites. **D.** Same analyses as in C for aged mice. Running speed of CA3-APP mice was lower in the choice and reward sections of the behavior task. See **Supplementary Table 8** for data and statistics. **E.** Correlation between theta frequency and running speed for periods when movement was >2 cm/s. For each mouse we pooled data from all behavior sessions and obtained a regression line of running speed vs. theta frequency (shaded grey lines, control mice; shaded green lines, CA3-APP mice). The average regression lines per genotype are presented as thick lines (black, control; green, CA3-APP). In young adult mice theta frequency was modulated by locomotion speed (slopes compared to zero, all p values < 0.05, Wilcoxon signed rank tests), but theta frequency did not change with speed in aged mice (all p values > 0.05, Wilcoxon signed rank tests; see **Supplementary Table 8** for statistics). In CA3-APP mice, there was no change in the speed-frequency slope compared to controls (all p values > 0.05, Wilcoxon signed rank tests), but the offset (y-intersect) shifted to lower values (CA1 recordings: young adult p = 0.043; aged p = 0.0043; CA3/DG recordings: young adult p = 0.18; aged p = 0.049, two sample t-test). **F.** Same analysis as in E, but for the correlation between running speed and theta power, rather than theta frequency. We did not find a relation between running speed and theta power (slopes compared to zero, all p values > 0.05, Wilcoxon signed rank test), and the intersects of the slopes did not differ between CA3-APP mice and age-matched controls. See **Supplementary Table 8** for statistical analyses of slopes and intersects.

**Supplementary Table 7.**
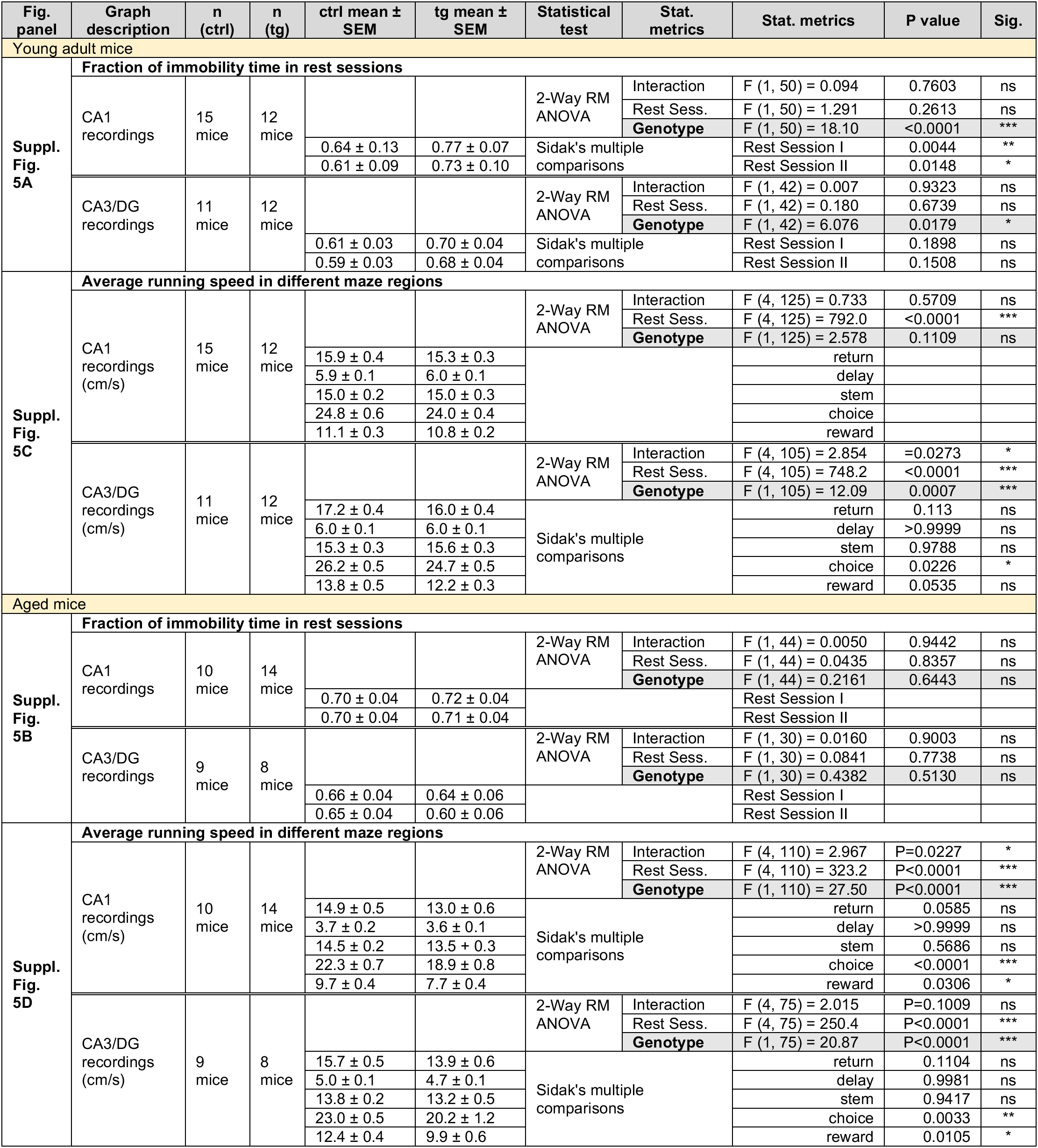
Control data for immobility time and running speed in different sections of the maze, related to Supplementary Figure 5A-D.

**Supplementary Table 8.**
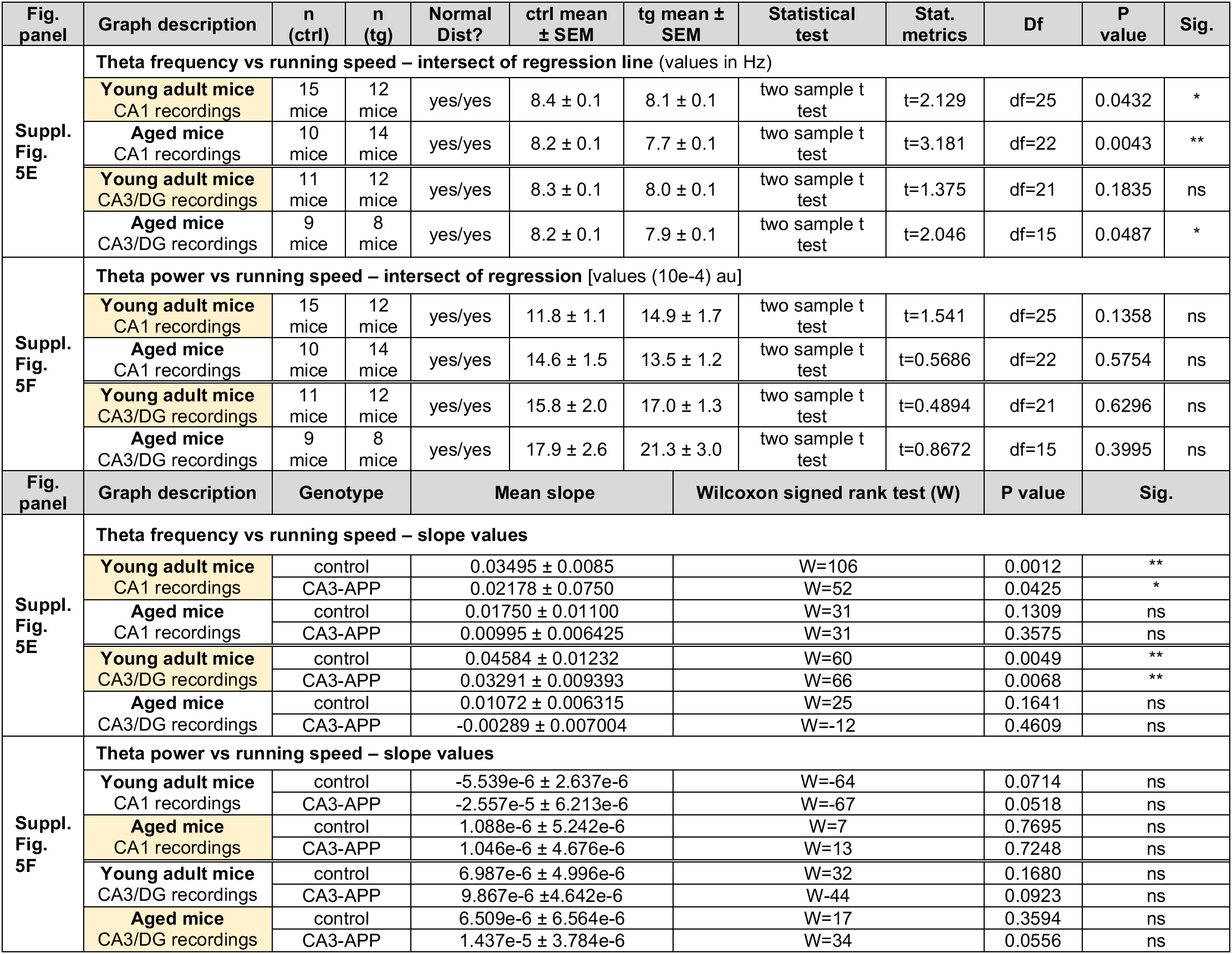
Correlation between theta oscillations and mouse velocity during the behavioral task, related to Supplementary Figure 5E and F.

**Supplementary Figure 6.**
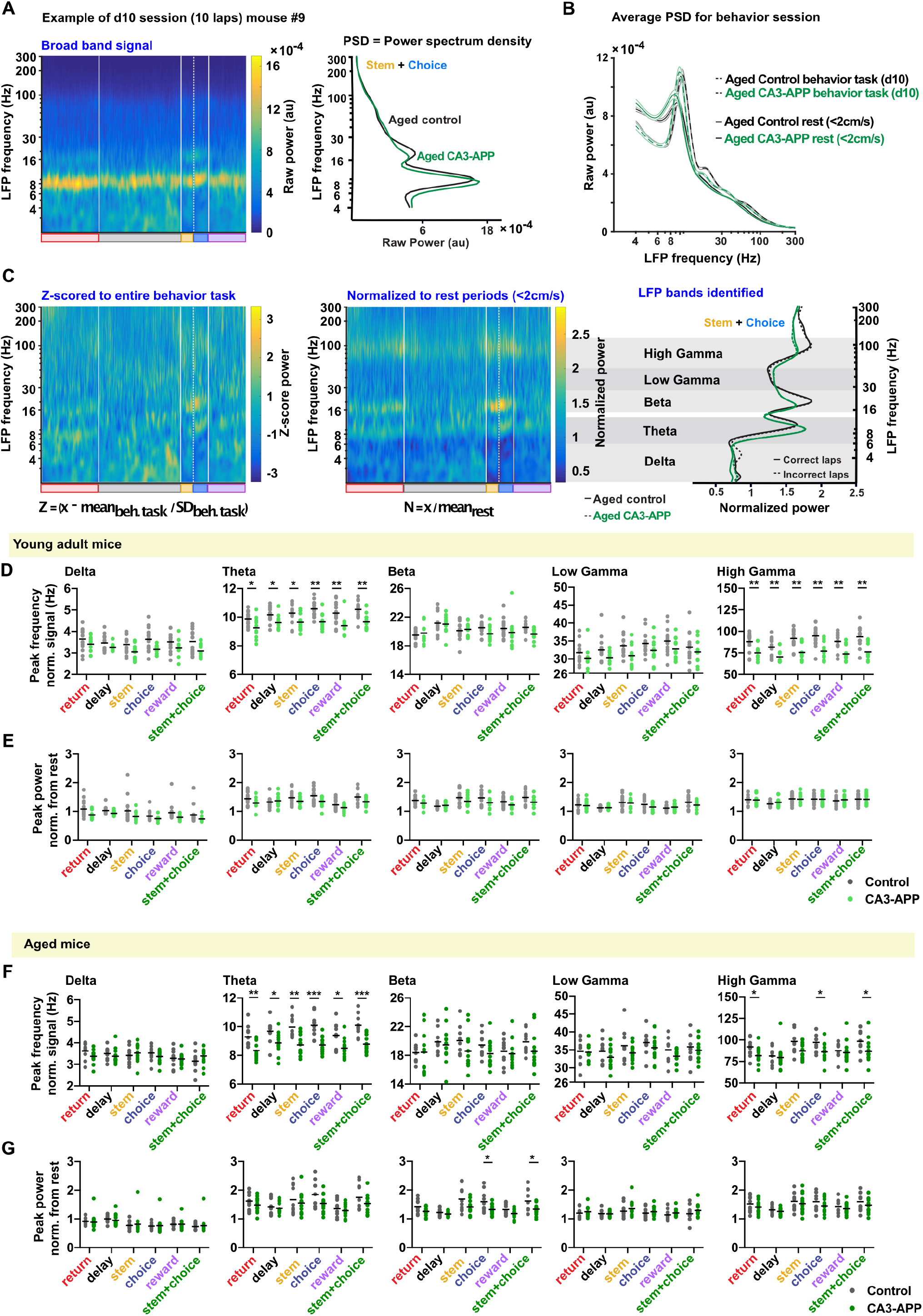
APP expression in CA3 reduced CA1 theta frequency for all maze sections and CA1 high-gamma frequency for most maze sections. **A.** Example spectrogram of broadband CA1 LFP signal from a figure-8 behavior session (10 laps in the 10-s delay condition from an aged control mouse). Power spectrum density (PSD) plot obtained for the same session (signal from stem and choice segments) where oscillations at higher frequencies are not well delineated. **B.** Average PSD from aged mice during the spatial alternation task (10-s delay laps – continuous line) and during a rest session in the home cage (rest - dashed line) before the behavioral task. During the rest sessions, only periods with no/low movement (<2 cm/s) were considered. **C.** Comparison between normalization by Z-scoring and normalization to rest sessions. The latter method more effectively corrected for the higher power at lower frequencies and was therefore selected for further analyses. Based on the normalized power spectra, five LFP bands were defined: delta (2-6 Hz), theta (6-12 Hz), beta (14-26 Hz), low gamma (26-50 Hz) and high gamma (50-120 Hz). We found no differences in LFP power and frequency between correct (continuous line) and incorrect laps (dashed lines). **D.** Quantification of peak frequency of the normalized LFP signal, by band and by sections of the maze. Theta frequency and high gamma frequency were reduced in all sections of the maze in young adult CA3-APP mice compared to age-matched controls (p < 0.05, MW test adjusted for multiple comparisons by Holm-Šídák method). No differences were observed for delta, beta and low gamma frequencies at this age. **E.** Quantification of peak power of the normalized LFP signal, by band and by sections of the maze. No differences in power were detected in any of the LFP bands (p > 0.5, MW test adjusted for multiple comparisons by Holm-Šídák method). **F.** A reduction in theta frequency was found in all sections of the maze for aged CA3-APP mice compared to age-matched controls. In the high gamma band, a frequency reduction was observed in the return and choice segments of the maze for aged CA3-APP mice compared to age-matched controls (p < 0.05, MW test adjusted for multiple comparisons by Holm-Šídák method). Data are displayed as in D. **G**. There was a reduction in the power of beta oscillations in CA3-APP aged mice, particularly in the choice segment of the maze (choice: control 1.6 ± 0.1, CA3-APP 1.3 ± 0.05, p = 0.042, MW test adjusted for multiple comparisons by Holm-Šídák method). Data are displayed as in E. See **Supplementary Table 9** for summary data and statistical analysis.

**Supplementary Table 9.** Analysis of normalized CA1 LFP data, related to Figure 4 and Supplementary Figure 6.

| Fig. panel | Graph description | n= |  | Frequency (Hz) |  | MW test<sup>#</sup> |  |  | Amplitude (normalized) |  | Mann-Whitney test<sup>#</sup> |  |  |
| --- | --- | --- | --- | --- | --- | --- | --- | --- | --- | --- | --- | --- | --- |
|  |  | ctrl | tg | ctrl | tg | U= | P value | Sig. | ctrl | tg | U= | P value | Sig. |
| Young adult mice |  |  |  |  |  |  |  |  |  |  |  |  |  |
| Suppl. Fig. 6D, E | CA1 delta normalized |  |  |  |  |  |  |  |  |  |  |  |  |
|  | Return | 15 | 12 | 3.6 ± 0.1 | 3.4 ± 0.1 | 63 | 0.359246 | ns | 1.1 ± 0.1 | 0.9 ± 0.04 | 44 | 0.09478 | ns |
|  | Delay | 15 | 12 | 3.5 ± 0.1 | 3.3 ± 0.05 | 66 | 0.359246 | ns | 1.0 ± 0.03 | 0.9 ± 0.02 | 39 | 0.059709 | ns |
|  | Stem | 15 | 12 | 3.4 ± 0.1 | 3.1 ± 0.1 | 44 | 0.138746 | ns | 1.0 ± 0.1 | 0.8 ± 0.05 | 59 | 0.138536 | ns |
|  | Choice | 15 | 12 | 3.6 ± 0.1 | 3.2 ± 0.1 | 44 | 0.138746 | ns | 0.8 ± 0.04 | 0.7 ± 0.02 | 49 | 0.09478 | ns |
|  | Reward | 15 | 12 | 3.5 ± 0.1 | 3.2 ± 0.1 | 50 | 0.15034 | ns | 1.0 ± 0.1 | 0.8 ± 0.02 | 38 | 0.059709 | ns |
| 4B, C | stem+choice | 15 | 12 | 3.5 ± 0.1 | 3.1 ± 0.1 | 47 | 0.138746 | ns | 0.9 ± 0.1 | 0.7 ± 0.03 | 44 | 0.09478 | ns |
| Suppl. Fig. 6D, E | CA1 theta normalized |  |  |  |  |  |  |  |  |  |  |  |  |
|  | return | 15 | 12 | 9.9 ± 0.1 | 9.3 ± 0.2 | 18 | 0.001066 | * | 1.4 ± 0.1 | 1.3 ± 0.1 | 59 | 0.449258 | ns |
|  | delay | 15 | 12 | 10.2 ± 0.1 | 9.6 ± 0.1 | 23 | 0.002999 | * | 1.3 ± 0.04 | 1.4 ± 0.1 | 80 | 0.648303 | ns |
|  | stem | 15 | 12 | 10.3 ± 0.2 | 9.6 ± 0.2 | 33 | 0.017649 | * | 1.5 ± 0.1 | 1.4 ± 0.05 | 62 | 0.454183 | ns |
|  | choice | 15 | 12 | 10.6 ± 0.1 | 9.7 ± 0.2 | 37 | 0.025902 | ** | 1.5 ± 0.1 | 1.34 ± 0.1 | 45 | 0.157405 | ns |
|  | reward | 15 | 12 | 10.3 ± 0.2 | 9.4 ± 0.2 | 42 | 0.036834 | * | 1.2 ± 0.05 | 1.1 ± 0.04 | 68 | 0.50936 | ns |
| 4B, C | stem+choice | 15 | 12 | 10.5 ± 0.1 | 9.7 ± 0.2 | 45 | 0.036834 | ** | 1.5 ± 0.1 | 1.3 ± 0.05 | 53 | 0.321309 | ns |
| Suppl. Fig. 6D, E | CA1 beta normalized |  |  |  |  |  |  |  |  |  |  |  |  |
|  | Return | 15 | 12 | 19.5 ± 0.2 | 19.8 ± 0.4 | 88 | 0.996711 | ns | 1.4 ± 0.03 | 1.3 ± 0.04 | 58 | 0.415764 | ns |
|  | Delay | 15 | 12 | 21.2 ± 0.4 | 21.0 ± 0.4 | 88 | 0.996711 | ns | 1.2 ± 0.03 | 1.2 ± 0.03 | 70 | 0.573964 | ns |
|  | Stem | 15 | 12 | 20.1 ± 0.3 | 20.3 ± 0.3 | 85 | 0.995008 | ns | 1.5 ± 0.1 | 1.3 ± 0.1 | 72 | 0.573964 | ns |
|  | Choice | 15 | 12 | 20.5 ± 0.2 | 19.7 ± 0.3 | 48 | 0.190717 | ns | 1.5 ± 0.1 | 1.3 ± 0.1 | 54 | 0.406274 | ns |
|  | Reward | 15 | 12 | 20.4 ± 0.3 | 19.9 ± 0.6 | 60 | 0.474647 | ns | 1.3 ± 0.05 | 1.2 ± 0.04 | 62 | 0.454183 | ns |
| 4B, C | stem+choice | 14 | 12 | 20.6 ± 0.2 | 19.6 ± 0.2 | 36 | 0.073651 | ns | 1.5 ± 0.1 | 1.3 ± 0.1 | 51 | 0.406274 | ns |
| Suppl. Fig. 6D, E | CA1 low gamma normalized |  |  |  |  |  |  |  |  |  |  |  |  |
|  | Return | 14 | 11 | 31.7 ± 0.8 | 30.2 ± 0.9 | 52 | 0.505029 | ns | 1.2 ± 0.05 | 1.2 ± 0.1 | 74 | 0.99522 | ns |
|  | Delay | 13 | 10 | 32.5 ± 1.0 | 30.3 ± 0.6 | 42 | 0.505029 | ns | 1.1 ± 0.02 | 1.1 ± 0.03 | 61 | 0.99522 | ns |
|  | Stem | 15 | 11 | 33.7 ± 0.8 | 30.9 ± 0.9 | 42 | 0.198602 | ns | 1.3 ± 0.1 | 1.3 ± 0.1 | 79 | 0.99522 | ns |
|  | choice | 13 | 11 | 34.3 ± 0.9 | 32.4 ± 1.0 | 48 | 0.505029 | ns | 1.2 ± 0.1 | 1.1 ± 0.05 | 49 | 0.750571 | ns |
|  | reward | 15 | 12 | 35.0 ± 0.9 | 32.8 ± 0.9 | 55 | 0.384866 | ns | 1.1 ± 0.03 | 1.1 ± 0.04 | 76 | 0.945289 | ns |
| 4B, C | stem+choice | 15 | 11 | 33.2 ± 1.1 | 32.0 ± .1 | 60 | 0.584845 | ns | 1.3 ± 0.1 | 1.2 ± 0.1 | 53 | 0.875465 | ns |
| Suppl. Fig. 6D, E | CA1 high gamma normalized |  |  |  |  |  |  |  |  |  |  |  |  |
|  | return | 15 | 12 | 88.1 ± 2.7 | 75.0 ± 2.5 | 21 | 0.002258 | ** | 1.4 ± 0.04 | 1.4 ± 0.1 | 79 | 0.994561 | ns |
|  | delay | 15 | 12 | 81.6 ± 2.7 | 70.5 ± 2.3 | 24 | 0.003752 | ** | 1.3 ± 0.02 | 1.3 ± 0.04 | 78 | 0.994561 | ns |
|  | stem | 15 | 12 | 91.8 ± 2.6 | 75.5 ± 2.6 | 25 | 0.003752 | ** | 1.4 ± 0.04 | 1.4 ± 0.1 | 81 | 0.994561 | ns |
|  | choice | 15 | 12 | 94.7 ± 2.9 | 77.1 ± 2.9 | 28 | 0.005209 | ** | 1.4 ± 0.04 | 1.4 ± 0.1 | 85 | 0.994561 | ns |
|  | reward | 15 | 12 | 88.4 ± 2.5 | 73.7 ± 2.6 | 31 | 0.006179 | ** | 1.4 ± 0.04 | 1.4 ± 0.1 | 82 | 0.994561 | ns |
| 4B, C | stem+choice | 15 | 12 | 93.7 ± 2.9 | 76.2 ± 2.8 | 32 | 0.006179 | ** | 1.4 ± 0.04 | 1.4 ± 0.1 | 84 | 0.994561 | ns |
| Aged Mice |  |  |  |  |  |  |  |  |  |  |  |  |  |
| Suppl. Fig. 6F, G | CA1 delta normalized |  |  |  |  |  |  |  |  |  |  |  |  |
|  | return | 10 | 14 | 3.6 ± 0.1 | 3.4 ± 0.1 | 36 | 0.095574 | ns | 0.9 ± 0.03 | 0.9 ± 0.1 | 40 | 0.159446 | ns |
|  | delay | 10 | 14 | 3.5 ± 0.1 | 3.4 ± 0.1 | 46 | 0.172086 | ns | 1.0 ± 0.04 | 1.0 ± 0.05 | 46 | 0.172086 | ns |
|  | stem | 10 | 14 | 3.4 ± 0.1 | 3.5 ± 0.1 | 57 | 0.471586 | ns | 0.8 ± 0.05 | 0.8 ± 0.1 | 69 | 0.977098 | ns |
|  | choice | 10 | 14 | 3.5 ± 0.1 | 3.4 ± 0.1 | 49 | 0.234983 | ns | 0.8 ± 0.04 | 0.8 ± 0.1 | 56 | 0.436617 | ns |
|  | reward | 10 | 14 | 3.3 ± 0.1 | 3.3 ± 0.1 | 66 | 0.840751 | ns | 0.8 ± 0.04 | 0.8 ± 0.5 | 59 | 0.54577 | ns |
| 4E,F | stem+choice | 10 | 14 | 3.2 ± 0.1 | 3.4 ± 0.1 | 48 | 0.212499 | ns | 0.7 ± 0.04 | 0.8 ± 0.07 | 64 | 0.752095 | ns |
| Suppl. Fig. 6F, G | CA1 theta normalized |  |  |  |  |  |  |  |  |  |  |  |  |
|  | Return | 10 | 14 | 9.3 ± 0.2 | 8.3 ± 0.2 | 23 | 0.00478 | ** | 1.6 ± 0.1 | 1.5 ± 0.1 | 53 | 0.340738 | ns |
|  | Delay | 10 | 14 | 9.7 ± 0.2 | 8.9 ± 0.2 | 27 | 0.010735 | * | 1.4 ± 0.1 | 1.4 ± 0.05 | 64 | 0.752095 | ns |
|  | Stem | 10 | 14 | 10.0 ± 0.2 | 8.7 ± 0.2 | 20 | 0.002432 | ** | 1.7 ± 0.2 | 1.5 ± 0.1 | 65 | 0.796094 | ns |
|  | Choice | 10 | 14 | 10.1 ± 0.3 | 8.7 ± 0.1 | 11 | 0.000198 | *** | 1.8 ± 0.1 | 1.5 ± 0.1 | 48 | 0.212499 | ns |
|  | Reward | 10 | 14 | 9.4 ± 0.3 | 8.5 ± 0.2 | 30 | 0.018545 | * | 1.4 ± 0.1 | 1.3 ± 0.1 | 62 | 0.666477 | ns |
| 4E, F | stem+choice | 10 | 14 | 10.1 ± 0.2 | 8.8 ± 0.2 | 10 | 0.000142 | *** | 1.7 ± 0.2 | 1.5 ± 0.1 | 56 | 0.436617 | ns |
| Suppl. Fig. 6F, G | CA1 beta normalized |  |  |  |  |  |  |  |  |  |  |  |  |
|  | Return | 10 | 14 | 18.4 ± 0.4 | 18.4 ± 0.8 | 59 | 0.54577 | ns | 1.4 ± 0.1 | 1.3 ± 0.04 | 41 | 0.095643 | ns |
|  | Delay | 10 | 14 | 19.9 ± 0.5 | 19.5 ± 0.7 | 61 | 0.625096 | ns | 1.2 ± 0.04 | 1.2 ± 0.02 | 54 | 0.371155 | ns |
|  | Stem | 10 | 14 | 20.1 ± 0.6 | 18.6 ± 0.7 | 37 | 0.055911 | ns | 1.7 ± 0.1 | 1.4 ± 0.1 | 42 | 0.108344 | ns |
|  | Choice | 10 | 14 | 19.4 ± 0.5 | 18.2 ± 0.5 | 38 | 0.101012 | ns | 1.6 ± 0.1 | 1.3 ± 0.05 | 32 | 0.042158 | * |
|  | Reward | 10 | 14 | 18.6 ± 0.6 | 18.2 ± 0.5 | 61 | 0.625096 | ns | 1.3 ± 0.1 | 1.2 ± 0.04 | 44 | 0.137503 | ns |
| 4E, F | stem+choice | 10 | 14 | 19.9 ± 0.5 | 18.6 ± 0.6 | 37 | 0.055911 | ns | 1.6 ± 0.1 | 1.3 ± 0.1 | 42 | 0.042681 | * |
| Suppl. Fig. 6F, G | CA1 low gamma normalized |  |  |  |  |  |  |  |  |  |  |  |  |
|  | Return | 10 | 13 | 34.6 ± 1.2 | 34.4 ± 0.6 | 63 | 0.927375 | ns | 1.2 ± 0.1 | 1.2 ± 0.05 | 56 | 0.604913 | ns |
|  | Delay | 10 | 14 | 34.6 ± 1.1 | 33.0 ± 0.7 | 50 | 0.240871 | ns | 1.2 ± 0.04 | 1.2 ± 0.02 | 66 | 0.840751 | ns |
|  | Stem | 10 | 14 | 36.2 ± 1.4 | 34.1 ± 0.6 | 51 | 0.265308 | ns | 1.3 ± 0.1 | 1.4 ± 0.1 | 54 | 0.371155 | ns |
|  | choice | 10 | 14 | 37.0 ± 1.0 | 35.6 ± 0.9 | 46 | 0.172086 | ns | 1.1 ± 0.1 | 1.2 ± 0.04 | 61 | 0.625096 | ns |
|  | reward | 10 | 14 | 35.0 ± 1.1 | 33.3 ± 0.5 | 44 | 0.137503 | ns | 1.1 ± 0.1 | 1.2 ± 0.04 | 43 | 0.122267 | ns |
| 4E, F | stem+choice | 10 | 14 | 35.9 ± 1.3 | 34.9 ± 0.8 | 57 | 0.471586 | ns | 1.2 ± 0.1 | 1.3 ± 0.1 | 51 | 0.284681 | ns |
| Suppl. Fig. 6F, G | CA1 high gamma normalized |  |  |  |  |  |  |  |  |  |  |  |  |
|  | return | 10 | 14 | 91.9 ± 3.5 | 81.8 ± 2.4 | 32 | 0.026016 | * | 1.5 ± 0.1 | 1.4 ± 0.1 | 54 | 0.371155 | ns |
|  | delay | 10 | 14 | 81.4 ± 3.0 | 79.6 ± 3.4 | 54 | 0.371155 | ns | 1.3 ± 0.5 | 1.3 ± 0.03 | 56 | 0.411455 | ns |
|  | stem | 10 | 14 | 98.3 ± 4.5 | 87.2 ± 2.3 | 38 | 0.064329 | ns | 1.6 ± 0.1 | 1.5 ± 0.1 | 60 | 0.584845 | ns |
|  | choice | 10 | 14 | 97.1 ± 3.5 | 86.2 ± 2.6 | 36 | 0.048404 | * | 1.5 ± 0.1 | 1.5 ± 0.1 | 50 | 0.25905 | ns |
|  | reward | 10 | 14 | 87.6 ± 2.8 | 85.3 ± 3.4 | 48 | 0.212499 | ns | 1.4 ± 0.1 | 1.3 ± 0.05 | 55 | 0.403111 | ns |
| 4E, F | stem+choice | 10 | 14 | 98.4 ± 4.3 | 86.7 ± 3.5 | 35 | 0.041717 | * | 1.6 ± 0.1 | 1.5 ± 0.1 | 55 | 0.403111 | ns |
| * mean + SEM # adjusted for multiple comparisons by Holm-Sidak method |  |  |  |  |  |  |  |  |  |  |  |  |  |

\* mean ± SEM # adjusted for multiple comparisons by Holm-Šidák method

**Supplementary Figure 7.**
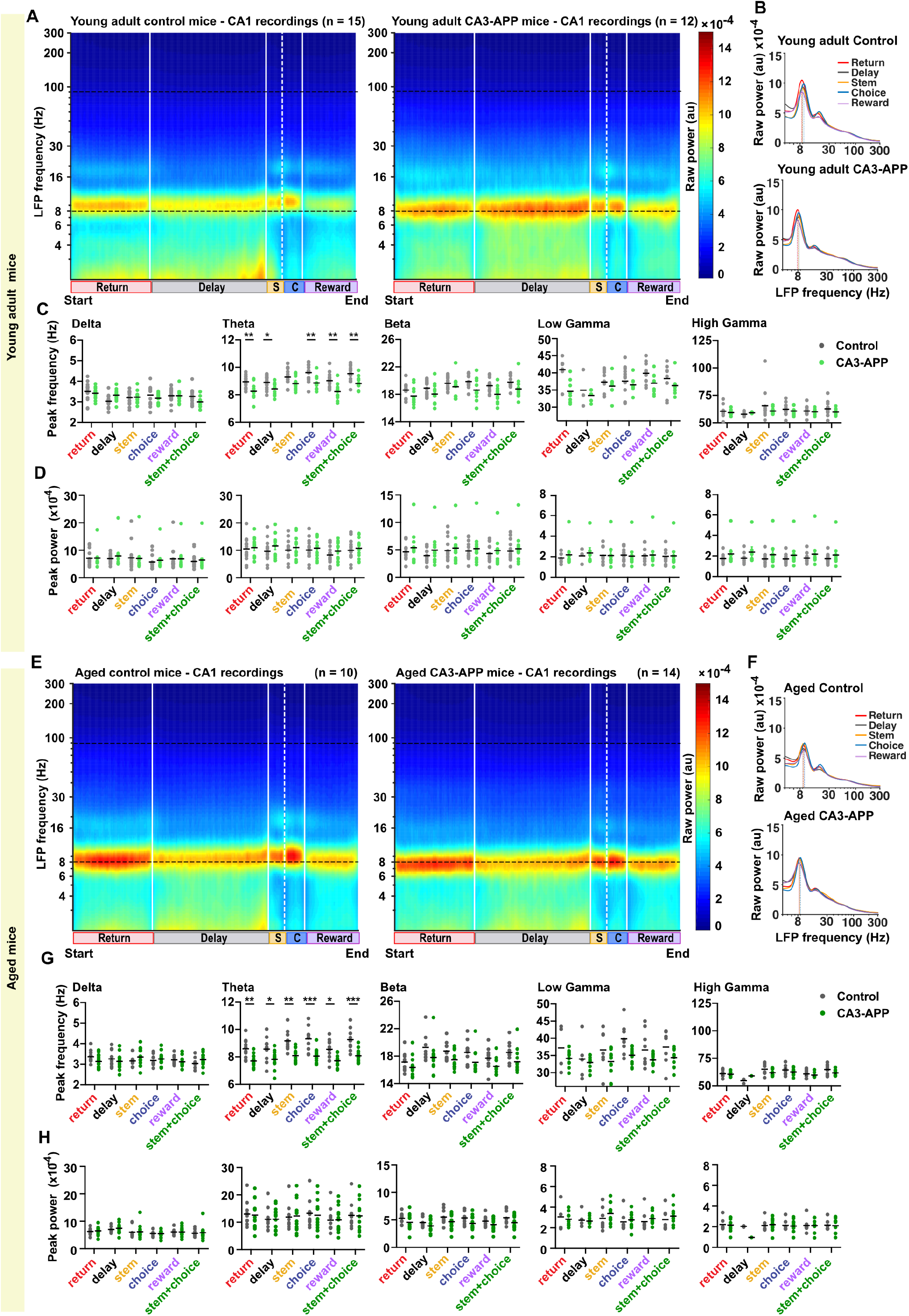
The decreased theta frequency in CA3-APP mice was also detected with quantification of broad-band filtered CA1 LFP signal. **A.** Average spectrogram of raw LFP recorded in CA1 of young adult control and young adult CA3-APP mice. The spectrogram was calculated over the entire lap, and data was first averaged across laps and then across mice. **B.** PSD of the raw LFP recorded from example young adult mice during the figure-8 behavior task (d10 condition), and peak frequency quantification in the theta band. Both genotypes are displayed and quantification was performed for each section of the maze separately. **C.** Quantification of peak frequency of raw LFP signal, by band and by sections of the maze. Theta frequency is reduced in all parts of the maze except stem (p = 0.059) in young adult CA3-APP mice compared to age-matched controls (other regions p < 0.05, MW test adjusted for multiple comparisons by Holm-Šídák method). We found no other differences in the remaining LFP bands that were analyzed, but failed to detect peaks in the PSD for higher frequencies (see **Methods** for more detail on peak detection without normalization). **D.** Quantification of peak power of raw LFP signal, by band and by sections of the maze. No differences in power were detected for any of the LFP bands (p > 0.05, MW test adjusted for multiple comparisons by Holm-Šídák method). **E.** Average spectrogram of raw LFP recorded in CA1 of aged control and aged CA3-APP mice. Data are displayed as in A. **F.** PSD of the raw LFP recorded from example aged mice during the figure-8 behavior task, and peak frequency quantification in the theta band. Data are displayed as described in B. **G.** A reduction in theta oscillation frequency was found in all parts of the maze for aged CA3-APP mice compared to age-matched controls (p < 0.05, MW test adjusted for multiple comparisons by Holm-Šídák method). Data are displayed as in C. **H**. Power did not differ between aged CA3-APP mice and littermate controls for any frequency bands in any of the sections (p > 0.05, MW test adjusted for multiple comparisons by Holm-Šídák method). Data are displayed as in D. See **Supplementary Table 10** for statistics

**Supplementary Table 10.** CA1 LFP analysis of raw data, related to Supplementary Figure 7.

| Fig. panel | Graph description | n= |  | Frequency (Hz) |  | MW test<sup>#</sup> |  |  | Amplitude (10e-4) |  | MW test<sup>#</sup> |  |  |
| --- | --- | --- | --- | --- | --- | --- | --- | --- | --- | --- | --- | --- | --- |
|  |  | ctrl | tg | ctrl* | tg* | U= | P value | Sig. | ctrl* | tg* | U= | P value | Sig. |
| Young adult mice |  |  |  |  |  |  |  |  |  |  |  |  |  |
| Suppl. Fig. 7C,D | CA1 delta |  |  |  |  |  |  |  |  |  |  |  |  |
|  | return | 15 | 12 | 3.5 ± 0.1 | 3.4 ± 0.1 | 74 | 0.83889 | ns | 7.1 ± 0.7 | 7.1 ± 1.0 | 87 | 0.999784 | ns |
|  | Delay | 15 | 12 | 3.0 ± 0.1 | 3.3 ± 0.1 | 38 | 0.059709 | ns | 6.9 ± 0.4 | 8.0 ± 1.3 | 87 | 0.999784 | ns |
|  | Stem | 15 | 12 | 3.2 ± 0.1 | 3.2 ± 0.1 | 87 | 0.956689 | ns | 7.2 ± 1.2 | 7.0 ± 1.4 | 83 | 0.999784 | ns |
|  | choice | 15 | 12 | 3.3 ± 0.1 | 3.2 ± 0.1 | 60 | 0.483604 | ns | 5.7 ± 0.6 | 6.3 ± 1.1 | 84 | 0.999784 | ns |
|  | reward | 15 | 12 | 3.3 ± 0.1 | 3.3 ± 0.1 | 84 | 0.956689 | ns | 6.8 ± 1.1 | 6.9 ± 1.2 | 85 | 0.999784 | ns |
|  | stem+choice | 15 | 12 | 3.3 ± 0.1 | 3.0 ± 0.1 | 50 | 0.237789 | ns | 5.9 ± 0.7 | 6.4 ± 1.3 | 88 | 0.999784 | ns |
| Suppl. Fig. 7C,D | CA1 theta |  |  |  |  |  |  |  |  |  |  |  |  |
|  | Return | 15 | 12 | 8.9 ± 0.1 | 8.3 ± 0.2 | 27 | 0.005766 | ** | 10.5 ± 1.0 | 10.9 ± 1.1 | 86 | 0.958278 | ns |
|  | Delay | 15 | 12 | 8.9 ± 0.1 | 8.4 ± 0.1 | 43 | 0.037946 | * | 9.6 ± 1.0 | 11.6 ± 1.3 | 69 | 0.903599 | ns |
|  | Stem | 15 | 12 | 9.3 ± 0.2 | 8.8 ± 0.1 | 51 | 0.059448 | ns | 10.0 ± 0.9 | 11.0 ± 1.2 | 77 | 0.958278 | ns |
|  | Choice | 15 | 12 | 9.6 ± 0.2 | 8.9 ± 0.1 | 27 | 0.005766 | ** | 10.1 ± 1.0 | 10.8 ± 1.0 | 79 | 0.958278 | ns |
|  | reward | 15 | 12 | 9.0 ± 0.1 | 8.2 ± 0.2 | 25 | 0.005593 | ** | 8.3 ± 0.8 | 9.7 ± 1.0 | 72 | 0.921872 | ns |
|  | stem+choice | 15 | 12 | 9.5 ± 0.1 | 8.8 ± 0.1 | 26 | 0.005766 | ** | 10.1 ± 0.9 | 10.7 ± 1.1 | 79 | 0.958278 | ns |
| Suppl. Fig. 7C,D | CA1 beta |  |  |  |  |  |  |  |  |  |  |  |  |
|  | Return | 14 | 11 | 18.6 ± 0.3 | 17.8 ± 0.4 | 44 | 0.208208 | ns | 4.6 ± 0.4 | 5.4 ± 0.9 | 73 | 0.999505 | ns |
|  | Delay | 14 | 12 | 18.9 ± 0.3 | 18.0 ± 0.4 | 51 | 0.208208 | ns | 4.0 ± 0.3 | 4.9 ± 0.8 | 68 | 0.966381 | ns |
|  | Stem | 14 | 12 | 19.6 ± 0.2 | 19.1 ± 0.4 | 61 | 0.251953 | ns | 4.9 ± 0.6 | 5.3 ± 0.8 | 77 | 0.998867 | ns |
|  | Choice | 14 | 12 | 19.9 ± 0.2 | 18.6 ± 0.4 | 41 | 0.103617 | ns | 4.8 ± 0.4 | 5.2 ± 0.8 | 83 | 0.999505 | ns |
|  | Reward | 15 | 11 | 19.2 ± 0.2 | 18.0 ± 0.4 | 33 | 0.053454 | ns | 4.3 ± 0.4 | 4.9 ± 0.8 | 80 | 0.999505 | ns |
|  | stem+choice | 14 | 12 | 19.8 ± 0.2 | 18.8 ± 0.4 | 34 | 0.053454 | ns | 4.8 ± 0.5 | 5.2 ± 0.8 | 82 | 0.999505 | ns |
| Suppl. Fig. 7C,D | CA1 low gamma |  |  |  |  |  |  |  |  |  |  |  |  |
|  | Return | 6 | 8 | 40.9 ± 1.6 | 34.6 ± 1.2 | 4.5 | 0.050837 | ns | 1.9 ± 0.2 | 3.0 ± 0.8 | 14 | 0.737046 | ns |
|  | Delay | 3 | 6 | 34.9 ± 3.1 | 33.4 ± 0.7 | 8 | 0.904762 | ns | 2.1 ± 0.4 | 3.5 ± 1.2 | 6 | 0.820799 | ns |
|  | Stem | 9 | 8 | 37.3 ± 1.1 | 36.1 ± 1.1 | 28 | 0.795925 | ns | 2.1 ± 0.3 | 3.3 ± 0.8 | 22 | 0.737046 | ns |
|  | Choice | 12 | 9 | 37.6 ± 1.3 | 36.5 ± 0.7 | 42 | 0.795925 | ns | 2.2 ± 0.2 | 3.1 ± 0.8 | 43 | 0.820799 | ns |
|  | Reward | 11 | 10 | 39.9 ± 0.9 | 37.0 ± 0.9 | 22 | 0.094531 | ns | 2.1 ± 0.2 | 2.9 ± 0.7 | 41 | 0.820799 | ns |
|  | stem+choice | 10 | 7 | 38.3 ± 1.3 | 36.3 ± 1.2 | 21 | 0.576365 | ns | 2.0 ± 0.2 | 3.2 ± 0.9 | 25 | 0.820799 | ns |
| Suppl. Fig. 7C,D | CA1 high gamma |  |  |  |  |  |  |  |  |  |  |  |  |
|  | Return | 12 | 9 | 60.5 ± 1.5 | 59.3 ± 0.8 | 41 | 0.718286 | ns | 1.7 ± 0.1 | 2.2 ± 0.4 | 42 | 0.910458 | ns |
|  | Delay | 9 | 7 | 57.8 ± 0.5 | 59.1 ± 0.2 | 13 | 0.270975 | ns | 1.8 ± 0.2 | 2.4 ± 0.5 | 22 | 0.910458 | ns |
|  | Stem | 13 | 11 | 65.5 ± 3.7 | 60.7 ± 0.9 | 51 | 0.686574 | ns | 1.7 ± 0.1 | 2.1 ± 0.4 | 54 | 0.910458 | ns |
|  | Choice | 14 | 11 | 62.3 ± 1.3 | 60.8 ± 1.2 | 59 | 0.718286 | ns | 1.7 ± 0.1 | 2.1 ± 0.3 | 61 | 0.910458 | ns |
|  | Reward | 13 | 11 | 60.7 ± 1.2 | 60.1 ± 0.7 | 65 | 0.73296 | ns | 1.8 ± 0.1 | 2.2 ± 0.4 | 62 | 0.910458 | ns |
|  | stem+choice | 14 | 10 | 62.7 ± 1.6 | 59.8 ± 0.9 | 40 | 0.355572 | ns | 1.7 ± 0.1 | 2.1 ± 0.4 | 60 | 0.910458 | ns |
| Aged mice |  |  |  |  |  |  |  |  |  |  |  |  |  |
| Suppl. Fig. 7G,H | CA1 delta |  |  |  |  |  |  |  |  |  |  |  |  |
|  | Return | 10 | 14 | 3.4 ± 0.1 | 3.1 ± 0.1 | 43 | 0.479029 | ns | 6.3 ± 0.4 | 6.5 ± 0.4 | 60 | 0.991217 | ns |
|  | Delay | 10 | 14 | 3.3 ± 0.1 | 3.1 ± 0.1 | 61 | 0.751326 | ns | 7.0 ± 0.3 | 7.4 ± 0.5 | 68 | 0.999357 | ns |
|  | Stem | 10 | 14 | 3.2 ± 0.1 | 3.4 ± 0.1 | 47 | 0.572781 | ns | 6.0 ± 0.6 | 6.2 ± 0.6 | 59 | 0.991217 | ns |
|  | Choice | 10 | 14 | 3.2 ± 0.1 | 3.3 ± 0.1 | 54 | 0.751326 | ns | 5.5 ± 0.3 | 5.5 ± 0.3 | 66 | 0.999357 | ns |
|  | Reward | 10 | 14 | 3.2 ± 0.1 | 3.1 ± 0.1 | 54 | 0.751326 | ns | 6.1 ± 0.3 | 6.1 ± 0.4 | 67 | 0.999357 | ns |
|  | stem+choice | 10 | 14 | 3.0 ± 0.1 | 3.2 ± 0.1 | 41 | 0.452934 | ns | 5.6 ± 0.4 | 5.8 ± 0.6 | 69 | 0.999357 | ns |
| Suppl. Fig. 7G,H | CA1 theta |  |  |  |  |  |  |  |  |  |  |  |  |
|  | Return | 10 | 14 | 8.6 ± 0.2 | 7.7 ± 0.1 | 20 | 0.007279 | ** | 13.0 ± 1.4 | 12.6 ± 1.4 | 65 | 0.999768 | ns |
|  | Delay | 10 | 14 | 8.6 ± 0.3 | 7.8 ± 0.2 | 33 | 0.030588 | * | 11.1 ± 1.3 | 11.1 ± 1.1 | 67 | 0.99983 | ns |
|  | Stem | 10 | 14 | 9.2 ± 0.3 | 8.1 ± 0.1 | 13 | 0.001492 | ** | 11.9 ± 1.4 | 12.3 ± 1.5 | 69 | 0.99983 | ns |
|  | Choice | 10 | 14 | 9.3 ± 0.3 | 8.1 ± 0.1 | 11 | 0.000989 | *** | 13.2 ± 1.6 | 12.6 ± 1.5 | 64 | 0.999768 | ns |
|  | Reward | 10 | 14 | 8.5 ± 0.2 | 7.7 ± 0.1 | 25 | 0.01445 | * | 10.7 ± 1.4 | 10.8 ± 1.2 | 68 | 0.99983 | ns |
|  | stem+choice | 10 | 14 | 9.3 ± 0.2 | 8.1 ± 0.1 | 9 | 0.000593 | *** | 12.5 ± 1.5 | 12.3 ± 1.5 | 67 | 0.99983 | ns |
| Suppl. Fig. 7G,H | CA1 beta |  |  |  |  |  |  |  |  |  |  |  |  |
|  | Return | 10 | 12 | 17.1 ± 0.4 | 16.3 ± 0.5 | 38 | 0.18618 | ns | 5.3 ± 0.3 | 4.6 ± 0.4 | 39 | 0.548307 | ns |
|  | Delay | 8 | 12 | 19.3 ± 0.7 | 17.8 ± 0.6 | 26 | 0.18618 | ns | 4.5 ± 0.3 | 3.9 ± 0.4 | 32 | 0.548307 | ns |
|  | Stem | 10 | 14 | 18.8 ± 0.4 | 17.5 ± 0.3 | 33 | 0.09823 | ns | 5.5 ± 0.4 | 4.7 ± 0.4 | 47 | 0.548307 | ns |
|  | Choice | 10 | 14 | 18.5 ± 0.5 | 17.1 ± 0.4 | 29 | 0.089669 | ns | 5.4 ± 0.4 | 4.4 ± 0.4 | 38 | 0.328975 | ns |
|  | Reward | 10 | 13 | 17.7 ± 0.5 | 16.5 ± 0.3 | 29 | 0.09823 | ns | 4.8 ± 0.3 | 4.2 ± 0.3 | 47 | 0.548307 | ns |
|  | stem+choice | 10 | 14 | 18.5 ± 0.5 | 17.2 ± 0.4 | 29 | 0.089669 | ns | 5.4 ± 0.4 | 4.5 ± 0.3 | 43 | 0.479029 | ns |
| Suppl. Fig. 7G,H | CA1 low gamma |  |  |  |  |  |  |  |  |  |  |  |  |
|  | Return | 4 | 4 | 37.2 ± 3.4 | 34.2 ± 0.8 | 17 | 0.928853 | ns | 3.1 ± 0.7 | 2.8 ± 0.2 | 19 | 0.996981 | ns |
|  | Delay | 5 | 5 | 34.0 ± 2.7 | 33.0 ± 0.7 | 22 | >0.999999 | ns | 2.7 ± 0.4 | 2.6 ± 0.1 | 22 | >0.999999 | ns |
|  | Stem | 7 | 7 | 36.6 ± 2.2 | 33.3 ± 1.1 | 26 | 0.581236 | ns | 2.9 ± 0.4 | 3.4 ± 0.3 | 30 | 0.874994 | ns |
|  | Choice | 8 | 8 | 39.8 ± 1.6 | 35.1 ± 0.7 | 17 | 0.091471 | ns | 2.6 ± 0.3 | 2.8 ± 0.2 | 35 | 0.874994 | ns |
|  | Reward | 9 | 9 | 36.6 ± 1.7 | 33.7 ± 0.6 | 36 | 0.700799 | ns | 2.6 ± 0.3 | 2.9 ± 0.2 | 24 | 0.293191 | ns |
|  | stem+choice | 7 | 7 | 37.4 ± 2.0 | 34.4 ± 0.8 | 22 | 0.558619 | ns | 2.8 ± 0.4 | 3.1 ± 0.3 | 28 | 0.874994 | ns |
| Suppl. Fig. 7G,H | CA1 high gamma |  |  |  |  |  |  |  |  |  |  |  |  |
|  | Return | 7 | 8 | 60.9 ± 1.3 | 60.4 ± 0.8 | 26 | 0.799223 | ns | 2.2 ± 0.2 | 2.2 ± 0.3 | 28 | >0.999999 | ns |
|  | Delay | 3 | 1 | 54.5 ± 1.3 | 58.9 | n | n too small |  | 2.0 ± 0.1 | 1.0 | n | n too small |  |
|  | Stem | 8 | 10 | 65.0 ± 1.8 | 61.9 ± 0.9 | 23 | 0.544985 | ns | 2.1 ± 0.2 | 2.2 ± 0.2 | 31 | 0.95398 | ns |
|  | Choice | 8 | 10 | 64.4 ± 1.5 | 62.6 ± 1.0 | 30 | 0.792783 | ns | 2.1 ± 0.2 | 2.1 ± 0.2 | 38 | 0.998899 | ns |
|  | Reward | 8 | 8 | 60.7 ± 1.2 | 59.7 ± 0.8 | 25 | 0.792783 | ns | 2.1 ± 0.2 | 2.1 ± 0.2 | 26 | 0.966985 | ns |
|  | stem+choice | 8 | 9 | 64.7 ± 1.7 | 61.7 ± 0.9 | 23 | 0.607725 | ns | 2.2 ± 0.2 | 2.1 ± 0.2 | 35 | 0.998899 | ns |
| * mean + SEM # adjusted for multiple comparisons by Holm-Sidak method |  |  |  |  |  |  |  |  |  |  |  |  |  |

\* mean ± SEM # adjusted for multiple comparisons by Holm-Šidák method

**Supplementary Figure 8.**
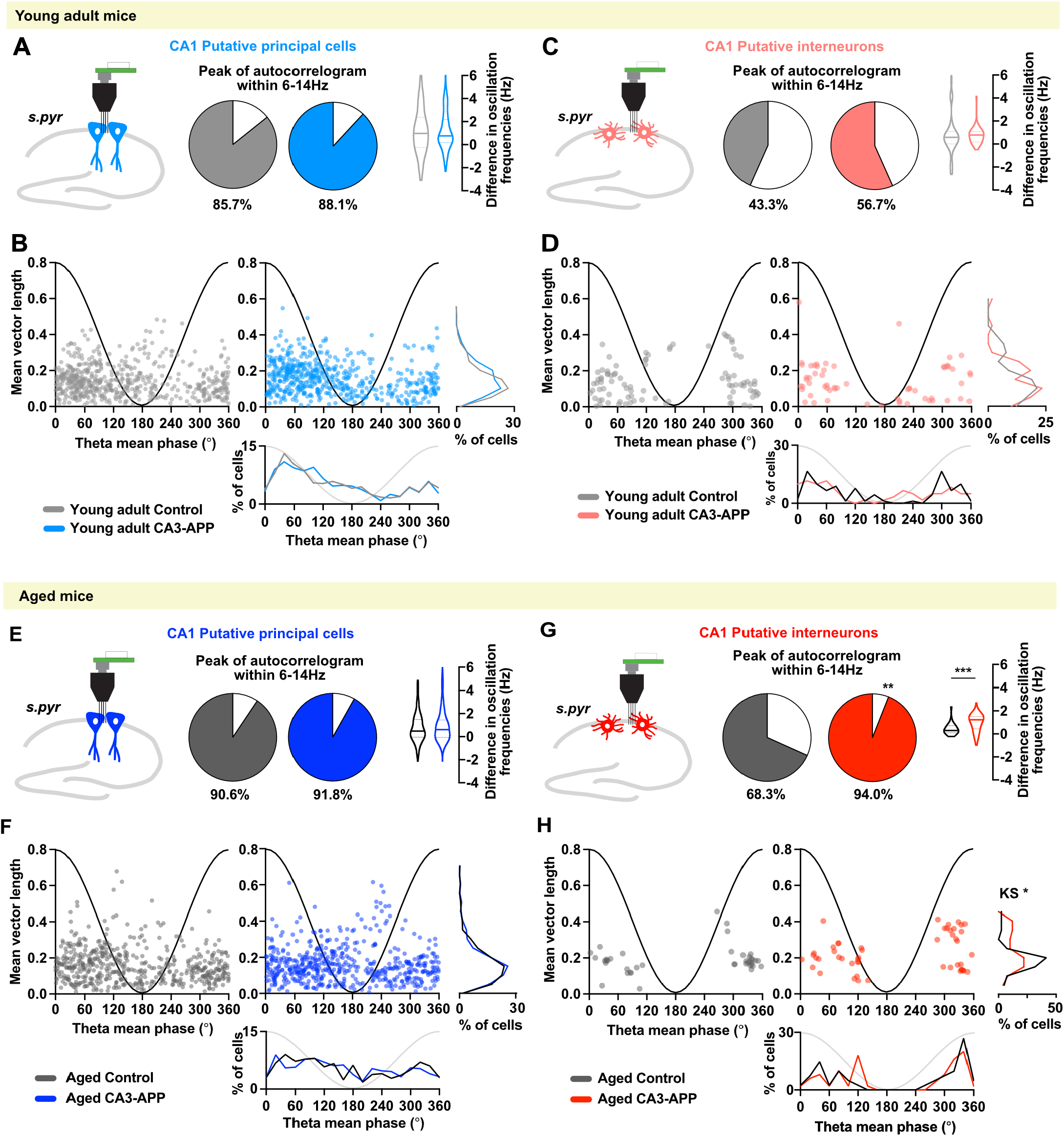
Temporal firing patterns of CA1 principal cells and interneurons. **A.** Spike oscillation frequencies in the theta range were obtained by performing the spike-time autocorrelation for each neuron and by using a fast-Fourier transform to obtain the predominant frequency (peak of autocorrelogram). The fraction of cells with detectable peaks in the theta range (6-14 Hz) is reported as the colored portion in the pie charts (p = 0.25, Chisquare test). For cells oscillating in the theta range, we then calculated the differences between the predominant spike oscillation frequency and the predominant LFP oscillation frequency in the same recording session. The LFP frequency and spike oscillation frequencies in the theta range were concurrently reduced by APP expression, such that the difference between the two oscillations did not change in young adult CA3-APP mice compared to age-matched controls (control 1.2 ± 0.1 Hz, CA3-APP 1.2 ± 0.1 Hz, p = 0.81, MW test). **B.** The preferred LFP phase when cells spiked (mean phase) is plotted against the amplitude of theta phase locking (mean vector length) for CA1 principal cells from control (grey) and CA3-APP mice (blue). Phase preference (control 142 ± 5°, CA3-APP 140 ± 5°, p = 0.93, MW test) or phase locking (control 0.15 ± 0.01, CA3-APP 0.16 ± 0.01, p = 0.056, MW test) of CA1 principal cells did not differ between young adult CA3-APP mice and age-matched control mice. **C.** The difference between the spike oscillation frequency of CA1 interneurons compared to the frequency of LFP oscillations did not change between young adult CA3-APP mice and age-matched control mice (p = 0.11, Chi-square test; control 0.9 ± 0.3 Hz, CA3-APP 0.8 ± 0.1 Hz, p = 0.33, MW test). Data are displayed as in A. **D.** Phase preference (control 165 ± 13°, CA3-APP 154 ± 16°, p = 0.40) or phase locking (control 0.15 ± 0.11, CA3-APP 0.14 ± 0.01, p = 0.54, MW test) of CA1 interneurons did not differ between young adult CA3-APP mice and age-matched controls. Data are displayed as in B. **E.** The difference between principal cell spike oscillation frequency and LFP oscillation frequency did not change in aged CA3-APP mice compared to age-matched controls (p = 0.48, Chi-square test; control 0.9 ± 0.1 Hz, CA3-APP 0.9 ± 0.1 Hz, p = 0.46, MW test). Data are displayed as in A. **F.** Phase preference (control 165 ± 13°, CA3-APP 154 ± 16°, p = 0.90, MW test) or phase locking of CA1 principal cells (control 0.15 ± 0.01, CA3-APP 0.14 ± 0.01, p = 0.96, MW test) did not differ between aged CA3-APP mice and age-matched control mice. Data are displayed as in B. **G.** CA1 interneurons from aged CA3-APP mice have a higher difference between spike oscillation frequency and LFP oscillation frequency than aged control mice, which implies that the reduction in LFP frequency was more pronounced than the interneurons’ spiking frequency (p = 0.0013, Chi-square test; control 0.4 ± 0.1, CA3-APP 1.0 ± 0.1, p = 0.0003, MW test). Data are displayed as in C. **H.** While the mean phase locking of CA1 interneurons in aged CA3-APP mice was not statistically different from littermate controls (control 206 ± 22°, CA3-APP 203 ±18°, p = 0.97, MW test), the distribution of the mean vector length in CA3-APP mice showed a biphasic distribution with a higher fraction of cells that were strongly theta modulated (control 0.19 ± 0.01, CA3-APP 0.23 ± 0.01, p = 0.068, MW test; p = 0.019 KS test of distribution). Data are displayed as in D. See **Supplementary Table 11** for statistics.

**Supplementary Table 11.**
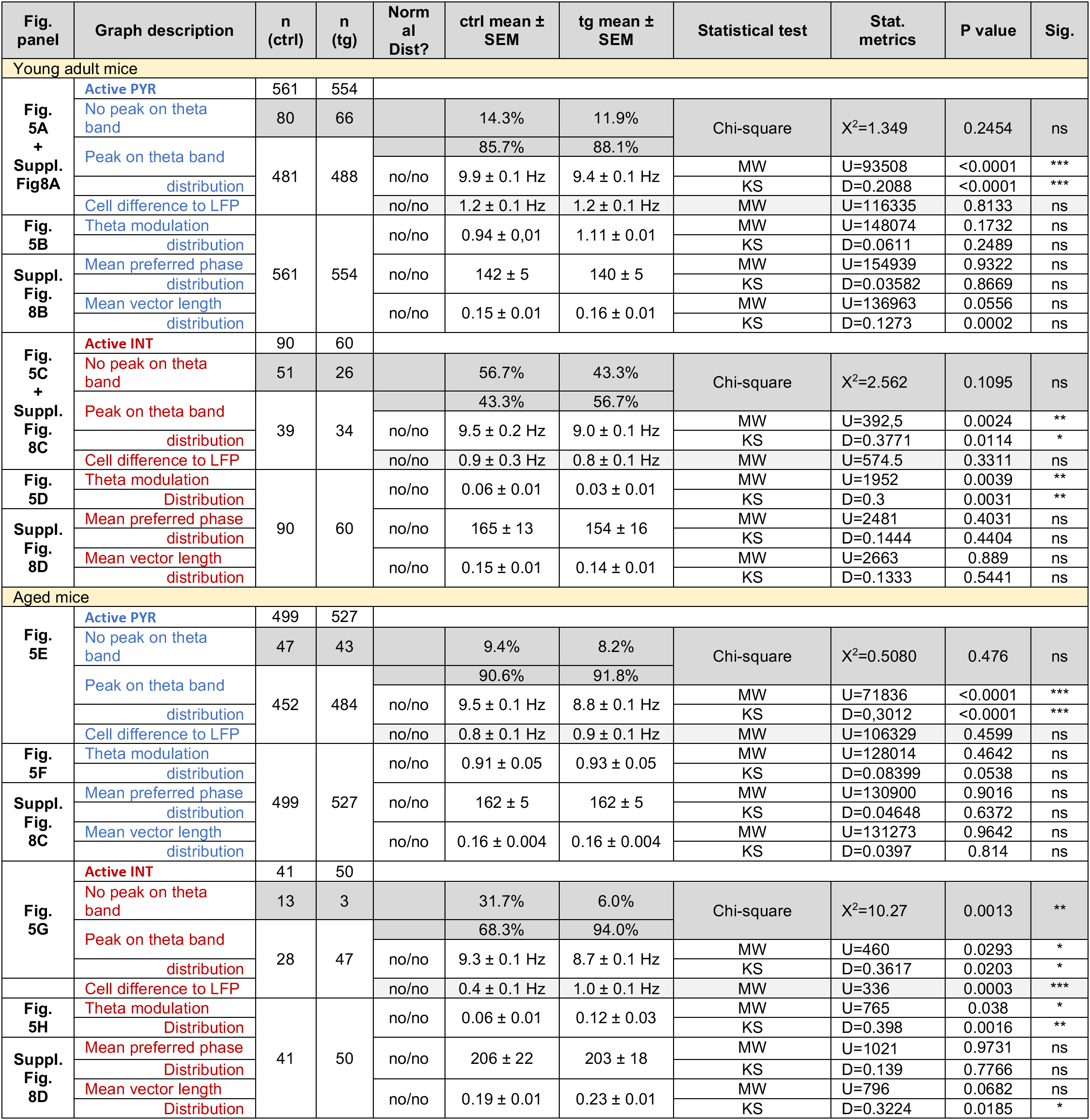
Theta modulation of CA1 single units, related to Figure 5 and Supplementary Figure 8.

**Supplementary Figure 9.**
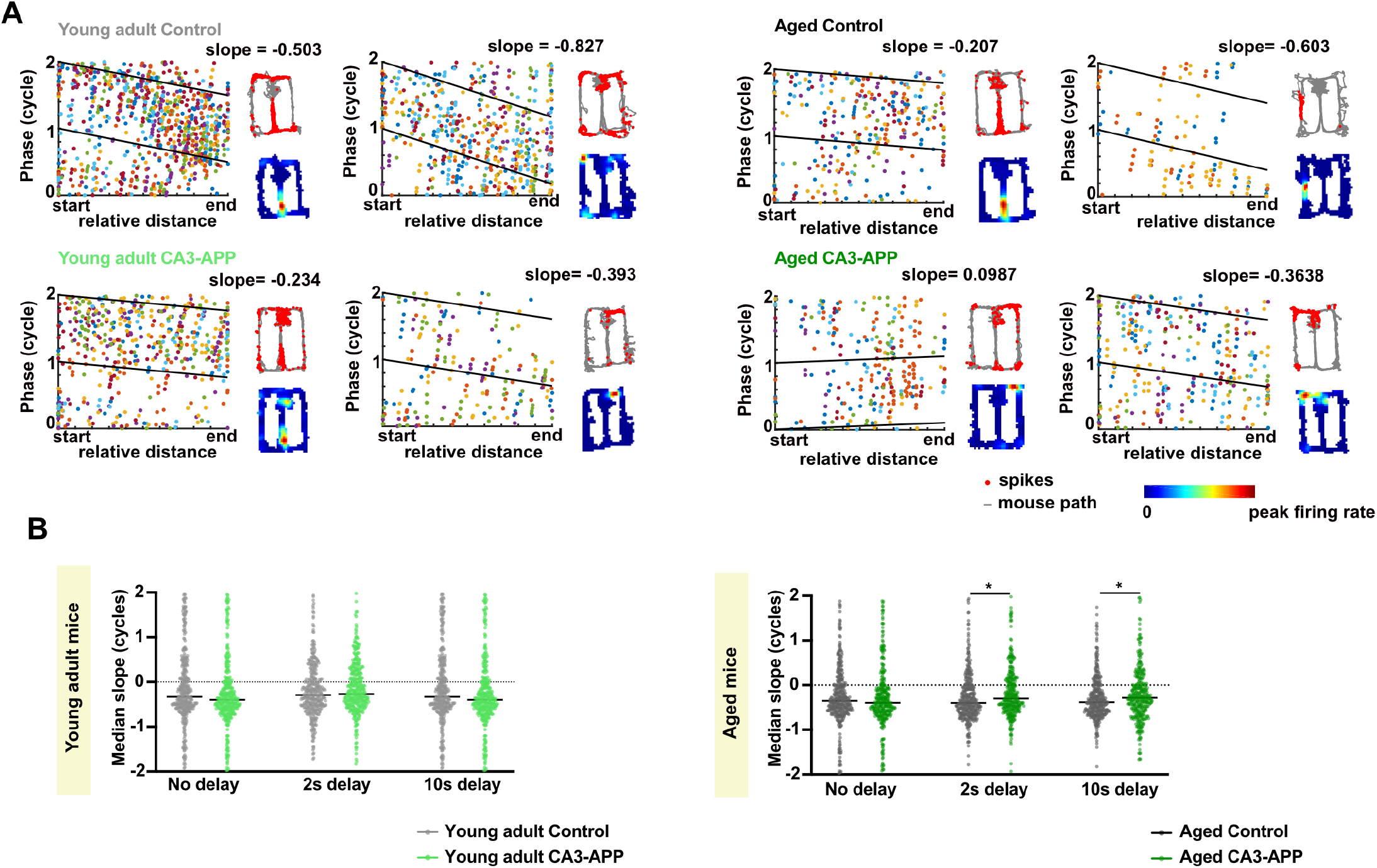
Phase precession of CA1 cells in different delay conditions in the figure-8 task. **A.** Examples of average phase precession slopes obtained by analysis of phase precession using trains of spikes (each spike train has a different color). Data from 10-s delay condition. **B.** The median phase precession slope per cell and delay condition was calculated by using the slopes obtained for each spike train of the cell (significant or not). Comparison between genotypes was performed using the cells’ median slopes during the continuous version of the figure-8 task (no delay in the central arm) as well as during the delayed versions of the task (2-s and 10-s delays in the central arm). Phase precession was not different from control levels in young adult CA3-APP mice (p = 0.057 for no delay, p = 0.094 for 2-s delay, p = 0.97 for 10-s delay, MW test) and was significantly reduced in aged CA3-APP mice only when there was a working memory component (2-s delay and 10-s delay) to the task (no delay p = 0.71, 2-s delay p = 0.049, 10-s delay p = 0.028 for 10-s delay, MW test). See **Supplementary Table 12** for statistics.

**Supplementary Figure 10.**
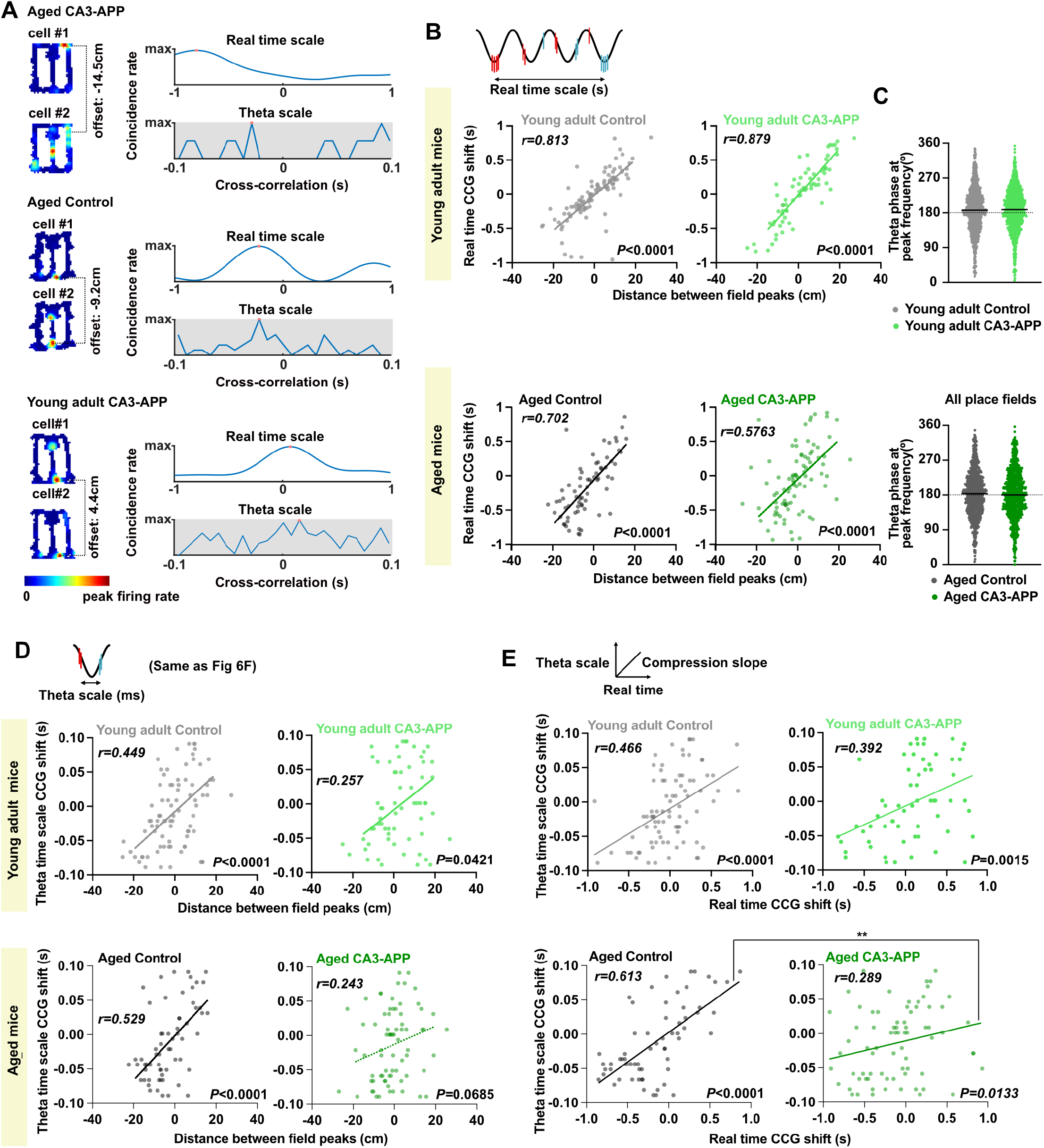
Compression of theta sequences was impaired in CA1 place cells recorded from aged CA3-APP mice. **A.** Examples of pairs of overlapping place fields in the figure-8 task (10-s delay). Color-coded place maps of both cells in the pair are depicted to the left. **B.** Correlation of the distance between place field peaks/center of mass (linear distance in cm) and the time shift between peak firing rates at the behavioral scale. Place fields in the delay region were excluded from the analysis. As expected, correlations between distance and time are high because it requires more time to run between more distant fields (p < 0.0001, Spearman correlation). **C.** Theta phase at the peak firing rate of each place field did not differ between CA3-APP mice at either age and their respective controls (young adult: control −0.17 ± 0.05, CA3-APP −0.19 ± 0.05, p = 0.6192; aged: control −0.44 ± 0.04, CA3-APP −0.26 ± 0.05, p = 0.0019, MW test). **D.** Same as Figure 6F. Correlation between distance between place field peaks/center of mass (linear distance in cm) and time shift within a theta cycle (control young and aged p < 0.0001; CA3-APP young p = 0.042, aged p = 0.069, Spearman correlation). **E.** Given the high correlation between place field distance and time to run between place field peaks, similar results as with distance (see D and Fig. 6F) are obtained when comparing real time to theta-time scale. With the comparison between two-time differences (x axis, real time shift; y axis, theta time shift), the compression of temporal sequences can be estimated by the slope of the regression line. Compression was maintained in young adult CA3-APP mice (n = 83 overlapping fields for control and n = 63 for CA3-APP, p = 0.40, ANCOVA test) but aged CA3-APP mice had reduced compression slopes in comparison to aged-matched controls (n = 63 overlapping fields for control and n = 73 for CA3-APP, p = 0.0035, ANCOVA test). See **Supplementary Table 12** for statistics.

**Supplementary Table 12.**
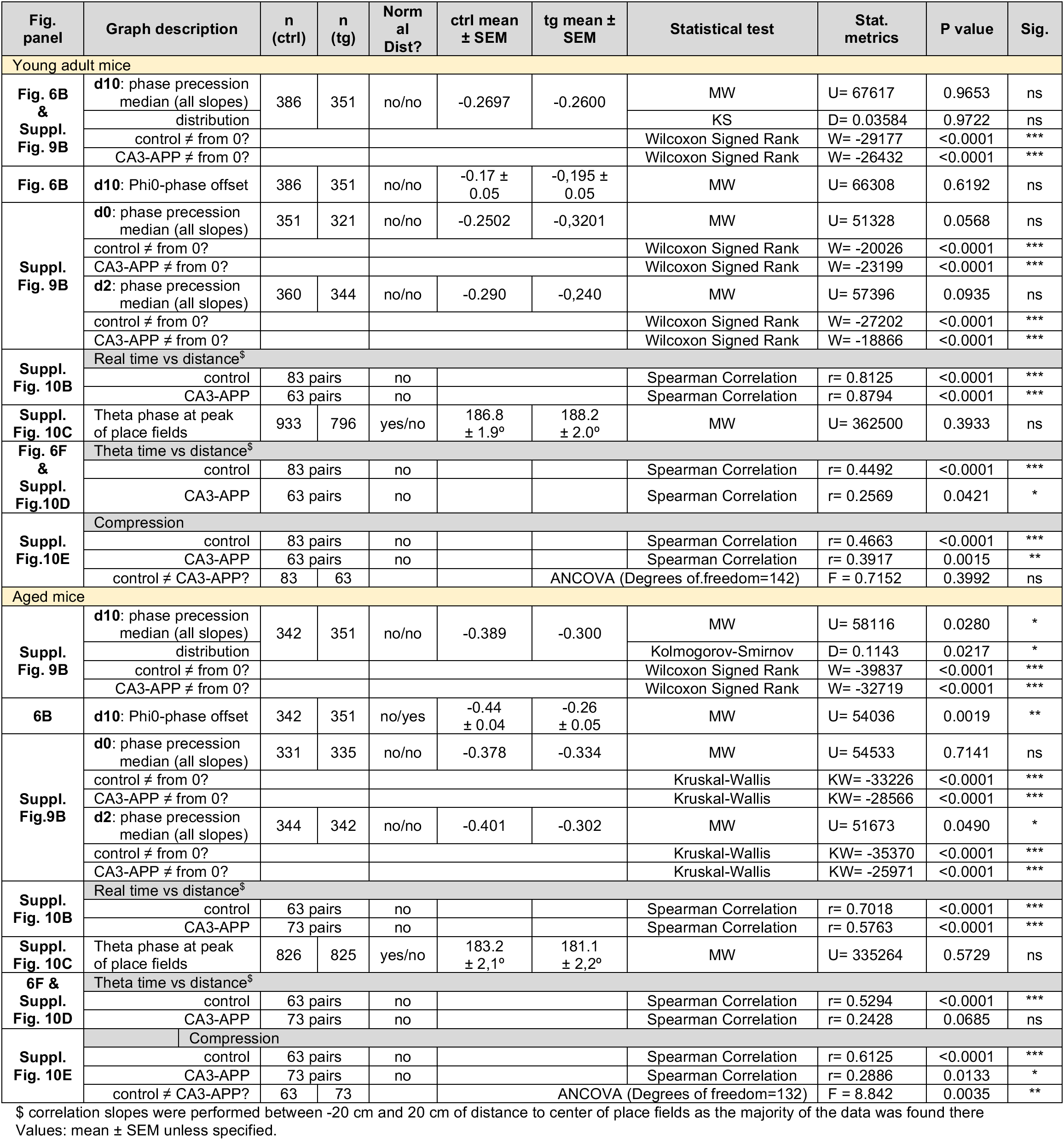
Quantification of phase precession metrics for CA1 place cells.

**Supplementary Figure 11.**
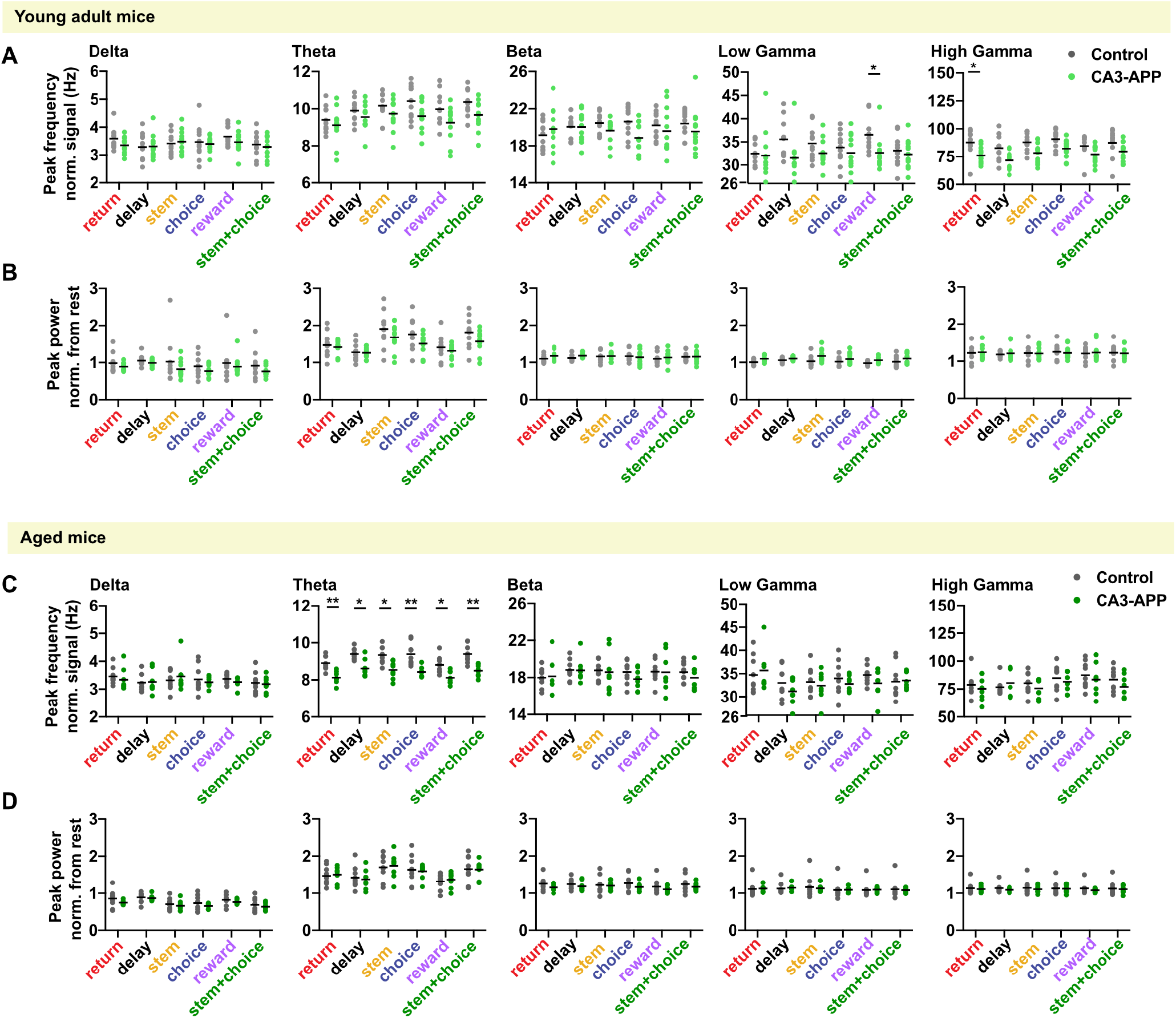
APP expression in CA3 neurons reduced theta oscillation frequency at CA3/DG recording sites in aged, but not in young adult mice. **A.** Quantification of peak frequency of normalized LFP signal, by band and by section of the maze. In the low gamma frequency band, a frequency reduction was observed in only the reward section of the maze for young adult CA3-APP mice compared to age-matched controls. No differences were observed for delta, theta, beta and high gamma frequencies at this age (p > 0.05, MW test adjusted for multiple comparisons by Holm-Šídák method). **B.** Quantification of peak power of normalized LFP signal, by band and by sub-section of the maze. No differences in power were detected in any of the LFP bands (p > 0.05, MW test adjusted for multiple comparisons by Holm-Šídák method). **C.** A reduction in theta oscillation frequency was found in all sections of the maze for aged CA3-APP mice, compared to aged matched-controls (p < 0.05, MW test adjusted for multiple comparisons by Holm-Šídák method). No differences to age-matched control mice were detected in other frequency bands of aged CA3-APP mice (p > 0.05, MW test adjusted for multiple comparisons by Holm-Šídák method). Data are displayed as in A. **D.** No differences in power were detected between aged CA3-APP mice and age-matched controls (p > 0.05, MW test adjusted for multiple comparisons by Holm-Šídák method). Data are displayed as in B. See **Supplementary Table 13** for statistics. Figure corresponds to **Supplementary Figure 6** for CA1 data.

**Supplementary Table 13.**
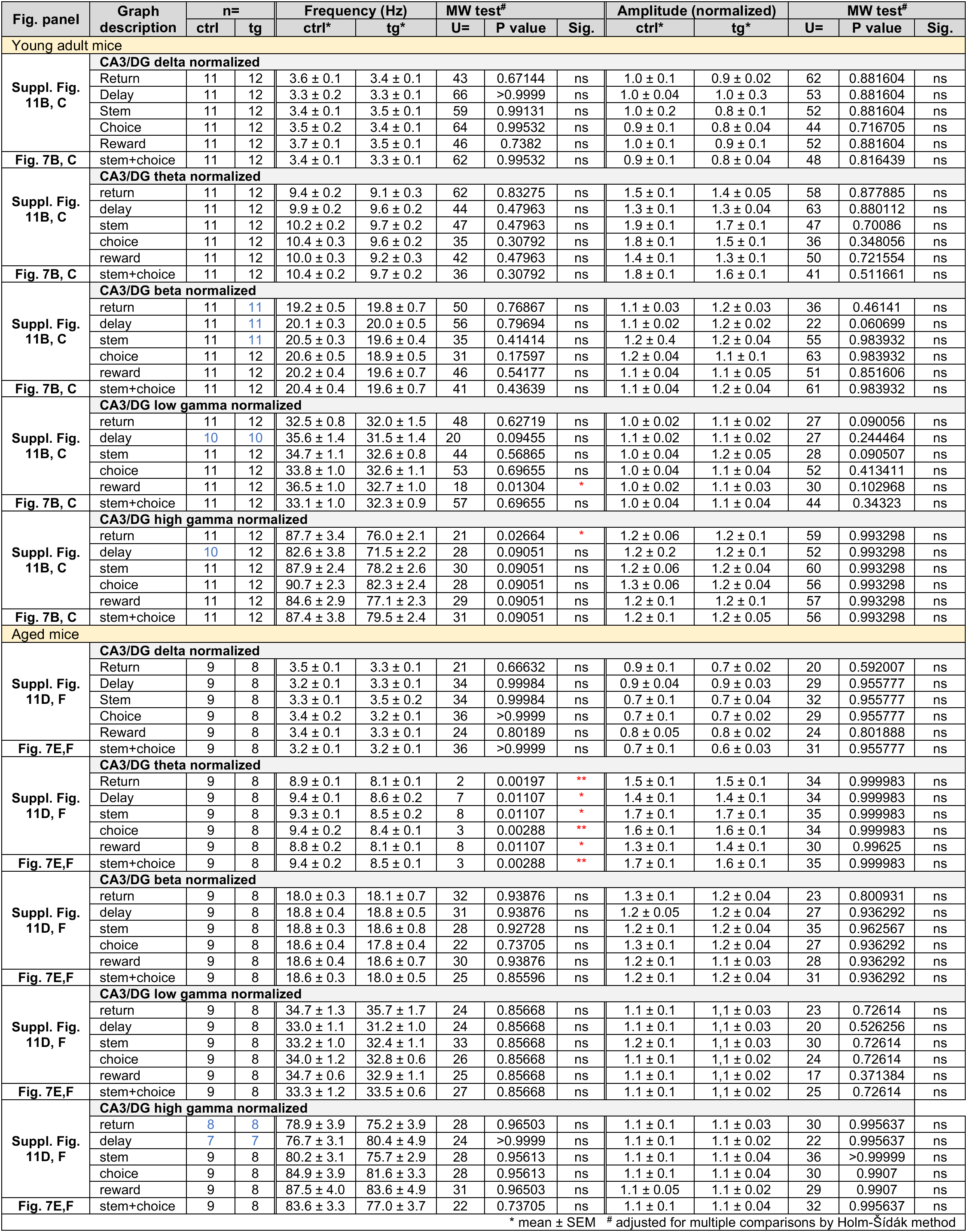
Data and statistics for Figure 7 and Supplementary Figure 11.

**Supplementary Figure 12.**
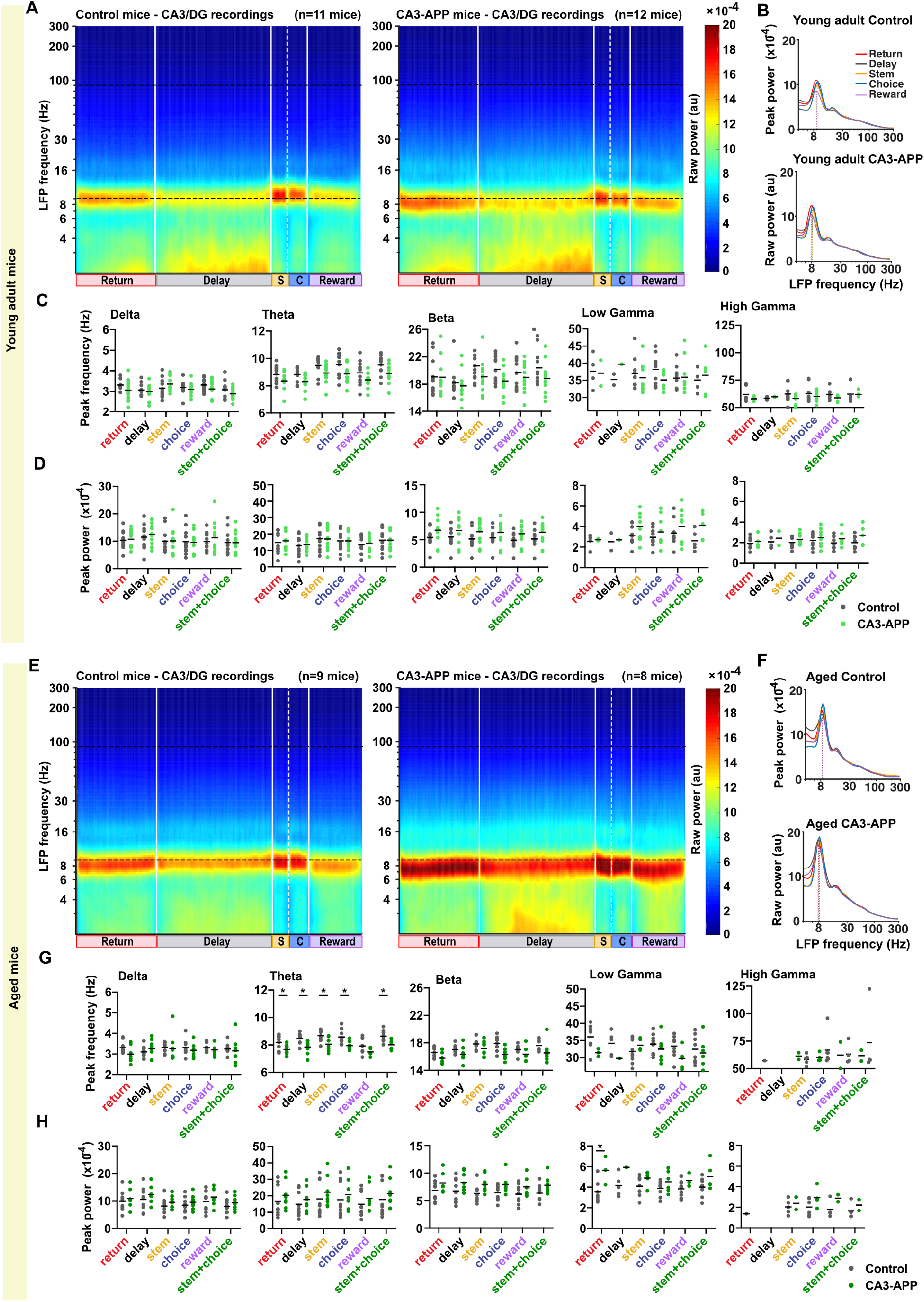
The reduction in theta frequency at CA3/DG recording sites in aged CA3-APP mice was also detected with quantification of broad-band filtered LFP. **A.** Average Spectrogram of raw LFP recorded at CA3/DG sites in young adult control and CA3-APP mice. The spectrogram was calculated over the entire lap, and data was first averaged across laps and then across mice. **B.** PSD of the raw LFP recorded from example young adult mice during the figure-8 behavior task (d10 condition), and peak frequency quantification in the theta band. Both genotypes are displayed and quantification was performed for each section of the maze separately. **C.** Quantification of peak frequency of raw CA3/DG LFP signal, by band and by sections of the maze. No differences in any of the LFP bands were found between young adult CA3-APP mice and age-matched controls (p > 0.05, MW test adjusted for multiple comparisons by Holm-Šídák method). Of note, we failed to detect peaks in the PSD for many of the higher LFP frequencies (see **Supplementary Table 14** for statistics and **Methods** for peak detection without normalization). **D.** Quantification of peak power of raw CA3/DG LFP signal, by band and by sections of the maze. No differences in power were detected for any of the LFP bands (p > 0.05, MW test adjusted for multiple comparisons by Holm-Šídák method). **E.** Same panel as in A, but for aged mice **F.** PSD of the raw LFP recorded from example aged mice during the figure-8 behavior task, and peak frequency quantification in the theta band. Data are displayed as described in B. **G.** A reduction in theta oscillation frequency was found in all parts of the maze for aged CA3-APP mice compared to age-matched controls except in sections where mice were mostly immobile (reward, p = 0.074). No differences in frequency were found for any of the other LFP bands (p > 0.05, MW test adjusted for multiple comparisons by Holm-Šídák method). Data are displayed as in C. **H.** No changes in power were detected in CA3-APP mice compared to littermate controls (p > 0.05, MW test adjusted for multiple comparisons by Holm-Šídák method). Data are displayed as in D. See **Supplementary Table 14** for all statistical analysis. Figure corresponds to **Supplementary Figure 7** for CA1 data.

**Supplementary Table 14.** Data and statistics for Supplementary Figure 12.

| Fig. panel | Graph description | n= |  | Frequency (Hz) |  | MW test<sup>#</sup> |  |  | Amplitude (10e-4) |  | MW test<sup>#</sup> |  |  |
| --- | --- | --- | --- | --- | --- | --- | --- | --- | --- | --- | --- | --- | --- |
|  |  | ctrl | tg | ctrl* | tg* | U= | P value | Sig. | ctrl* | tg* | U= | P value | Sig. |
| Young adult mice |  |  |  |  |  |  |  |  |  |  |  |  |  |
| Suppl. Fig. 12C, D | CA3/DG delta |  |  |  |  |  |  |  |  |  |  |  |  |
|  | return | 11 | 12 | 3.3 ± 0.1 | 3.1 ± 0.1 | 45 | 0.6952 | ns | 10.3 ± 1.0 | 10.8 ± 0.9 | 63 | 0.999869 | ns |
|  | delay | 11 | 12 | 3.1 ± 0.1 | 3.0 ± 0.1 | 62 | 0.83275 | ns | 11.5 ± 1.3 | 12.5 ± 1.2 | 57 | 0.996345 | ns |
|  | stem | 11 | 12 | 3.2 ± 0.1 | 3.4 ± 0.1 | 40 | 0.529 | ns | 10.0 ± 1.3 | 10.1 ± 1.3 | 62 | 0.999869 | ns |
|  | choice | 11 | 12 | 3.2 ± 0.1 | 3.1 ± 0.1 | 55 | 0.79816 | ns | 9.9 ± 1.4 | 9.6 ± 1.2 | 63 | 0.999869 | ns |
|  | reward | 11 | 12 | 3.3 ± 0.1 | 3.1 ± 0.1 | 46 | 0.6952 | ns | 9.9 ± 1.0 | 11.3 ± 1.6 | 62 | 0.999869 | ns |
|  | stem+choice | 11 | 12 | 3.1 ± 0.1 | 2.9 ± 0.1 | 52 | 0.79816 | ns | 9.6 ± 1.3 | 9.5 ± 1.1 | 65 | 0.999869 | ns |
| Suppl. Fig. 12C, D | CA3/DG theta |  |  |  |  |  |  |  |  |  |  |  |  |
|  | Return | 11 | 12 | 8.8 ± 0.2 | 8.4 ± 0.2 | 42 | 0.24926 | ns | 14.9 ± 2.0 | 15.9 ± 1.6 | 62 | 0.999904 | ns |
|  | Delay | 11 | 12 | 8.8 ± 0.2 | 8.3 ± 0.2 | 36 | 0.24813 | ns | 12.9 ± 1.8 | 13.5 ± 1.2 | 64 | 0.999904 | ns |
|  | Stem | 11 | 12 | 9.5 ± 0.2 | 8.9 ± 0.2 | 30 | 0.1504 | ns | 17.2 ± 2.1 | 17.0 ± 1.9 | 65 | 0.999904 | ns |
|  | Choice | 11 | 12 | 9.6 ± 0.3 | 8.9 ± 0.2 | 37 | 0.24813 | ns | 15.8 ± 2.0 | 15.8 ± 1.6 | 61 | 0.999904 | ns |
|  | Reward | 11 | 12 | 9.0 ± 0.3 | 8.4 ± 0.2 | 41 | 0.24926 | ns | 13.6 ± 1.7 | 14.3 ± 1.4 | 64 | 0.999904 | ns |
|  | stem+choice | 11 | 12 | 9.5 ± 0.2 | 8.9 ± 0.2 | 33 | 0.20097 | ns | 16.3 ± 2.0 | 16.2 ± 1.7 | 62 | 0.999904 | ns |
| Suppl. Fig. 12C, D | CA3/DG beta |  |  |  |  |  |  |  |  |  |  |  |  |
|  | Return | 11 | 11 | 19.1 ± 0.8 | 19.0 ± 0.8 | 58 | 0.93728 | ns | 5.4 ± 0.5 | 6.7 ± 0.7 | 35 | 0.41414 | ns |
|  | Delay | 9 | 11 | 18.2 ± 0.8 | 17.8 ± 0.7 | 42 | 0.93728 | ns | 5.5 ± 0.6 | 6.7 ± 0.7 | 31 | 0.41414 | ns |
|  | Stem | 11 | 12 | 20.7 ± 0.7 | 19.1 ± 0.7 | 40 | 0.46603 | ns | 5.1 ± 0.5 | 6.5 ± 0.6 | 40 | 0.41414 | ns |
|  | Choice | 11 | 12 | 20.2 ± 0.6 | 18.5 ± 0.5 | 35 | 0.30792 | ns | 5.3 ± 0.5 | 6.3 ± 0.6 | 42 | 0.41414 | ns |
|  | Reward | 11 | 11 | 19.7 ± 0.7 | 19.0 ± 0.6 | 52 | 0.93728 | ns | 4.9 ± 0.5 | 6.1 ± 0.5 | 32 | 0.332579 | ns |
|  | stem+choice | 11 | 12 | 20.4 ± 0.8 | 18.8 ± 0.7 | 44 | 0.56865 | ns | 5.1 ± 0.5 | 6.3 ± 0.6 | 40 | 0.41414 | ns |
| Suppl. Fig. 12C, D | CA3/DG low gamma |  |  |  |  |  |  |  |  |  |  |  |  |
|  | Return | 4 | 2 | 37.7 ± 2.5 | 37.1 ± 3.8 | 4 | >0.999999 | ns | 2.5 ± 0.4 | 2.7 ± 0.1 | 3 | 0.906239 | ns |
|  | Delay | 2 | 1 | 35.2 ± 1.6 | 39.7 |  |  | x | 2.5 ± 0.9 | 2.7 |  |  | x |
|  | Stem | 10 | 8 | 37.0 ± 1.4 | 35.8 ± 1.6 | 31 | 0.91482 | ns | 3.2 ± 0.3 | 4.0 ± 0.5 | 24 | 0.53182 | ns |
|  | Choice | 9 | 9 | 38.2 ± 1.5 | 35.1 ± 0.9 | 22 | 0.44003 | ns | 3.0 ± 0.3 | 3.5 ± 0.5 | 33 | 0.906239 | ns |
|  | Reward | 10 | 10 | 35.7 ± 1.1 | 35.9 ± 1.6 | 49 | 0.99991 | ns | 3.4 ± 0.4 | 4.0 ± 0.5 | 44 | 0.906239 | ns |
|  | stem+choice | 5 | 5 | 35.1 ± 1.4 | 36.5 ± 2.7 | 12 | >0.999999 | ns | 2.6 ± 0.4 | 4.1 ± 0.6 | 4 | 0.393722 | ns |
| Suppl. Fig. 12C, D | CA3/DG high gamma |  |  |  |  |  |  |  |  |  |  |  |  |
|  | return | 7 | 4 | 61.8 ± 2.4 | 58.0 ± 1.3 | 11 | 0.87684 | ns | 1.9 ± 0.2 | 2.1 ± 0.3 | 11 | 0.785964 | ns |
|  | delay | 5 | 2 | 58.4 ± 0.5 | 59.8 ± 0.3 | 1 | 0.70511 | x | 2.1 ± 0.3 | 2.4 ± 0.7 | 4 | 0.865714 | ns |
|  | stem | 9 | 9 | 62.4 ± 2.6 | 58.3 ± 1.4 | 33 | 0.87684 | ns | 2.0 ± 0.2 | 2.3 ± 0.2 | 25 | 0.57646 | ns |
|  | choice | 9 | 9 | 62.7 ± 2.6 | 60.3 ± 1.5 | 40 | 0.94924 | ns | 2.2 ± 0.2 | 2.5 ± 0.3 | 33 | 0.785964 | ns |
|  | reward | 7 | 6 | 61.9 ± 2.0 | 58.8 ± 1.1 | 14 | 0.8384 | ns | 2.0 ± 0.2 | 2.4 ± 0.3 | 14 | 0.739254 | ns |
|  | stem+choice | 8 | 5 | 62.2 ± 3.0 | 61.9 ± 1.3 | 11 | 0.70511 | ns | 2.0 ± 0.2 | 2.7 ± 0.4 | 9 | 0.57646 | ns |
| Aged mice |  |  |  |  |  |  |  |  |  |  |  |  |  |
| Suppl. Fig. 12G, H | CA3/DG delta |  |  |  |  |  |  |  |  |  |  |  |  |
|  | Return | 9 | 8 | 3.3 ± 0.1 | 3.0 ± 0.1 | 15 | 0.24804 | ns | 10.0 ± 1.2 | 10.9 ± 1.2 | 31 | 0.855956 | ns |
|  | Delay | 9 | 8 | 3.1 ± 0.1 | 3.3 ± 0.2 | 27 | 0.84285 | ns | 10.6 ± 1.4 | 12.3 ± 1.1 | 26 | 0.855956 | ns |
|  | Stem | 9 | 8 | 3.3 ± 0.1 | 3.3 ± 0.2 | 23 | 0.73948 | ns | 8.1 ± 1.0 | 9.6 ± 0.8 | 25 | 0.855956 | ns |
|  | Choice | 9 | 8 | 3.3 ± 0.1 | 3.2 ± 0.1 | 30 | 0.84464 | ns | 8.4 ± 1.0 | 9.6 ± 0.9 | 27 | 0.855956 | ns |
|  | Reward | 9 | 8 | 3.3 ± 0.1 | 3.2 ± 0.1 | 26 | 0.84285 | ns | 9.7 ± 1.2 | 11.4 ± 1.1 | 29 | 0.855956 | ns |
|  | stem+choice | 9 | 8 | 3.2 ± 0.1 | 3.1 ± 0.2 | 31 | 0.84464 | ns | 8.0 ± 0.9 | 9.4 ± 0.9 | 23 | 0.800931 | ns |
| Suppl. Fig. 12G, H | CA3/DG theta |  |  |  |  |  |  |  |  |  |  |  |  |
|  | Return | 9 | 8 | 8.2 ± 0.2 | 7.7 ± 0.1 | 13 | 0.04504 | * | 16.7 ± 3.2 | 20.3 ± 3.0 | 23 | 0.737046 | ns |
|  | Delay | 9 | 8 | 8.5 ± 0.1 | 7.8 ± 0.2 | 9 | 0.03887 | * | 14.8 ± 2.5 | 17.7 ± 2.5 | 25 | 0.737046 | ns |
|  | Stem | 9 | 8 | 8.7 ± 0.1 | 8.1 ± 0.2 | 10 | 0.04369 | * | 18.1 ± 3.5 | 22.2 ± 3.2 | 22 | 0.737046 | ns |
|  | Choice | 9 | 8 | 8.6 ± 0.2 | 8.0 ± 0.1 | 6 | 0.01472 | * | 17.6 ± 3.6 | 20.8 ± 3.2 | 25 | 0.737046 | ns |
|  | Reward | 9 | 8 | 7.9 ± 0.2 | 7.5 ± 0.1 | 17 | 0.07446 | ns | 14.8 ± 2.5 | 18.5 ± 2.7 | 22 | 0.737046 | ns |
|  | stem+choice | 9 | 8 | 8.6 ± 0.1 | 8.0 ± 0.1 | 10 | 0.04369 | * | 17.7 ± 3.5 | 21.2 ± 3.2 | 24 | 0.737046 | ns |
| Suppl. Fig. 12G, H | CA3/DG beta |  |  |  |  |  |  |  |  |  |  |  |  |
|  | Return | 9 | 7 | 16.6 ± 0.3 | 15.8 ± 0.3 | 13 | 0.21145 | ns | 6.8 ± 0.6 | 8.2 ± 0.7 | 21 | 0.48721 | ns |
|  | Delay | 9 | 8 | 17.0 ± 0.2 | 16.3 ± 0.4 | 18 | 0.25316 | ns | 6.7 ± 0.7 | 8.3 ± 0.6 | 22 | 0.48721 | ns |
|  | Stem | 9 | 8 | 17.8 ± 0.3 | 17.2 ± 0.4 | 23 | 0.25316 | ns | 6.3 ± 0.5 | 8.0 ± 0.6 | 15 | 0.248038 | ns |
|  | Choice | 9 | 8 | 17.9 ± 0.4 | 16.3 ± 0.3 | 8 | 0.03262 | ns | 6.4 ± 0.5 | 7.9 ± 0.6 | 18 | 0.385235 | ns |
|  | Reward | 9 | 7 | 17.0 ± 0.2 | 16.3 ± 0.4 | 16 | 0.25316 | ns | 6.2 ± 0.6 | 7.5 ± 0.6 | 20 | 0.48721 | ns |
|  | stem+choice | 9 | 8 | 17.6 ± 0.3 | 16.5 ± 0.6 | 15 | 0.21145 | ns | 6.4 ± 0.5 | 7.8 ± 0.5 | 19 | 0.385235 | ns |
| Suppl. Fig. 12G, H | CA3/DG low gamma |  |  |  |  |  |  |  |  |  |  |  |  |
|  | return | 7 | 3 | 36.1 ± 1.5 | 31.5 ± 0.7 | 3 | 0.39117 | ns | 3.6 ± 0.4 | 5.6 ± 0.8 | 2 | 0.291754 | ns |
|  | delay | 5 | 1 | 34.2 ± 1.5 | 29.8 |  |  | x | 4.2 ± 0.5 | 5.9 |  |  | x |
|  | stem | 9 | 5 | 31.8 ± 1.1 | 33.7 ± 0.8 | 15 | 0.59504 | ns | 4.1 ± 0.3 | 5.0 ± 0.3 | 9 | 0.29265 | ns |
|  | choice | 9 | 7 | 34.0 ± 1.0 | 32.5 ± 1.4 | 20 | 0.58195 | ns | 3.9 ± 0.2 | 4.5 ± 0.4 | 19 | 0.395062 | ns |
|  | reward | 9 | 5 | 33.4 ± 1.0 | 29.7 ± 1.4 | 7 | 0.14334 | ns | 3.8 ± 0.3 | 4.7 ± 0.4 | 5 | 0.37597 | ns |
|  | stem+choice | 8 | 7 | 32.5 ± 1.4 | 31.4 ± 1.5 | 25 | 0.75587 | ns | 4.0 ± 0.3 | 5.0 ± 0.6 | 11 | 0.395062 | ns |
| Suppl. Fig. 12G, H | CA3/DG high gamma |  |  |  |  |  |  |  |  |  |  |  |  |
|  | return | 1 | 0 | 57.1 |  |  |  | x | 1.4 |  |  |  | x |
|  | delay | 0 | 0 |  |  |  |  | x |  |  |  |  | x |
|  | stem | 5 | 2 | 58.4 ± 2.1 | 61.4 ± 3.1 | 3 | 0.96626 | ns | 2.0 ± 0.3 | 2.4 ± 0.6 | 4 | 0.857143 | ns |
|  | choice | 6 | 4 | 66.8 ± 5.9 | 60.2 ± 1.9 | 10 | 0.96626 | ns | 2.0 ± 0.3 | 2.9 ± 0.5 | 6 | 0.695476 | ns |
|  | reward | 4 | 2 | 63.2 ± 4.9 | 62.2 ± 12.1 | 3 | 0.96626 | ns | 1.8 ± 0.4 | 2.9 ± 0.3 | 1 | 0.695476 | ns |
|  | stem+choice | 4 | 2 | 73.6 ± 16.3 | 61.8 ± 4.9 | 4 | >0.999999 | ns | 1.7 ± 0.4 | 2.2 ± 0.5 | 2 | 0.782222 | ns |
| * mean ± SEM # adjusted for multiple comparisons by Holm-Sidak method |  |  |  |  |  |  |  |  |  |  |  |  |  |

\* mean ± SEM # adjusted for multiple comparisons by Holm-Šidák method

**Supplementary Figure 13.**
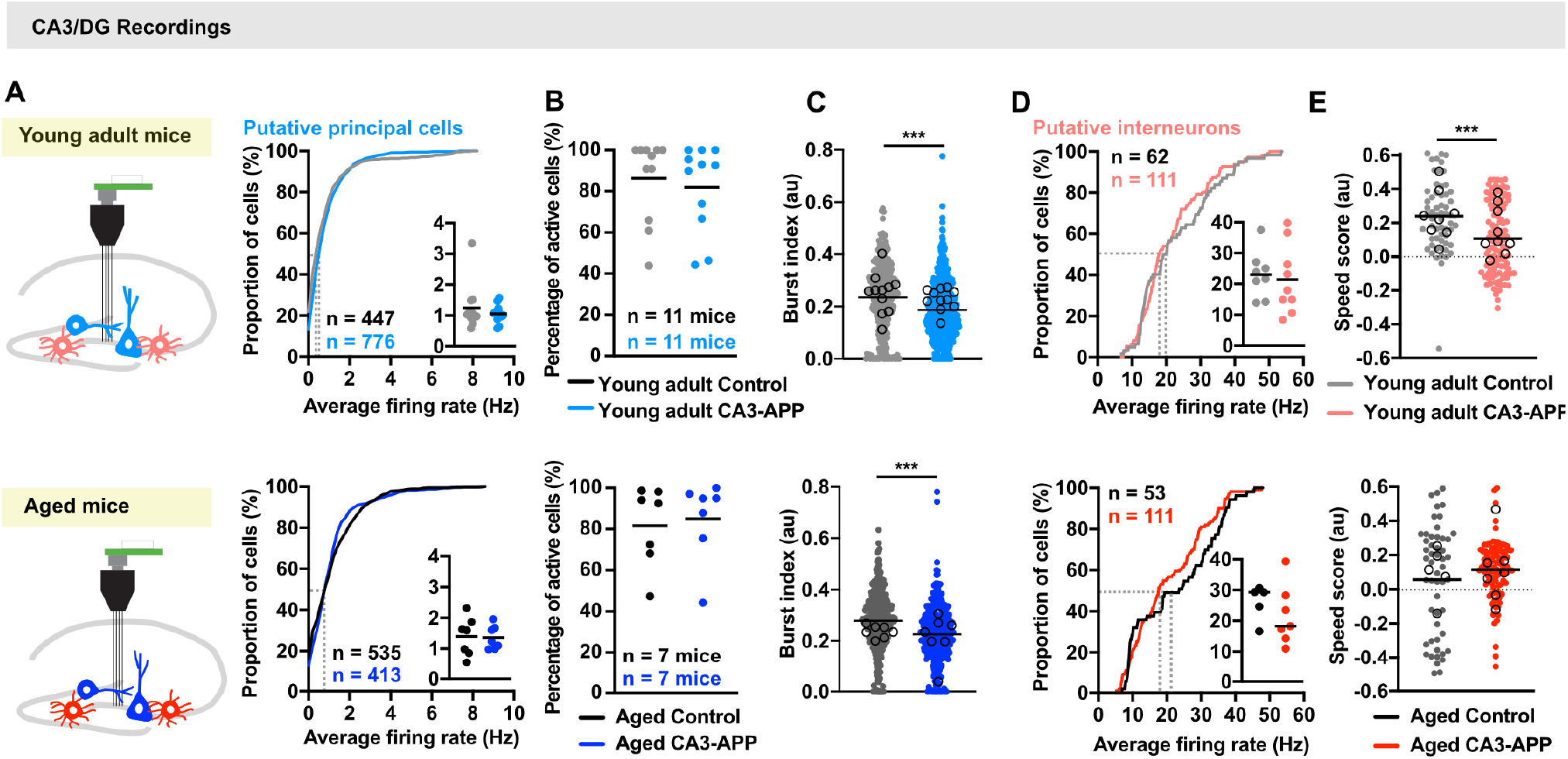
Firing rates of CA3/DG principal cells and interneurons are largely unchanged in CA3-APP mice compared to controls. **A.** Schematic of recordings at CA3/DG sites and cartoon of the recording device. Principal cells (dentate granule cells and CA3 pyramidal cells) are shown in blue and interneurons in red. Tetrode tracks often ended in the hilar region where DG and CA3c are intertwined. **B.** Cumulative density of average firing rates of all CA3/DG putative principal cells. Data include the entire recording session (r1, r2, d0, d2, d10). Inset shows the means of same data plotted per mouse (n = 11, young adult control; n = 11, CA3-APP; n = 7 aged control; n = 7, aged CA3-APP). Upper panel shows data for young adult mice and lower panel for aged mice. No differences were found between CA3-APP mice and age-matched controls (young adult: control 0.84 ± 0.06 Hz, CA3-APP 0.80 ± 0.03 Hz, p = 0.058; aged: control 1.2 ± 0.01 Hz, CA3-APP 1.1 ± 0.1 Hz, p = 0.96, MW test). **C.** Percentage of CA3/DG cells active during the figure-8 task did not differ between CA3-APP and age-matched control mice at both age points (young adult: control 86 ± 6%, CA3-APP 82 ± 16%, p = 0.56; aged: control 82 ± 7%, CA3-APP 85 ± 7%, p = 0.71, MW test). **D.** Burst index of principal cells. Horizontal lines depict the mean of all cells in a group, and open circles the mean per mouse. The burst index was reduced for principal cells of young adult CA3-APP mice compared to age-matched controls (young adult: control 0.24 ± 0.1, CA3-APP 0.20 ± 0.004, p <0.0001; aged: control 0.28 ± 0.01, CA3-APP 0.23 ± 0.01, p <0.0001). **E.** Cumulative density of average firing rates of all CA3/DG putative interneurons. Inset shows the means of the same data plotted per mouse. Average firing rates of CA3/DG interneurons did not differ between CA3-APP mice and age-matched controls (young adult: control 22.9 ± 1.4 Hz, CA3-APP 21.6 ± 0.9 Hz, p = 0.65; aged: control 22.6 ± 1.7 Hz, CA3-APP 20.5 ± 1.0 Hz, p = 0.37). **F.** Speed scores of interneurons, calculated using periods of mobility in the maze. The scores were reduced in young adult, but not in aged CA3-APP mice compared to age-matched controls. The lack of effect in aged mice can be explained by the already low scores in controls (young adult: control 0.20 ± 0.02, CA3-APP 0.11 ± 0.02, p = 0.0002; aged: control 0.03 ± 0.04, CA3-APP 0.1 ± 0.02, p = 0.60, MW test). See **Supplementary Table 15** for statistics. Figure corresponds to **Figure 2** and **Supplementary Figure 3** for CA1 data.

**Supplementary Table 15.** Data and statistics for Supplementary Figure 13.

| Fig. panel | Graph description | n (ctrl) | n (tg) | Norm Dist? | ctrl mean ± SEM | tg mean ± SEM | Statistical test | Stat. metrics | Df | P value | Sig. |
| --- | --- | --- | --- | --- | --- | --- | --- | --- | --- | --- | --- |
| Young adult mice |  |  |  |  |  |  |  |  |  |  |  |
| Suppl. Fig. 13B | Avg. firing rate PYR CA3/DG - all session | 447 | 777 | no/no | 0.84 ± 0.06 Hz | 0.80 ± 0.03 Hz | MW | U=15868 | | 0.0576 | ns |
|  | distribution all cells |  |  |  |  |  | KS | D=0.1002 |  | 0.0670 | ns |
| | pooled per mouse | 11 | 11 | no/yes | 1.2 ± 0.2 Hz | 1.1 ± 0.1 Hz | MW | U=60 | | >0.9999 | ns |
| Suppl. Fig. 13C | % active cells CA3/DG | 11 | 11 | no/no | 86 ± 6% | 82 ± 16% | MW | U=51.5 | | 0.5625 | ns |
| Suppl. Fig. 13D | Burst index PYR CA3/DG - behavior | 297 | 566 | no/no | 0.24 ± 0.01 | 0.20 ± 0.004 | MW | U=62638 | | <0.0001 | *** |
| | pooled per mouse | 11 | 11 | yes/yes | 0.24 ± 0.02 | 0.22 ± 0.01 | two sample t test | t=0.8363 | df=20 | 0.4129 | ns |
| Suppl. Fig. 13E | Avg. firing rate INT CA3/DG - all session | 62 | 111 | no/no | 22.9 ± 1.4 Hz | 21.6 ± 0.9 Hz | MW | U=3296 | | 0.6465 | ns |
|  | Distribution all cells |  |  |  |  |  | KS | D=0.1206 |  | 0.6092 | ns |
| | Pooled per mouse | 8 | 9 | yes/yes | 23.0 ± 2.7 Hz | 21.4 ± 3.7 Hz | two sample t test | t=0.7313 | df=15 | 0.7313 | ns |
| Suppl. Fig. 13F | INT CA3/DG speed modulation index | 59 | 111 | no/no | 0.20 ± 0.02 | 0.105 ± 0.02 | MW | U=2144 | | 0.0002 | *** |
|  | control different from 0? |  |  |  |  |  | Wilcoxon Signed Rank | W=1636 |  | <0.0001 | *** |
|  | CA3-APP different from 0? |  |  |  |  |  | Wilcoxon Signed Rank | W=3510 |  | <0.0001 | *** |
| | pooled per mouse | 8 | 9 | yes/yes | 0.24 ± 0.05 | 0.15 ± 0.05 | two sample t test | t=1.341 | df=15 | 0.1997 | ns |
| Aged mice |  |  |  |  |  |  |  |  |  |  |  |
| Suppl. Fig. 13B | Avg. firing rate PYR CA3/DG - all session | 535 | 413 | no/no | 1.2 ± 0.01 Hz | 1.1 ± 0.1 Hz | MW | U=110249 | | 0.9565 | ns |
|  | distribution all cells |  |  |  |  |  | KS | D=0.0866 |  | 0.0607 | ns |
| | pooled per mouse | 7 | 7 | yes/yes | 1.4 ± 0.2 Hz | 1.4 ± 0.1 Hz | two sample t test | t=0.1305 | df=12 | 0.8984 | ns |
| Suppl. Fig. 13C | % active cells CA3/DG | 7 | 7 | yes/no | 82 ± 7% | 85 ± 7% | MW | U=21 | | 0.7104 | ns |
| Suppl. Fig. 13D | Burst index PYR CA3/DG - behavior | 425 | 316 | no/no | 0.28 ± 0.01 | 0.23 ± 0.01 | MW | U=47452 | | <0.0001 | *** |
| | pooled per mouse | 7 | 7 | yes/yes | 0.24 ± 0.01 | 0.22 ± 0.03 | two sample t test | t=2.004 | df=12 | 0.0664 | ns |
| Suppl. Fig. 13E | Avg. firing rate INT CA3/DG - all session | 53 | 111 | no/no | 22.6 ± 1.7 Hz | 20.5 ± 1.0 Hz | MW | U=2685 | | 0.3692 | ns |
|  | Distribution all cells |  |  |  |  |  | KS | D=0.1882 |  | 0.1576 | ns |
| | Pooled per mouse | 5 | 7 | yes/yes | 26.1 ± 2.6 Hz | 21.7 ± 3.6 Hz | two sample t test | t=0.9011 | df=10 | 0.3887 | ns |
| Suppl. Fig. 13F | INT CA3/DG speed modulation index | 52 | 106 | no/no | 0.03 ± 0.04 | 0.1 ± 0.02 | MW | U=2615 | | 0.6042 | ns |
|  | control different from 0? |  |  |  |  |  | Wilcoxon Signed Rank | W=178 |  | 0.4234 | ns |
|  | CA3-APP different from 0? |  |  |  |  |  | Wilcoxon Signed Rank | W=3667 |  | <0.0001 | *** |
| | pooled per mouse | 5 | 7 | yes/yes | 0.10 ± 0.07 | 0.11 ± 0.07 | two sample t test | t=0.1384 | df=10 | 0.8927 | ns |

**Supplementary Figure 14.**
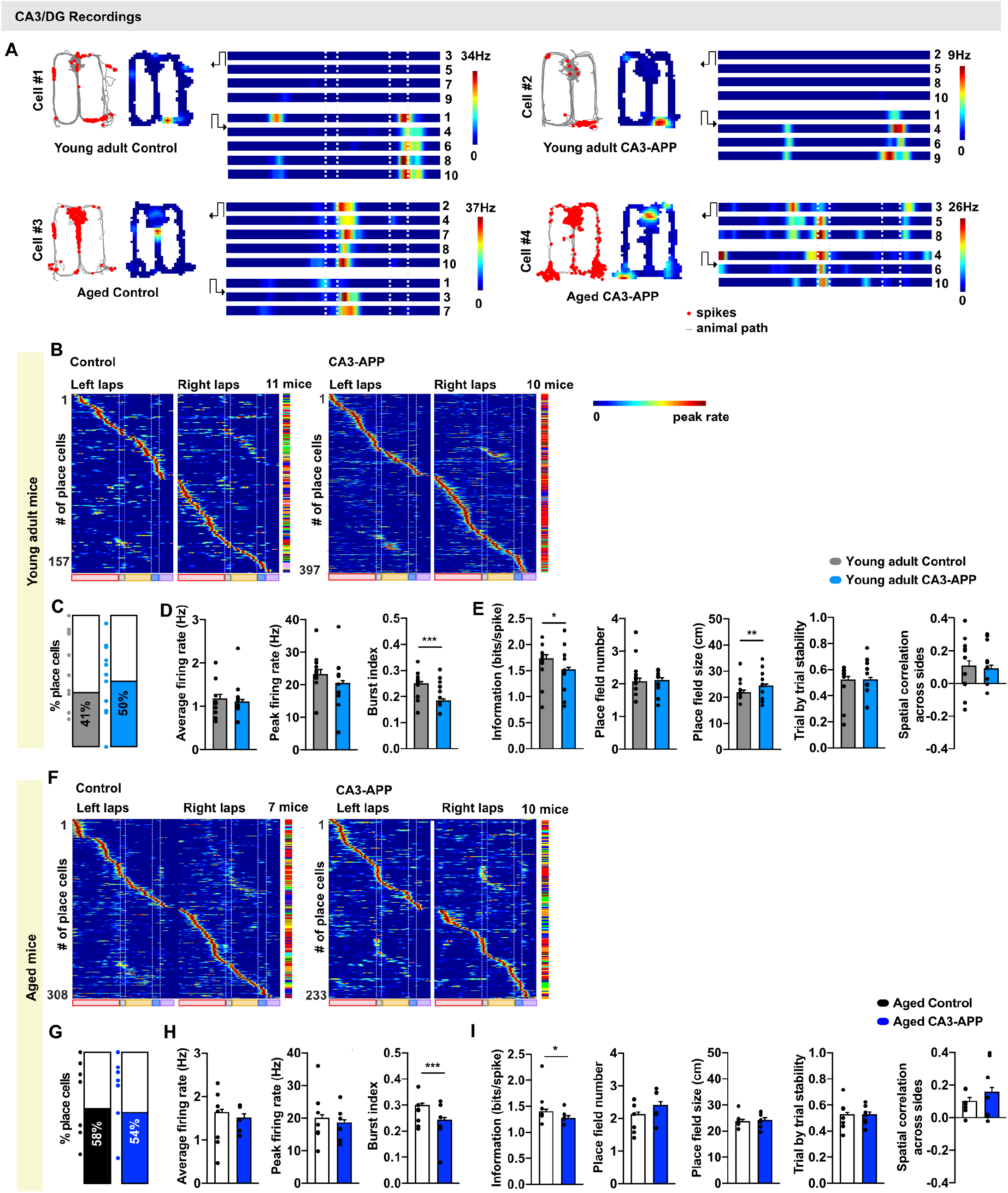
The spatial firing patterns of CA3/DG cells in CA3-APP mice remained accurate. **A.** One example place cell from each group is shown. Each panel includes the path (gray) with spike locations (red dots), a map of spike density (color-coded; 0 Hz, blue; maximum rate, red), linearized color-coded maps for each trial (0 Hz, blue; maximum rate, red; right-turn trials, top; left-turn trials, bottom). **B.** Average linearized rate maps for all CA3/DG place cells recorded from young adult control and from young adult CA3-APP mice. Left-turn and right-turn trials are shown separately. Cells are ordered by the location of the bin with the highest firing rate. In the vertical bar to the right of each panel, each individual mouse is assigned a color to show the distribution of cells across mice. **C.** Percent CA3/DG principal cells with place fields. Dots represented same data per mouse. No differences between young adult CA3-APP mice and age-matched controls were found (control 64 ± 9%, CA3-APP 52 ± 8 %, p = 0.31, two sample t-test). **D.** <u>Firing properties</u> of CA3/DG place cells. Black dots are the average per mouse. Bars and error bars are the mean and SEM of all cells. Average and peak firing rates of place cells did not differ between young adult CA3-APP mice and age-matched controls (control 23.3 ± 1.5 Hz, CA3-APP 20.6 ± 0.8 Hz, p = 0.46; MW test). The burst index of place cells was lower in young adult CA3-APP mice than in age-matched control mice (control 0.25 ± 0.01, CA3-APP 0.19 ± 0.01, p <0.0001, MW test). **E.** <u>Spatial properties</u> of CA3/DG place cells. Black dots are the average per mouse. Bars and error bars are the mean and SEM of all cells. Information content of place cells was slightly lower in young adult CA3-APP mice than in age-matched controls (control 1.7 ± 0.1, CA3-APP 1.5 ± 0.04, p = 0.013, MW test). Place field number per cell did not differ (control 2.1 ± 0.1, CA3-APP 2.1 ± 1.2, p = 0.69, MW test), but place field size showed a small increase in young adult CA3-APP mice compared to age-matched controls (control 22.0 ± 0.8, CA3-APP 25.5 ± 0.6, p = 0.0034, MW test). No differences in stability across trials or in correlation coefficients across left and right maze arms (indicating that spatial firing patterns were distinct) were found for CA3/DG place cells of young adult CA3-APP mice compared to age-matched controls (p = 0.77, MW test). **F.** Average linearized rate maps for all CA3/DG place cells recorded from aged control and from aged CA3-APP mice. Data are displayed as in B. **G.** No differences in % place cells were found between aged CA3-APP mice and age-matched controls (control 66 ± 11%, CA3-APP 72 ± 10%, p = 0.7003, two sample T-test). Data are displayed as in C. **H.** The average and peak firing rates (control 20.1 Hz ± 0.9, CA3-APP 18.7 ± 0.8 Hz, p = 0.70, MW test) of place cells did not differ between aged CA3-APP mice and age-matched controls, but the burst index of place cells in aged CA3-APP mice was lower than in age-matched control mice (control 0.30 ± 0.01, CA3-APP 0.24 ± 0.03, p <0.0001, MW test). Data are displayed as in D. **I.** Information content of place cells was slightly reduced in aged CA3-APP mice compared to age-matched controls (control 1.4 ± 0.03, CA3-APP 1.3 ± 0.04, p = 0.019, MW test). No other differences in spatial properties of CA3/DG place cells from aged CA3-APP compared to age-matched control mice were found [number of place fields (control 2.1 ± 0.1, CA3-APP 2.4 ± 0.1, p = 0.013), place field size (control 23.9 ± 0.6, CA3-APP 24.3 ± 0.3, p = 0.75, MW test), stability across trials (p = 0.98, MW test)]. Data are displayed as in E. See **Supplementary Table 16** for statistics. Figure corresponds to **Figure 3** and **Supplementary Figure 4** for CA1 data.

**Supplementary Figure 15.**
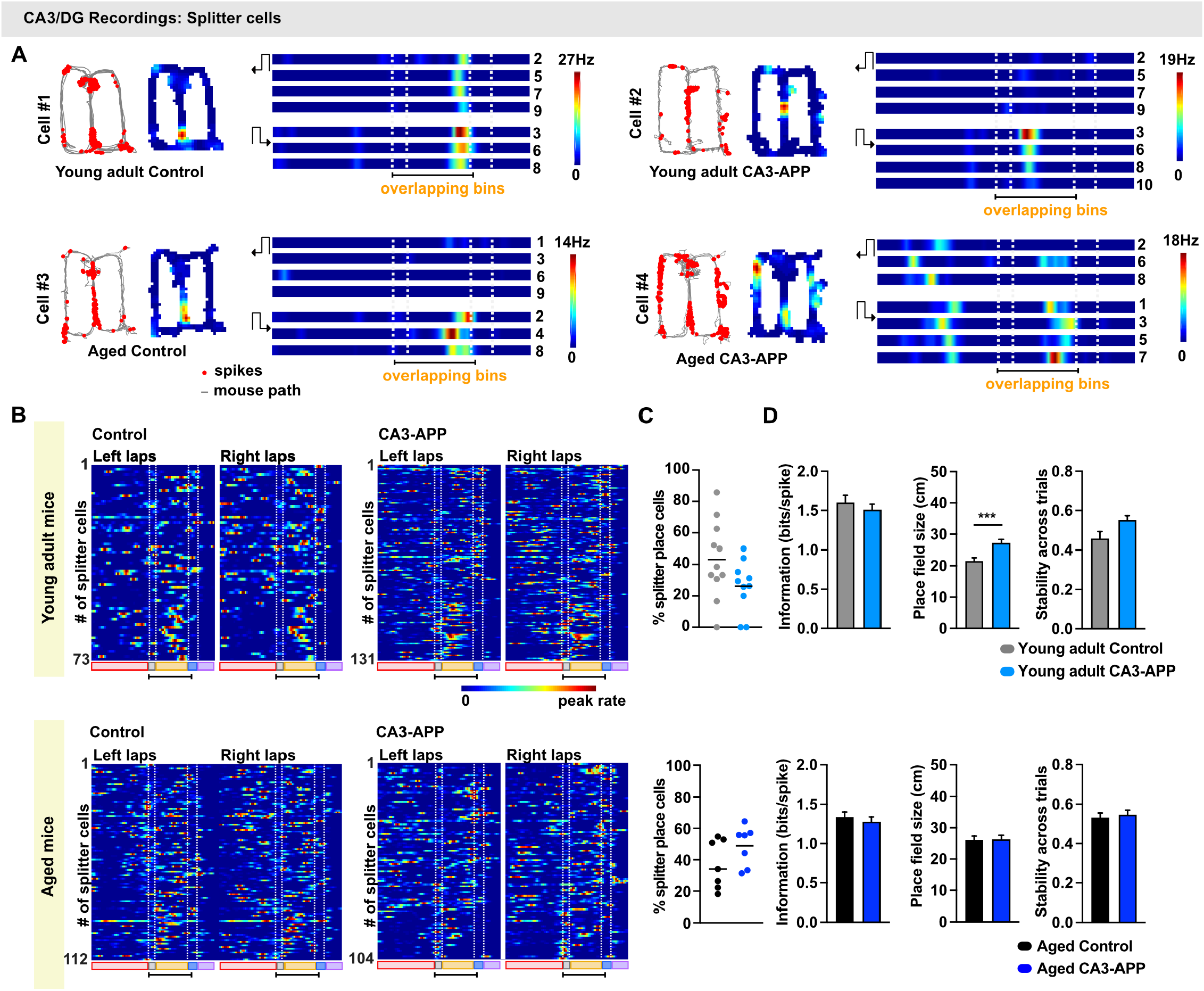
Fraction and properties of CA3/DG splitter place cells were largely maintained. **A.** Example cells that were selectively active on the center arm in either only right-turn or left-turn laps (‘splitter cells’). One example cell from each group is shown. Each panel includes the path (gray) with spike locations (red dots), a map of spike density (color-coded; 0 Hz, blue; maximum rate, red), linearized color-coded maps for each trial (0 Hz, blue; maximum rate, red; right-turn trials, top; left-turn trials, bottom). **B.** Color-coded firing rates of all splitter cells in each experimental group. Cells are ordered by the position of their peak firing field along the lap. Spatial bins that overlap between left-turn and right-turn trials are indicated with a black line at the bottom of each panel. Cells are ordered by their degree of selectivity for either correct right-turn or left-turn trials (n = 73 splitter cells for young adult control and n = 131 for CA3-APP mice; n = 112 splitter cells for aged control and n = 104 for CA3-APP mice). **C.** The percentage of place cells that reached the criteria for splitter cells in the figure-8 maze were calculated per mouse. The percentage was not statistically different between genotypes in both the young adult mice and aged mice (young adult: control 39 ± 7%, n= 11 mice, CA3-APP 23 ± 5%, n = 11 mice, p = 0.10; aged: control 33 ± 7%, n = 7 mice, CA3-APP 43 ± 5%, n= 7 mice, p = 0.32, two sample t-test). **D.** Place field size is transiently increased in young adult CA3-APP place cells (control 21.9 ± 1.1cm, CA3-APP 27.6 ± 1.2 cm, p = 0.001), but not in place cells from aged CA3-APP (control 26.0 ± 1.4 cm, CA3-APP 27.5 ± 1.4 cm, p = 0.24, MW test). The information score (young adult: control 1.6 ± 0.1, CA3-APP 1.5 ± 0.1, p = 0.34; aged: control 1.3 ± 0.1, CA3-APP 1.3 ± 0.1, p = 0.55, MW test) and trial-by-trial stability of CA3/DG splitter cells did not differ between CA3-APP mice and age-matched control mice (p = 0.056 for young adult and p = 0.74 for aged, MW test). All bar graphs show the mean ± SEM over all cells in a group. See **Supplementary Table 16** for statistics. Figure corresponds to **Supplementary Figure 4** for CA1 data.

**Supplementary Table 16.**
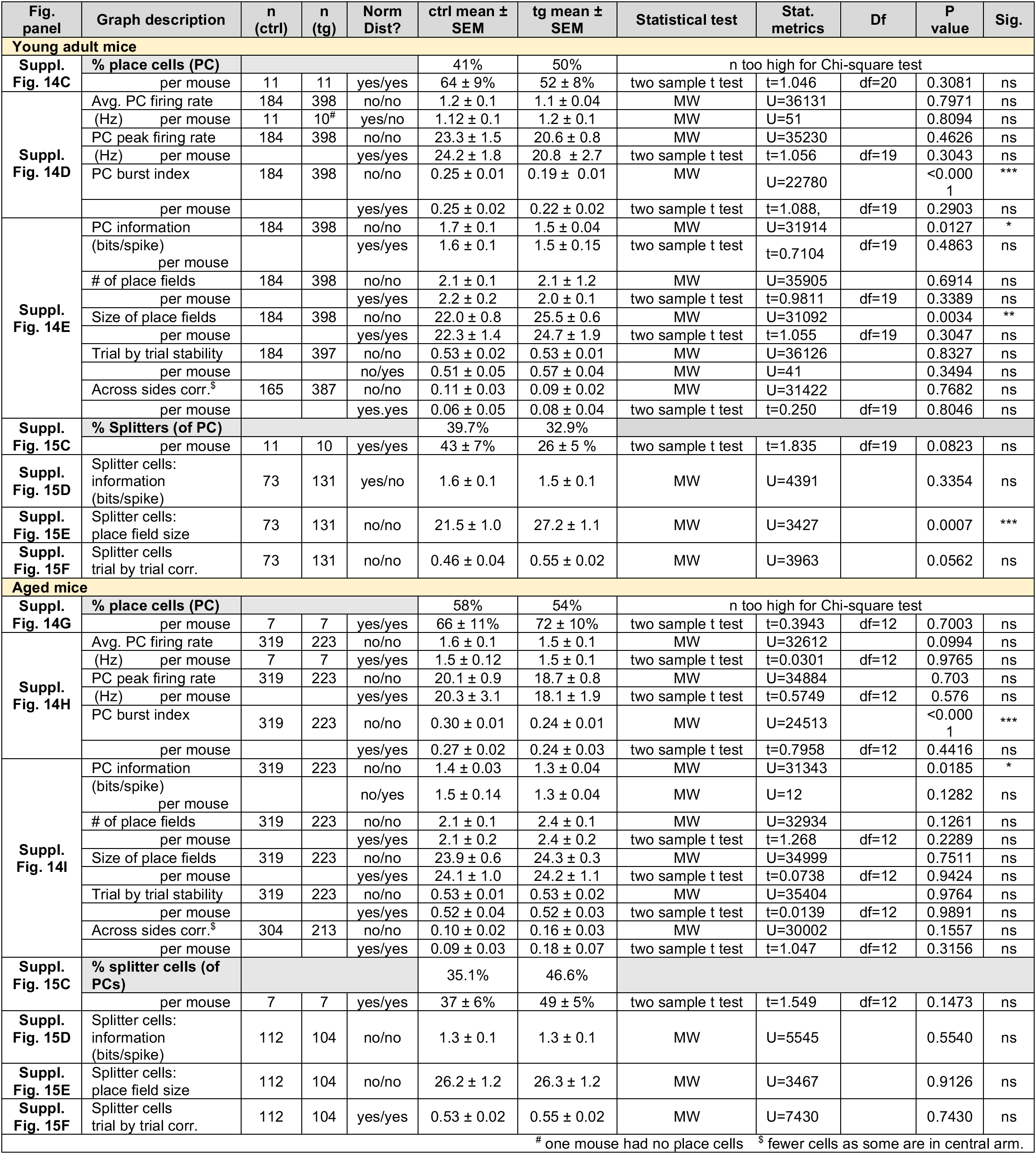
CA3/DG place cell properties, related to Supplementary Figures 14 and 15.

**Supplementary Figure 16.**
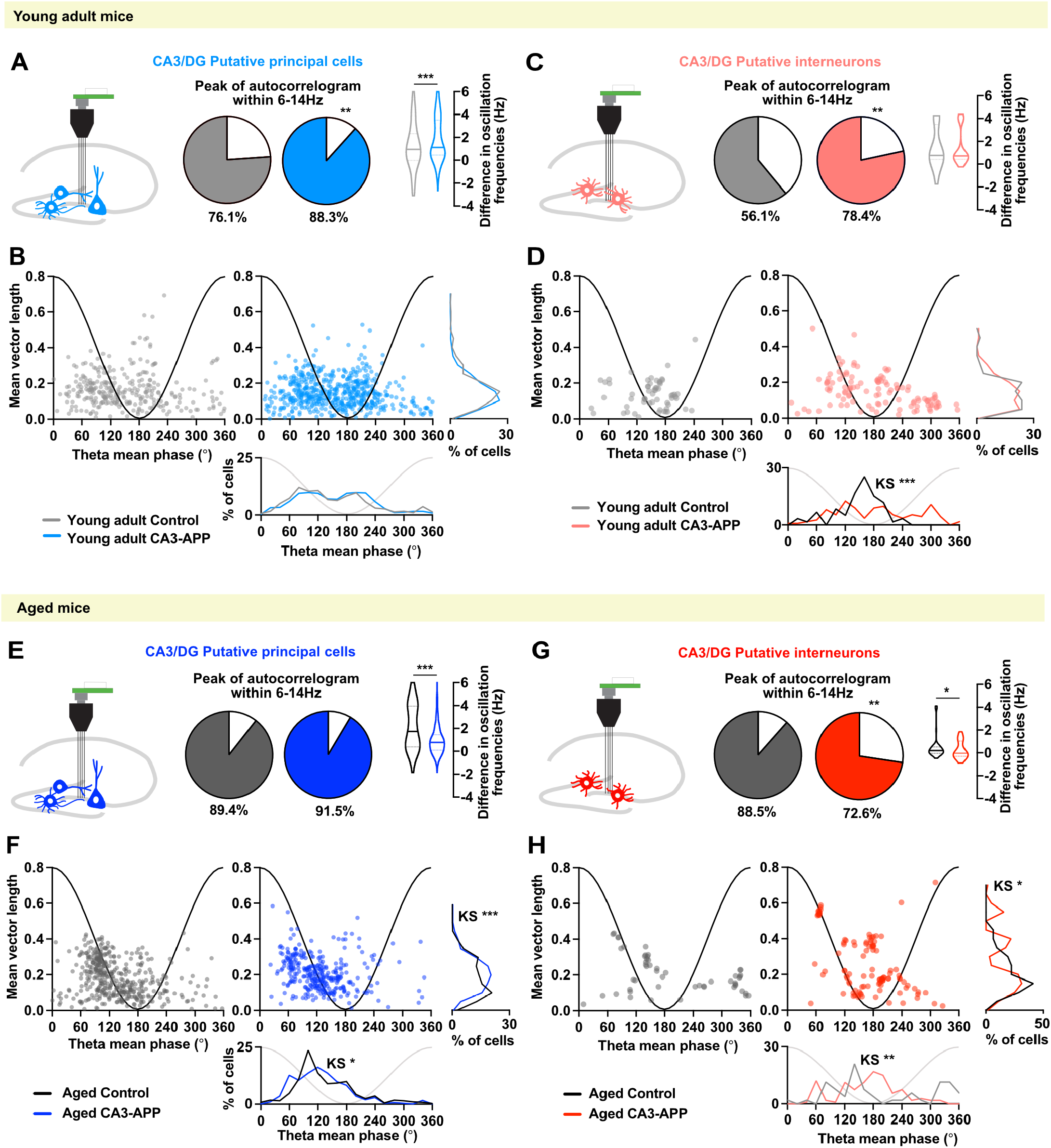
Temporal firing patterns of CA3/DG principal cells and interneurons. **A.** Spike oscillation frequencies in the theta range were obtained by performing the spike-time autocorrelation for each neuron and by using a fast-Fourier transform to obtain the predominant frequency (peak of autocorrelogram). The fraction of cells with detectable peaks in the theta range (6-14Hz) is reported as the colored portion of the pie charts. A higher proportion of CA3/DG principal cells from CA3-APP mice than from age-matched control mice showed oscillations in the theta range (p < 0.0001, Chi-square test). For cells oscillating in the theta range, we then calculated the differences between the predominant spike oscillation frequency and the predominant LFP oscillation frequency in the same recording session. The difference between spike and LFP oscillation frequencies was larger in young adult CA3-APP mice compared to age-matched controls (control 1.2 ± 0.1 Hz, CA3-APP 1.9 ± 0.1 Hz, p = 0.0009, MW test). **B.** The preferred LFP phase when cells spiked (mean phase) is plotted against the amplitude of the theta phase locking (mean vector length) for CA3/DG principal cells from control (grey) and CA3-APP mice (blue). No differences were found in the phase preference (control 150.8 ± 4.7°, CA3-APP 155.5 ± 3.2°, p = 0.15, MW test) or in phase locking of CA3/DG principal cells in young adult CA3-APP mice compared to control littermates (control 0.17 ± 0.01, CA3-APP 0.15 ± 0.003, p = 0.18, MW test). **C.** Similar to principal cells, the fraction of CA3/DG interneurons with spiking frequency in the theta range was higher in young adult CA3-APP mice compared to control littermates (p = 0.016, Chi-square test). The difference between the spike oscillation frequency of interneurons and the LFP oscillation frequency did not change between young adult CA3-APP mice and age-matched control mice (control 1.4 ± 0.3 Hz, CA3-APP 1.4 ± 0.2 Hz, p = 0.64, MW test). Data are displayed as in A. **D.** The distribution of theta phase preference shifted towards later values in the cycle for CA3/DG interneurons in young adult CA3-APP mice compared to age-matched controls (control 148.0 ± 6.4°, CA3-APP 185.0 ± 7.9°, p < 0.0001, MW test). Phase locking of spikes did not differ (control 0.14 ± 0.01, CA3-APP 0.15 ± 0.01, p = 0.68, MW test). Data are displayed as in B. **E.** The fraction of cells with spiking frequencies in the theta range did not differ between aged CA3-APP mice and age-matched control mice (p = 0.3529, Chi-square test). The difference between principal cell spike oscillation frequency and LFP oscillation frequency was substantially reduced in aged CA3-APP mice compared to aged control mice (control 2.2 ± 0.1 Hz, CA3-APP 0.9 ± 0.1 Hz, p < 0.0001, MW test). Data are displayed as in A. **F.** While the mean phase preference of CA3/DG principal cells did not differ in aged CA3-APP mice compared to littermate controls (control 135.5 ± 3.0°, CA3-APP 126.8 ± 3.2°, p = 0.028, MW test), the phase locking of spikes to theta oscillations was increased in CA3-APP mice (control 0.183 ± 0.005, CA3-APP 0.208 ± 0.005, p = 0.0002, MW test). Data are displayed as in B. **G.** The fraction of CA3/DG interneurons with spiking frequency in the theta range was lower in aged CA3-APP mice compared to control littermates (p = 0.024, Chi-square test). The difference between the spike oscillation frequency of CA1 interneurons and the LFP oscillation frequency was decreased in aged CA3-APP mice compared to age-matched control mice (control 0.6 ± 0.2 Hz, CA3-APP 0.3 ± 0.1 Hz, p = 0.028, MW test), with most interneurons now oscillating at frequencies close to the lower LFP frequency of CA3-APP mice. Data are displayed as in C. **H.** The distribution of theta phase preference for CA3/DG interneurons spikes from control mice shows a clear peak at the trough of theta oscillations. This is completely absent for CA3/DG interneurons of CA3-APP mice, such that the two distributions differed (p = 0.0057, KS test) even though the medians of the two distributions were not significantly different (control 198.5 ±14.3°, CA3-APP 168.4 ± 6.0°, p = 0.33, MW test). Phase locking of CA3/DG interneurons to theta oscillations did not differ between CA3-APP mice and age-matched controls (control 0.20 ± 0.01, CA3-APP 0.26 ± 0.02, p = 0.15, MW test). Data are displayed as in D. For statistical analysis see **Supplementary Table 17**. Figure corresponds to **Supplementary Figure 8** for CA1 data.

**Supplementary Table 17.**
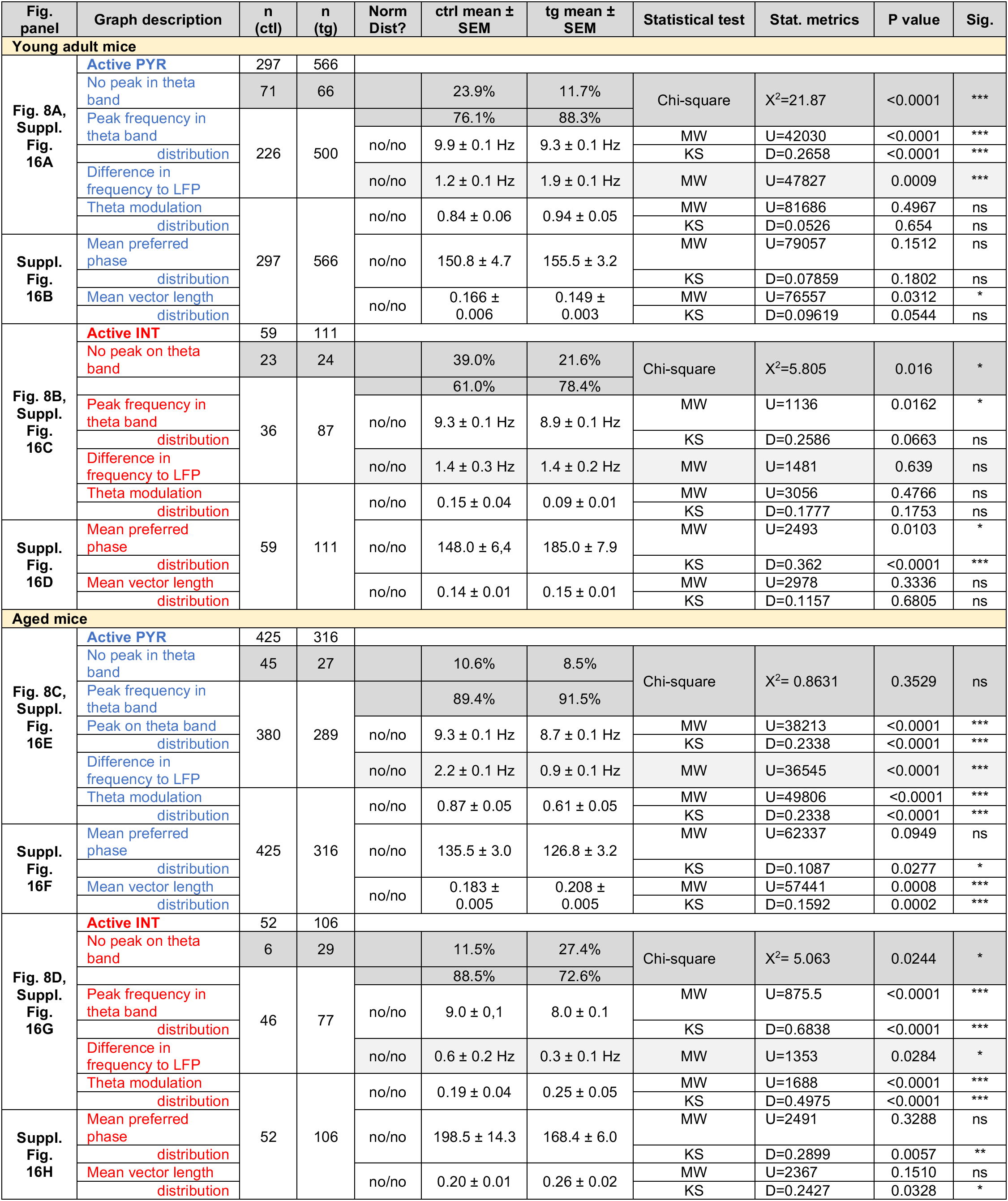
Theta modulation of CA3/DG single units, related to Figure 8 and Supplementary Figure 16.

## Notes

### Competing Interest Statement

The authors have declared no competing interest.

